# TXN, a Xanthohumol Derivative, Attenuates High-Fat Diet Induced Hepatic Steatosis by Antagonizing PPARγ

**DOI:** 10.1101/2021.01.11.426043

**Authors:** Yang Zhang, Gerd Bobe, Cristobal L. Miranda, Malcolm B. Lowry, Victor L. Hsu, Christiane V. Löhr, Carmen P. Wong, Donald B. Jump, Matthew M. Robinson, Thomas J. Sharpton, Claudia S. Maier, Jan F. Stevens, Adrian F. Gombart

## Abstract

We previously reported xanthohumol (XN), and its synthetic derivative tetrahydro-XN (TXN) attenuates high-fat diet (HFD) induced obesity and metabolic syndrome in C57BL/6J mice. The objective of the current study was to determine the effect of XN and TXN on lipid accumulation in the liver. Non-supplemented mice were unable to adapt their caloric intake to 60% HFD, resulting in obesity and hepatic steatosis; however, TXN reduced weight gain and decreased hepatic steatosis. Liver transcriptomics indicated TXN might antagonize lipogenic PPARγ actions *in vivo*. XN and TXN inhibited rosiglitazone-induced 3T3-L1 cell differentiation concomitant with decreased expression of lipogenesis-related genes. A PPARγ competitive binding assay showed XN and TXN bind to PPARγ with an IC_50_ similar to pioglitazone and 8-10 times stronger than oleate. Molecular docking simulations demonstrated XN and TXN bind in the PPARγ ligand-binding domain pocket. Our findings are consistent with XN and TXN acting as antagonists of PPARγ.

## Introduction

Non-alcoholic fatty liver disease (NAFLD) is a major global health threat characterized by excessive hepatic lipid droplet accumulation with a history of little or no alcohol consumption (Hashimoto, Taniai and Tokushige, 2013). About one-quarter of the US population suffers from NAFLD (Estes *et al*., 2018) with rates in the rest of the world ranging from 14% in Africa to 32% in the Middle East (Younossi *et al*., 2016). The continuing obesity and diabetes epidemic drives increasing rates of NAFLD (Estes *et al*., 2018). Unfortunately, no FDA-approved drugs exist for its treatment. Sustained healthy life style changes and weight loss are the only interventions proven effective in preventing the onset and progression of NAFLD (Stefan, Häring and Cusi, 2019). Thus, there is a critical need for novel and effective interventions.

As a central hub for lipid metabolism, a healthy liver maintains homeostasis among uptake, esterification, oxidation and secretion of fatty acids (FAs) (Goldberg and Ginsberg, 2006). Overconsumption of saturated FAs or sugars can overload the liver, disrupt lipid homeostasis, resulting in excess storage of triacylglycerols (TAG) in hepatocytes and the onset and progression of hepatic steatosis (Ipsen, Lykkesfeldt and Tveden-Nyborg, 2018). Given that PPARγ is important in hepatic lipogenesis (Sharma and Staels, 2007), it has attracted considerable attention as a therapeutic target for NAFLD (Almeda-Valdés, Cuevas-Ramos and Aguilar-Salinas, 2009).

Attenuated PPARγ activity in heterozygous PPARγ-deficient (*PPARγ*^+/-^) C57BL/6J mice protects against HFD-induced obesity, liver steatosis and adipocyte hypertrophy; however, treatment with the PPARγ agonist pioglitazone (PGZ) abrogates the protection against hypertrophy and decreases insulin sensitivity (Kubota *et al*., 1999) suggesting a potential beneficial use for PPARγ antagonists to treat hepatic steatosis. PPARγ antagonists tanshinone IIA (Gong *et al*., 2009), β-cryptoxanthine (Goto *et al*., 2013), protopanaxatriol (Zhang *et al*., 2014), isorhamnetin (Zhang *et al*., 2016), and Gleevec (Choi *et al*., 2016) improved multiple metabolic parameters in diet-induced obese (DIO) mice. These observations strongly suggest that moderate inhibition of PPARγ activity may reduce the risk for developing hepatic steatosis induced by diet, and PPARγ antagonists may be useful for the treatment and prevention of NAFLD.

Xanthohumol (XN), a prenylated flavonoid found in hops (*Humulus lupulus* L.), improves multiple parameters of MetS in rat and mouse models (Legette *et al*., 2014; Miranda *et al*., 2016, 2018). Tetrahydroxanthohumol (TXN), a non-estrogenic synthetic XN derivative (**Fig. 1**), appears more effective in ameliorating MetS in DIO mice than XN possibly due to its 5-, 10-, and 12-fold higher levels in the muscle, plasma and liver, respectively, as compared with XN (Miranda *et al*., 2018). Both compounds likely mediate their benefits via multiple mechanisms. XN inhibits differentiation of preadipocytes and induces apoptosis in mature adipocytes (Yang *et al*., 2007; Rayalam *et al*., 2009), attenuates the function of SREBP-1 by repressing its maturation (Miyata *et al*., 2015) and induces beiging of white adipose tissue, decreases adipogenesis, and induces lipolysis (Samuels, Shashidharamurthy and Rayalam, 2018). We recently showed that XN and TXN significantly change gut microbiota diversity and abundance, alter bile acid metabolism and reduce inflammation in mice fed a HFD (Zhang *et al*., 2020). Collectively, these data suggest both XN and TXN are effective for treatment of metabolic disorders and are promising candidates for NAFLD prevention and treatment.

**Figure 1.**
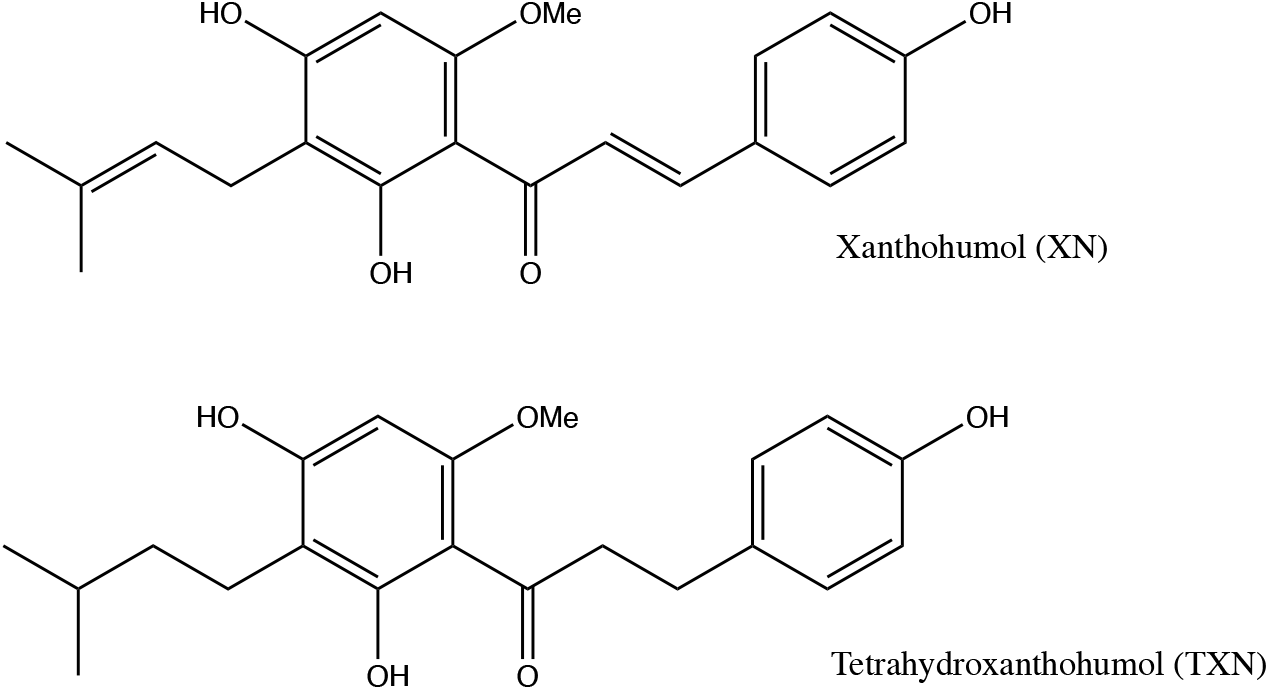
Structures of XN and its synthetic derivative TXN.

In the present study, we show a daily oral intake of TXN at 30 mg/kg body weight (BW) or XN at a daily dose of 60 mg/kg BW strongly suppresses diet-induced liver steatosis in C57BL/6J male mice. Supervised machine learning of liver RNA-seq data identified perturbations in PPARγ signaling. Based on cell culture experiments, a PPARγ competitive binding assay and molecular docking studies, we provide evidence that XN and TXN act as novel PPARγ antagonists with moderate binding activity. Collectively, our findings suggest appropriate functional antagonism of PPARγ is a logical approach to prevent and treat diet-induced liver steatosis and other related metabolic disorders. The structures of XN and TXN could serve as scaffolds for synthesis of more effective compounds to treat NAFLD.

## Results

### 1.TXN attenuates HFD-induced weight gain independent of caloric intake

As expected, C57BL/6J mice on a 60% HFD (**Fig. 2A**, solid blue line) gained more BW than mice on the LFD (**Fig. 2A**, dotted black line) throughout the experimental period (week 1: *p* < 0.05; week 2-16: *p* < 0.001; repeated measures). TXN-supplementation (**Fig. 2A**, solid dark green line) attenuated HFD-induced BW gain throughout the experimental period (week 1: *p* < 0.05; week 2-16: *p* < 0.001; repeated measures). XN supplementation showed a dose response effect: the higher dosage (HXN; **Fig. 2A**, solid red line), but not the lower dosage (LXN; **Fig. 2A**, solid yellow line), attenuated HFD-induced BW gain between week 8 and 16. When BW gain was expressed as % of initial BW, HFD-fed mice almost doubled their initial BW (+98.3 ± 2.7%), whereas TXN-treated mice gained 33% less (+66.2 ± 5.8%, *p* < 0.0001), and LFD-fed mice gained 53% less (+45.8 ± 4.3%, *p* < 0.0001) than HFD-fed mice (**Fig. 2B**). Although not statistically significant, both LXN- and HXN-treated mice gained 7.5% and 11% less, respectively (90.0 ± 3.3%, *p* = 0.20; 87.6 ± 3.9%, *p* = 0.07; **Fig. 2B**). In male C57BL/6J mice, a BW of approximately 40 g is a critical tipping point from which metabolic dysfunction occurs (van Beek *et al*., 2015). After 16 weeks, mean BW for these mice was LFD (37.5 ± 1.1g), HFD (50.3 ± 0.6g), LXN (49.9 ± 1.1g), HXN (47.4 ± 1.1g) and TXN (42.2 ± 1.6g).

**Figure 2.**
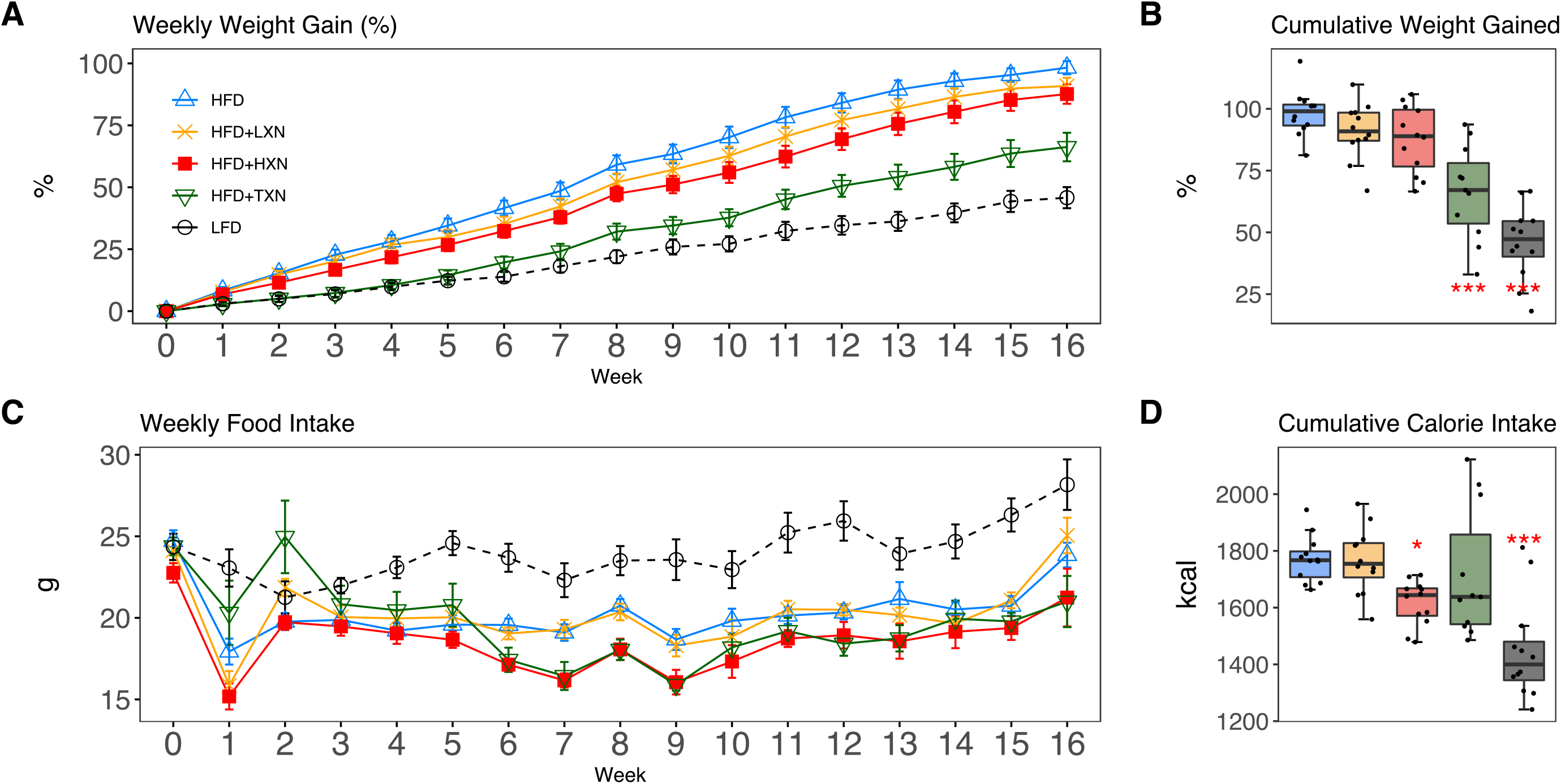
TXN and HXN suppress HFD-induced BW gain independent of caloric intake. Mice were fed either a LFD (black dashed line with empty circles, n = 12), a HFD (blue solid line with empty triangles, n = 12), HFD+LXN (yellow solid line with crosses, n = 12), HFD+HXN (red solid line with squares, n = 12), or HFD+TXN (green solid line with empty triangles, n = 11) for 16 weeks. (A) BW gain was assessed once per week. Data is expressed as means ± SEM. Repeated measurement of ANOVA was used to calculate *p*-values for the percentage of weight gained weekly. (B) Total percent BW gained at the end of the16-week feeding period. Data is expressed as quartiles. (C) Food intake was assessed once per week during the 16-week feeding period. Data is expressed as means ± SEM. Repeated measurement of ANOVA was used to calculate *p*-values for weekly food intake. (D) Total calories consumed at the end of 16-week feeding period. Data are expressed as quartiles. Source files of data used for the analysis and visualization are available in the Figure 2—source data 1.

Overtime, mice adapted to the HFD by consuming less food than LFD-fed mice (**Fig. 2C**). However, the discrepancy in food consumption was insufficient to counteract the elevated caloric intake (**Fig. 2D**). HXN-treated mice adapted better to the HFD, indicated by decreased food intake at week 1, 6-10, 13, and 16 (*p* < 0.05), and caloric intake (*p* = 0.01) (**Fig. 2C-D**), resulting in less BW gain. In contrast, the attenuated BW gain in TXN-treated mice was not accompanied by a significant reduction in food or caloric intake (**Fig. 2C-D**).

### 2. TXN attenuates hepatic steatosis and HFD-induced obesity

HFD-induced BW gain was primarily body fat accumulation, as indicated by measurements obtained from DEXA scans. HFD mice had greater fat mass than LFD mice (*p* < 0.0001; **Fig. 3A**). Linear regression of total fat mass to total caloric intake revealed a strong relationship between caloric intake and fat mass among groups (r = +0.52; *p* < 0.0001) and within LFD-fed mice (r = +0.79; *p* = 0.002; **Fig. 3A1**). In contrast, caloric intake was not correlated to fat mass in any HFD group (**Fig. 3A2-5**), indicating a disconnection between caloric intake and fat mass after prolonged HFD consumption. Supplementation with HXN (−9.93%; *p* < 0.05) and even more so with TXN (−27.7%; *p* < 0.001) decreased body fat mass on HFD (**Fig. 3A**), indicating that HXN and TXN attenuated the HFD-induced body fat accumulation and that this effect was not explained by changes to caloric intake (**Fig. 3A4-5**).

**Figure 3.**
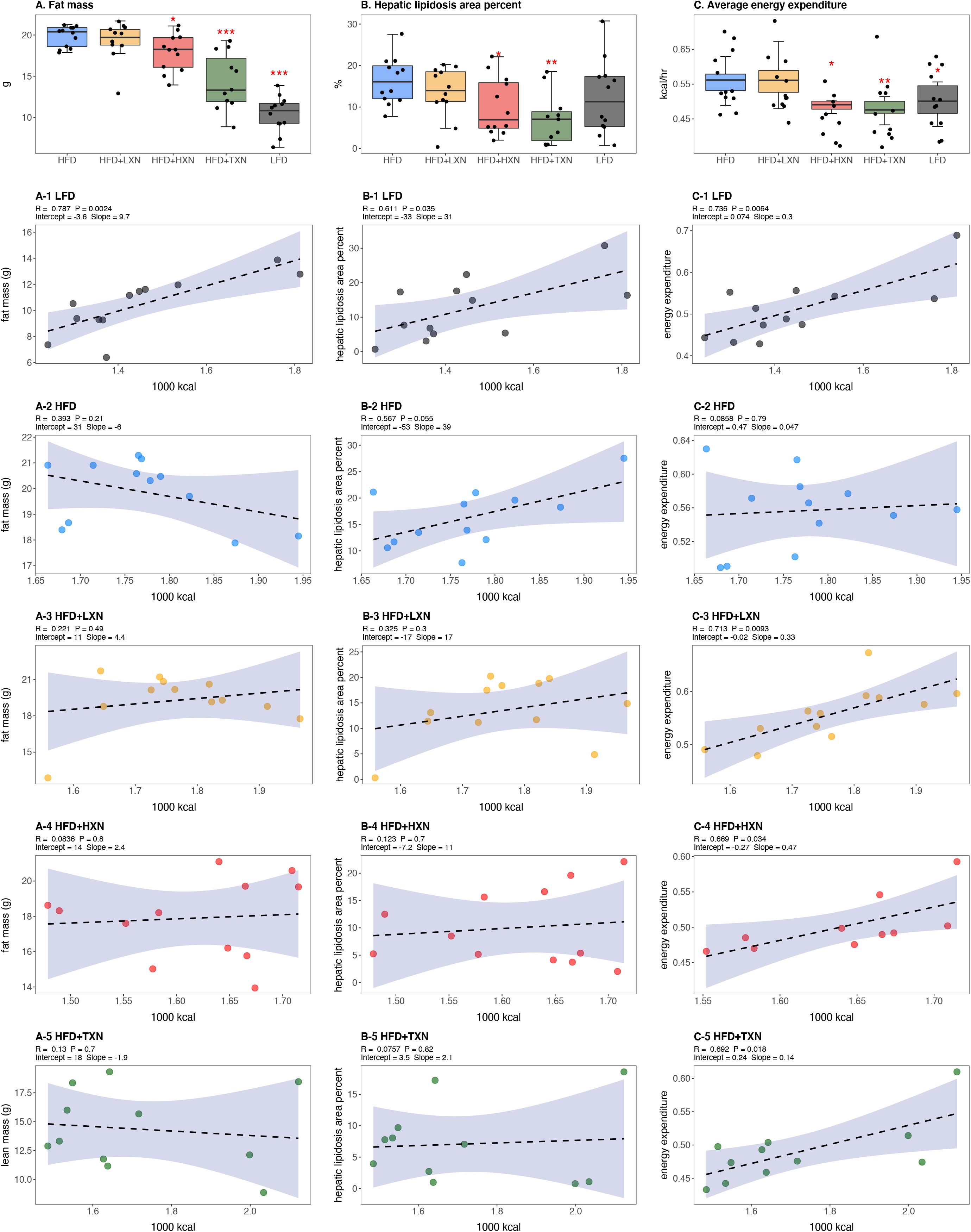
Energy homeostasis imbalance induced by HFD is prevented by XN and TXN supplementation. Mice were fed either a LFD (black, n = 12), a HFD (blue, n = 12), HFD+LXN (yellow, n = 12), HFD+HXN (red, n = 12), or HFD+TXN (green, n = 11) for 16 weeks. (A) Total fat mass measured by DXA scan two days prior to necropsy is expressed as quartiles. (A-1) Relationship between total fat mass and total caloric intake over 16 weeks of feeding for LFD; (A-2) HFD; (A-3) HFD+LXN; (A-4) HFD+HXN; and (A-5) HFD+TXN groups. (B) Hepatic lipidosis area percent expressed as quartiles. (B-1) Relationship between hepatic lipidosis area percent and total caloric intake over 16 weeks of feeding for LFD; (B-2) HFD; (B-3) HFD+LXN; (B-4) HFD+HXN; and (B-5) HFD+TXN groups. (C) Average energy expenditure over two light-dark cycles (48 hours) obtained using metabolic cages and expressed as quartiles. (C-1) Relationship between energy expenditure and total caloric intake over 16 weeks of feeding for LFD; (C-2) HFD; (C-3) HFD+LXN; (C-4) HFD+HXN (with removal of two outliers); (C-5) for HFD+TXN groups. Pre-planned general linear model with contrasts were used to calculate *p*-values in A, B and C. \**p* < 0.05, \*\**p* < 0.01, \*\*\**p* < 0.001. Linear regression analyses of total calories versus total fat mass (A1-5), hepatic lipidosis area percent (B1-5), and average energy expenditure (C1-5) in mice were done using stats package version 3.6.2 in R. Blue shading represents 95% CI of the regression line. Absolute value of R, p-value, intercept, and slope for the regression are reported above each corresponding panel. Source files of data used for the analysis are available in the Figure 3—source data 1.

Hepatic steatosis was measured by percent surface area occupied by lipid vacuoles in formalin-fixed, paraffin-embedded liver by image analysis of photomicrographs. In the absence of supplementation, HFD- and LFD-fed mice shared similar hepatic lipid areas (**Fig. 3B**). Caloric intake was positively correlated with hepatic lipid area on both LFD-fed mice (r = +0.61, *p* = 0.03; **Fig. 3B1**) and a HFD (r = +0.57, *p* = 0.05; **Fig. 3B2**). Supplementation with HXN (*p* < 0.05) and TXN (*p* < 0.01) mitigated hepatic steatosis, independent of caloric intake (**Fig. 3B4-5**).

Changes in energy balance may drive changes in obesity-related steatosis. We investigated TXN on whole-body energy metabolism to determine mechanisms of TXN protection from weight gain, which can influence steatosis. Towards the end of the study, we measured whole-body expenditure for all 59 mice using a computer-controlled indirect calorimetry system (metabolic cages). Energy expenditure was calculated from the oxygen and carbon dioxide exchange ratio using the Weir equation (Weir, 1949). Total energy expenditure contains energy expenditure for basal metabolism, body tissue synthesis, digestion, and physical activity (Speakman, 2013). Mice consuming HFD and mice supplemented with LXN had higher (*p* < 0.05) energy expenditure than mice on LFD, HXN and TXN (**Fig. 3C**). Caloric intake was positively correlated with energy expenditure in LFD- (**Fig. 3C1**), LXN- (**Fig. 3C3**), HXN- (**Fig. 3C4**) and TXN-fed mice (**Fig. 3C5**) but was not correlated with energy expenditure in HFD mice (**Fig. 2C2**). We investigated the influence of body mass on energy expenditure using analysis of covariance (ANCOVA) of body mass upon entry into the cages between diets (Tschöp *et al*., 2011). ANCOVA revealed that LXN, HXN, or TXN supplementation did not change the positive relationship between energy expenditure and body mass (**Figure 3—figure supplement 1**).

As a marker of hepatic lipid uptake and export, fasting plasma TAG level was measured at the end of the study. Similar to hepatic lipid area, fasting plasma TAG did not reflect the caloric density of the diet (**Fig. 3—figure supplement 2**); namely, there was an inverse relationship between caloric intake and plasma TAG among LFD mice (Spearman, r = -0.60, *p* = 0.04; **Fig. 3—figure supplement 2 A1**), which was lost on the HFD (Spearman, r = 0.12, *p* = 0.70; **Fig. 3—figure supplement 2 A2**). TXN treatment restored the negative correlation between caloric intake and plasma TAG (Spearman r = -0.65, *p* = 0.04; **Fig. 3—figure supplement 2 A5**). One explanation for the higher plasma TAG (*p* < 0.01) observed could be that TXN inhibited hepatic lipid uptake, and promoted hepatic lipid export, or both. TAG levels remained in the normal physiological range (40 to 60 mg/dL) for all groups (Bogue *et al*., 2020).

We collected fecal pellets over a 3-day period and measured fecal TAG at the end of the study as an indicator of fecal energy excretion. Fecal TAG levels did not differ among all groups (**Fig. 3—figure supplement 2 B**). No relationship was observed between caloric intake and fecal TAG among or within groups (**Fig. 3—figure supplement 2 B1-5**), suggesting that the attenuated BW gain and hepatic steatosis in TXN- and HXN-treated mice was not related to increased fecal TAG excretion.

### 3. Effects of XN and TXN on food intake frequency, physical activity and energy expenditure

We considered if physical activity level could explain attenuated weight gain of XN- and TXN-treated groups. We differentiated activity measured in the metabolic cages into directed ambulatory locomotion (sum of all locomotion of 1 cm/second or above within the x, y beam-break system) (**Fig. 4A**) and fine movements (e.g., grooming, nesting and scratching) (**Fig. 4B**). In addition, we approximated the ambulatory movement for food consumption by measuring feeding frequency (**Fig. 4C**). In contrast to energy expenditure (**Fig. 4C**), directed ambulatory locomotion was lower in HFD- than LFD-fed mice (**Fig. 4A**), while fine movement level (**Fig. 4B**) and feeding frequency (**Fig. 4C**) were not changed. TXN-treated mice exhibited higher directed ambulatory locomotion and fine movement levels than HFD mice (**Fig. 4A, B**), whereas feeding frequency was unchanged (**Fig. 4C**). XN-treated HFD mice showed higher directed ambulatory locomotion activity and feeding frequency than HFD mice (**Fig. 4A, C**), whereas fine movement activity levels were not affected (**Fig. 4B**).

**Figure 4.**
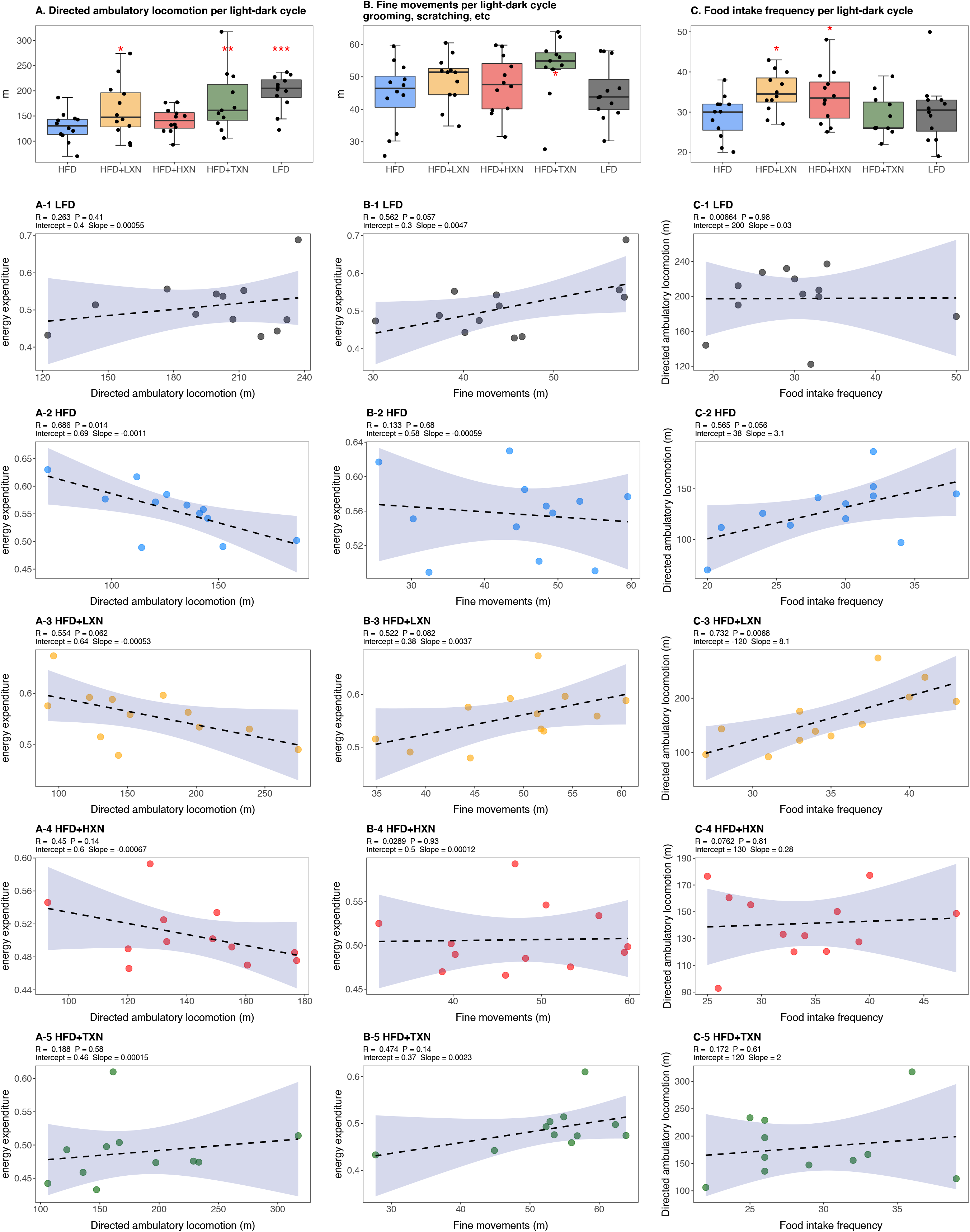
Effects of XN and TXN on food intake frequency, physical activity and energy expenditure. Mice were fed either a LFD (black, n = 12), a HFD (blue, n = 12), HFD+LXN (yellow, n = 12), HFD+HXN (red, n = 12), or HFD+TXN (green, n = 11) for 16 weeks. (A) Directed ambulatory locomotion per 24-hour cycle obtained using a computer controlled indirect calorimetry system. Data expressed as quartiles. (A-1) Relationship between directed ambulatory locomotion and energy expenditure for LFD; (A-2) HFD; (A-3) HFD+LXN; (A-4) HFD+HXN and (A-5) HFD+TXN groups. (B) Fine movements per 24-hour cycle calculated by subtracting directed ambulatory locomotion from sum of all distances traveled within the beam-break system. Data is expressed as quartiles. (B-1) Relationship between fine movements and energy expenditure for LFD; (B-2) HFD; (B-3) HFD+LXN; (B-4) HFD+HXN; and (B-5) HFD+TXN groups. (C) Number of food intake events recorded in metabolic cages. Data expressed as quartiles. (C-1) Relationship between number of food intake events and directed ambulatory locomotion for LFD; (C-2) HFD; (C-3) HFD+LXN; (C-4) HFD+HXN; (C-5) for HFD+TXN groups. Pre-planned general linear model with contrasts were used to calculate *p*-values in A, B and C. \**p* < 0.05, \*\**p* < 0.01, \*\*\**p* < 0.001. Linear regression analyses of energy expenditure versus directed ambulatory locomotion (A1-5), fine movements (B1-5), and directed ambulatory locomotion and number of food intake events (C1-5) in mice were done using stats package version 3.6.2 in R. Blue shading represents 95% CI of the regression line. Absolute value of R, p-value, intercept, and slope for the regression are reported above each corresponding panel. Source files of data used for the analysis are available in the Figure 4—source data 1.

In HFD-fed and LXN-treated mice, directed ambulatory locomotion levels were positively correlated with food frequency (**Fig. 4C2-3**) but negatively correlated with energy expenditure (**Fig. 4A2-3**), suggesting that food-driven activity may account for a major part of total directed ambulatory motion and that these mice spent the majority of their time and energy moving around for food consumption. In summary, HFD-fed mice used more energy for maintaining basal metabolism, body tissue turnover, or digestion as indicated by a higher energy expenditure and lower directed ambulatory locomotion activity than LFD mice. Compared to HFD only mice, XN- and TXN-treated mice had lower energy expenditure and higher directed ambulatory locomotion and fine movement activities, indicating lower energy for maintaining basal metabolism, body tissue turnover, or digestion and remained more physically active than untreated HFD-fed mice.

### 4. TXN attenuates HFD-induced lipid accumulation in white adipose tissue (WAT)

To assess the effect of XN and TXN on lipid accumulation, fat pads from three distinct sites: subcutaneous (sWAT), epididymal (eWAT), and mesenteric (mWAT) adipose tissue were carefully removed and weighed during necropsy. Diet-induced lipid accumulation differed by adipose site. Compared to the LFD, the HFD-induced increase in mWAT fat mass was much greater than the increase in sWAT fat mass (three-fold vs. 2.5-fold increase, respectively), with the smallest increase (15%) observed in eWAT fat mass (**Fig. 5A, C**). Supplementation with HXN (*p* < 0.05), and even more so TXN (*p* < 0.0001), decreased sWAT and mWAT fat mass. A smaller but significant increase in eWAT adipose tissue weight was observed in HXN- and TXN-treated mice (**Fig. 5B**).

**Figure 5.**
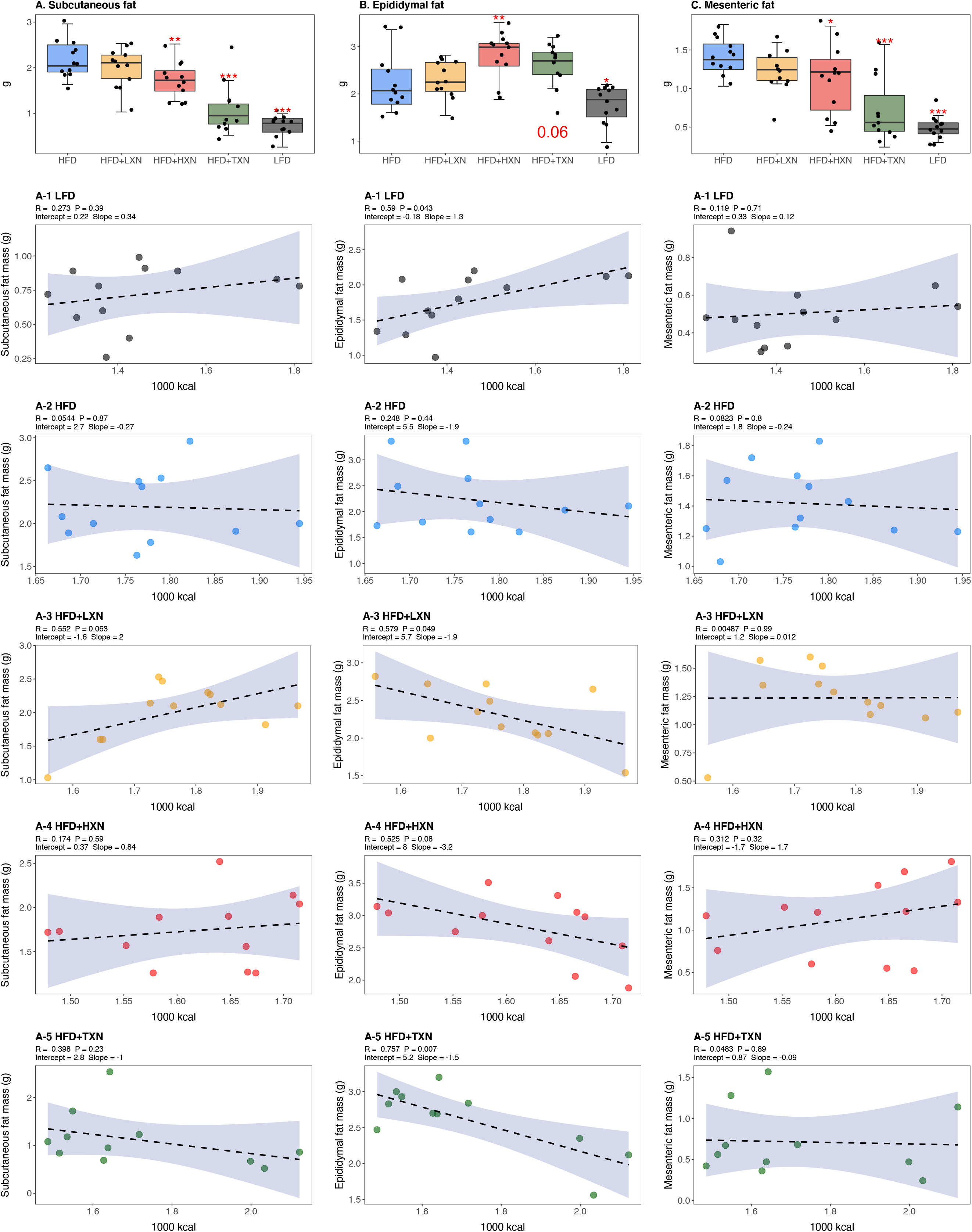
TXN decreases and alters the regional distribution of fat tissue accumulation. Mice were fed either a LFD (black, n = 12), a HFD (blue, n = 12), HFD+LXN (yellow, n = 12), HFD+HXN (red, n = 12), or HFD+TXN (green, n = 11) for 16 weeks. All fat masses were weighed on day of necropsy. (A) sWAT fat mass expressed as quartiles. (A-1) Relationship between sWAT fat mass and total caloric intake over 16 weeks of feeding for LFD; (A-2) HFD; (A-3) HFD+LXN; (A-4) HFD+HXN; and (A-5) HFD+TXN groups. (B) eWAT fat mass expressed as quartiles. (B-1) Relationship between eWAT fat mass and total caloric intake over 16 weeks of feeding for LFD; (B-2) HFD; (B-3) HFD+LXN; (B-4) HFD+HXN; and (B-5) HFD+TXN groups. (C) mWAT fat mass expressed as quartiles. (C-1) Relationship between mWAT fat mass and total caloric intake over 16 weeks of feeding for LFD; (C-2) HFD; (C-3) HFD+LXN; (C-4) HFD+HXN (with removal of two outliers); (C-5) and for HFD+TXN groups. Pre-planned general linear model with contrasts were used to calculate *p*-values in A, B and C. \**p* < 0.05, \*\**p* < 0.01, \*\*\**p* < 0.001. Linear regression analyses of total calories versus sWAT (A1-5), eWAT (B1-5), and mWAT fat masses (C1-5) in mice were done using stats package version 3.6.2 in R. Blue shading represents 95% CI of the regression line. Absolute value of R, p-value, intercept, and slope for the regression are reported above each corresponding panel. Source files of data used for the analysis are available in the Figure 5—source data 1.

Caloric intake across diets was positively correlated with sWAT (r = +0.47; *p* = 0.0002; **Fig. 5A**) and mWAT fat mass (r = +0.39; *p* = 0.002; **Fig. 5C**), but no relationship was observed within XN- or TXN-treated groups (**Fig. 5A3-5, C3-5**), indicating lipid accumulation in sWAT and mWAT fat depots was primarily linked to diet rather than the amount of food consumed. In eWAT adipose depot, we observed the opposite. Unlike sWAT and mWAT fat depots, caloric intake across diets was not correlated with eWAT fat mass (r = +0.03; *p* = 0.82; **Fig. 5B**). Instead, a positive correlation between caloric intake and eWAT fat mass was found within LFD- fed mice (**Fig. 5B1**), and a negative correlation between caloric intake and eWAT fat mass was observed in both XN- and TXN-treated mice (**Fig. 5B3-5**). No correlation was found in HFD-fed control mice (**Fig. 5B2**). These observations are consistent with distinct WAT depots in mice differing in expandability (van Beek *et al*., 2015).

### 5. HXN and TXN protect against NAFLD on a high-fat diet

NAFLD is characterized by accumulation of number and size of intrahepatic microvesicular and macrovesicular lipid vacuoles. Mice on a LFD diet possessed hepatic lipid vacuoles and resembled livers of LDLR^-/-^ mice on a similar synthetic diet (Lytle and Jump, 2016); however their liver to BW ratio of about 4% was in a normal healthy range (Lytle, Wong and Jump, 2017). HFD fed mice had many smaller lipid vacuoles (**Fig. 6A**). Supplementation with XN decreased in a dose-dependent manner the number and size of intrahepatic lipid vacuoles in HFD mice. Supplementation with TXN almost completely prevented hepatic lipid vacuole accumulation in HFD mice, resulting in less lipid accumulation than in LFD mice.

**Figure 6.**
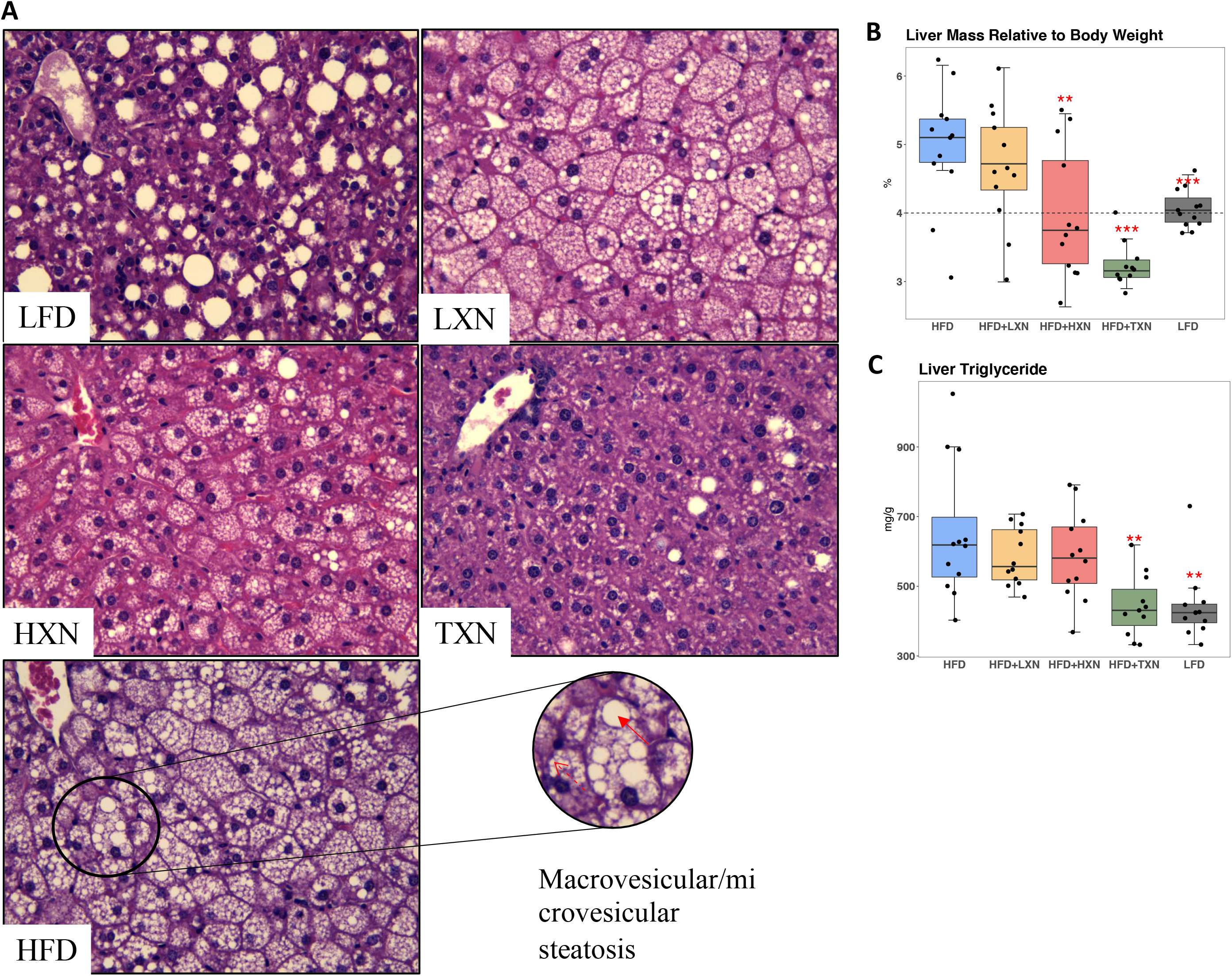
TXN prevents HFD induced liver steatosis in mice. Mice were sacrificed at the end of the study and liver samples were freshly collected and processed. (A) Representative histological images of H&E staining of liver sections. An enlarged image representative of a liver section from a HFD fed mouse is shown as a circle on the bottom right. Macrovesicular steatosis or large lipid droplets are indicated by the red bold arrow; microvesicular steatosis or small lipid droplets are indicated by the broken red line arrow. Liver mass to BW ratio. (C) Hepatic triglyceride content. P-values of orthogonal *a priori* comparisons of the HFD versus each of the other groups are shown. \*\**p* < 0.01, \*\*\**p* < 0.001. Source files of data used for the analysis are available in the Figure 6—source data 1 and source data 2.

The liver to BW ratio is an indicator of NAFLD with a ratio above 4% indicating NAFLD (Lytle, Wong and Jump, 2017). The majority of mice (10 of 12) on a HFD diet had a liver to BW ratio above 4.5%, whereas all LFD mice had a liver to BW ratio between 3.8 and 4.3% (**Fig. 6B**). Supplementation with HXN decreased the number of mice with a liver to BW ratio above 4% to four of 12 mice and all TXN-supplemented mice had a liver to BW ratio below 3.6% except for one, which had a liver to BW ratio of 4%. These data are consistent with TXN and, to a smaller extent, HXN reducing NAFLD. Hepatic lipid extracts from TXN-supplemented HFD mice and LFD-fed mice had lower liver triglyceride concentrations than from mice fed with HFD, LXN or HXN (**Fig. 6C**).

Another indicator of NAFLD is the liver area occupied by lipids; the histological lower cut-off for NAFLD is over 5% of liver area (Brunt and Tiniakos, 2010). Using this cut-off, all control HFD mice had NAFLD and 10 out of 12 LFD mice had NAFLD (**Fig. 3B**). Both HXN- and TXN supplementation decreased liver lipid accumulation on a HFD by two-fold, as three out of 12 HXN-supplemented mice and five out of 11 TXN-supplemented mice had less than 5% lipid area and nine out of 11 TXN-supplemented mice had less than 10% lipid area (**Fig. 3B**). In comparison, only one out of 12 HFD control mice and seven out of 12 HXN-supplemented mice were below 10% lipid area. The supplement-induced decrease was independent of caloric intake (**Fig 3. B3-5**).

### 5. RNA-seq reveals suppression of hepatic FA biosynthesis processes and pathways by HXN and TXN treatments

We conducted RNA-seq analysis of the livers obtained from mice after 16 weeks on the diet to determine transcriptional mechanisms by which HXN and TXN supplementation could ameliorate hepatic steatosis induced by HFD. Gene counts were calculated to quantify gene expression in the four diet groups LFD, HFD, HFD+HXN, and HFD+TXN. The differentially expressed genes (DEGs) were determined using a false discovery rate (FDR) cutoff of < 0.4, as compared to HFD.

To visualize expression patterns of DEGs in the four groups we used hierarchical clustering with a heat map (**Fig. 7A**). The DEGs clustered into two major types, one with higher expression (red) in the LFD and HFD groups but lower expression (blue) in the HXN and TXN groups and the other with lower expression in the LFD and HFD groups but higher expression in the HXN and TXN groups (**Fig. 7A**). Individual mice clustered into two major nodes. All HFD mice clustered with six LFD and four HXN mice and all TXN mice clustered with six HXN and four LFD mice (**Fig. 7A**). This likely reflects the variability observed in phenotypic outcomes (Fig. 6B). The volcano plot analysis of gene expression revealed that both HXN and TXN treatments induced significant changes in gene expression compared with the HFD group (**Fig. 7B**). TXN treatment had the greatest effect with 295 identified DEGs while HXN treatment only resulted in six DEGs. We identified 212 DEGs in comparing the LFD and HFD groups.

**Figure 7.**
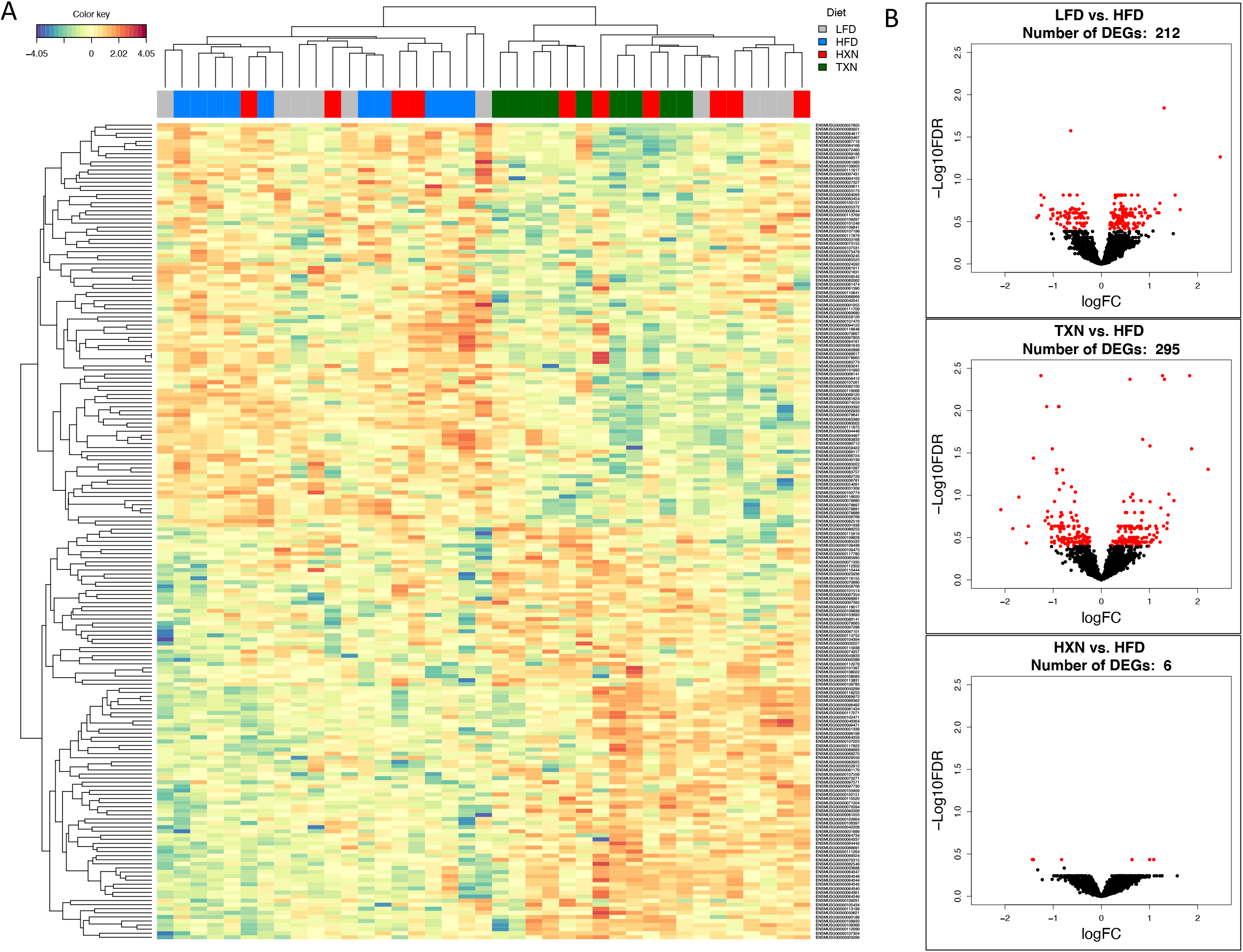
TXN treatment significantly alters liver transcriptome of mice after 16 weeks of feeding. (A) Hierarchical clustering of the top 200 differentially expressed genes (DEGs) in each treatment group (labeled at the top right corner: gray indicates LFD group, blue indicates HFD, red indicates HXN and green indicates TXN.) as determined by RNAseq analysis. Color key is based on the log2 fold change. (B) Volcano plots show DEGs (red dots) in the comparison of different treatment groups. Source files of data used for the analysis are available in the Figure 7—source data 1.

We next conducted gene ontology (GO) enrichment and pathway analysis of DEGs using Enrichr (Chen *et al*., 2013). We assigned the DEGs in the TXN treatment group to GO terms describing biological processes. The enriched GO terms and pathways with adjusted p values < 0.05 are summarized in **Figure 8** **and** **Figure 8** **source data**. GO enrichment analysis indicated that TXN treatment significantly downregulated genes involved in biological processes including xenobiotic catabolism, FA metabolism, glucose metabolism and regulation of lipid metabolism (**Fig. 8**, top panel). Furthermore, KEGG pathway analysis demonstrated that TXN upregulated expression of genes in six pathways including complement and coagulation cascades, prion diseases, steroid hormone biosynthesis, arachidonic acid metabolism, retinol metabolism and linoleic acid metabolism (**Fig. 8**, bottom right panel). Many of these included genes encoding Cyp450 enzymes. On the other hand, expression of genes in 25 KEGG pathways were significantly downregulated by TXN treatment compared to HFD (**Fig. 8**, bottom left panel). The top 10 significantly enriched KEGG pathways based on statistical significance and combined score ranking included the biosynthesis of unsaturated FAs, glutathione metabolism, amino sugar and nucleotide sugar metabolism, glycolysis and gluconeogenesis, pentose phosphate pathway, fluid shear stress and atherosclerosis, chemical carcinogenesis, drug metabolism, FA elongation and the PPAR signaling pathway.

**Figure 8.**
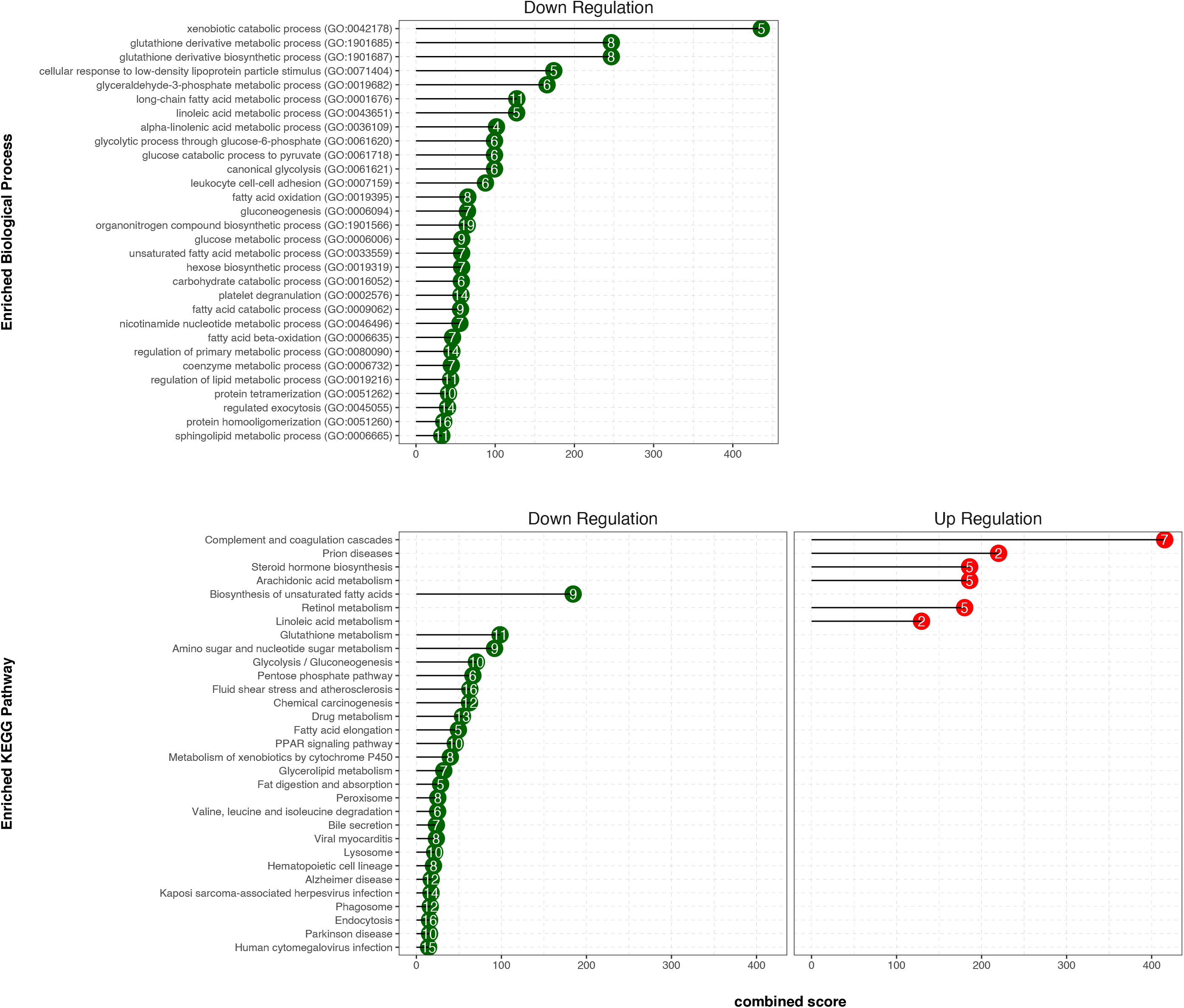
TXN decreases expression of numerous gene ontology and KEGG pathways. Analysis of DEGs from the livers of mice that consumed a HFD+TXN versus a HFD revealed mostly downregulation of biological processes and KEGG pathways. The significant (adjusted p < 0.05) enriched biological process terms in gene ontology (upper panel) and enriched KEGG pathways (lower panel) were selected by Enrichr Tools based on significance and combined scores. The number inside each lollipop represents the number of identified DEG genes in that specific biological process or KEGG pathway. Source files of data used for the analysis are available in the Figure 8—source data 1.

### 6. Identification of key hepatic genes regulated by TXN and involved in ameliorating hepatic steatosis

We implemented SVM to identify a set of signature genes that can distinguish TXN-treated mice from HFD-fed control mice. Briefly, we used the DaMirSeq R package to determine a set of genes whose principal components best correlated with TXN treatment by performing backward variable elimination with partial least-squares regression and removing redundant features by eliminating those that were very highly correlated (Chiesa, Colombo and Piacentini, 2018). Repeating this process 30 times, we used all 13 genes identified here as input into our SVM models (**Fig. 9**, left panel). Genes identified classified HFD- and TXN-fed mice into two distinct groups (**Fig. 9**, right panel). Eight of 13 genes showed significant, differential expression between TXN and HFD diet samples (**Table 1**). Consistent with the GO analysis, three of the eight genes – uncoupling protein 2 (*Ucp2*), cell death-inducing DFFA-like effector c (*Cidec*), and monoacylglycerol O-acyltransferase 1 (*Mogat1*) – are involved in lipid metabolism and known target genes of PPARγ (Medvedev *et al*., 2001; Kim *et al*., 2008; Matsusue *et al*., 2008); (Bugge *et al*., 2010; Karbowska and Kochan, 2012; Wolf Greenstein *et al*., 2017).

**Figure 9.**
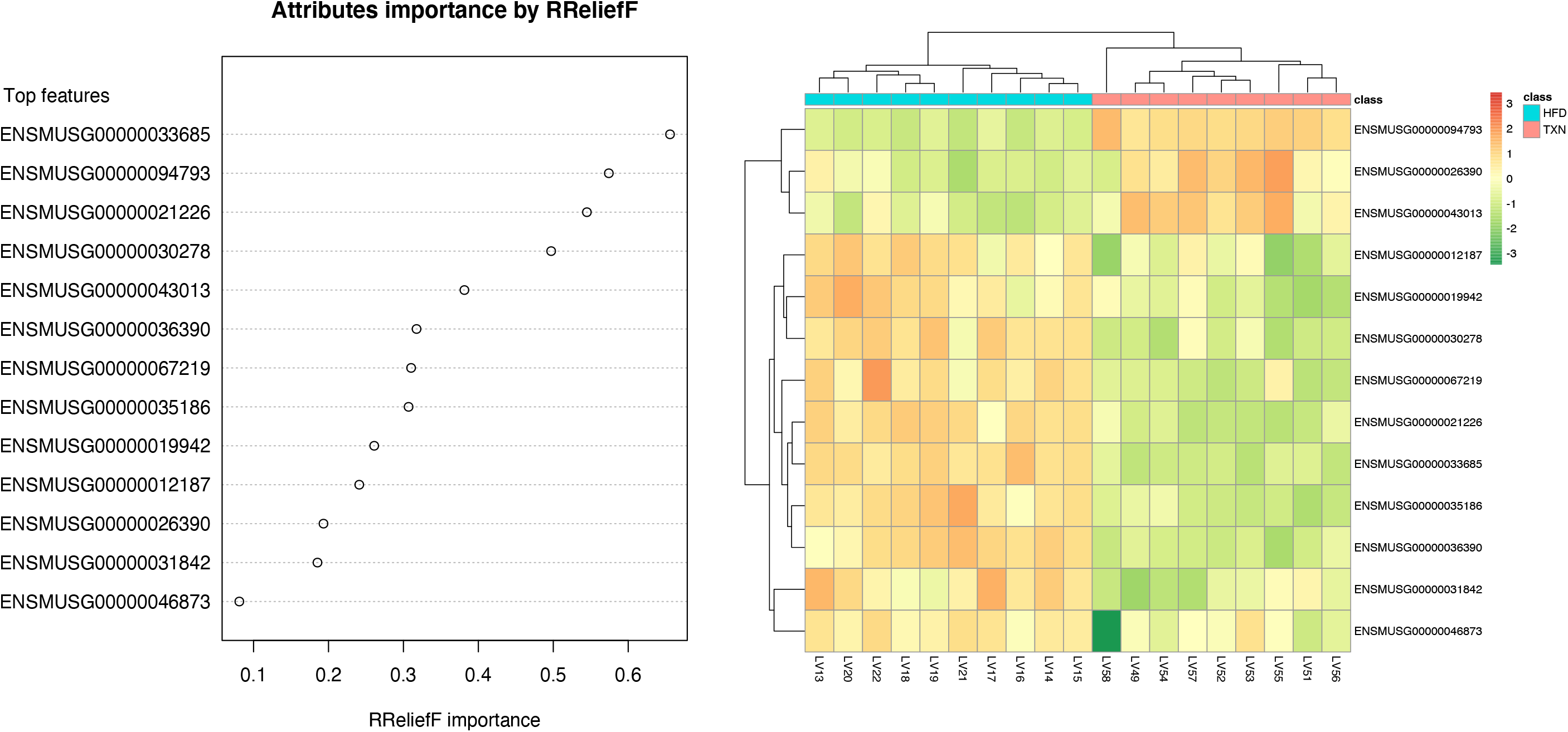
SVM identified signature genes that distinguish mice that consumed TXN. Left panel: the dot chart shows the top 13 genes, sorted by RReliefF importance score. This plot was used to select the most important predictors to be used for classification. Right panel: Colors in the heatmap highlight the gene expression level in fold change: color gradient ranges from *dark orange*, meaning “upregulated”, to *dark green*, meaning “downregulated”. On the top of the heatmap, horizontal bars indicate HFD (blue) and HFD+TXN (pink) treatments. On the top and on the left side of the heatmap the dendrograms obtained by Spearman’s correlation metric are shown. Plots were produced with DaMiRseq R package 1.10.0. Source files of data used for the analysis are available in the Figure 9—source data 1.

**Table 1.**
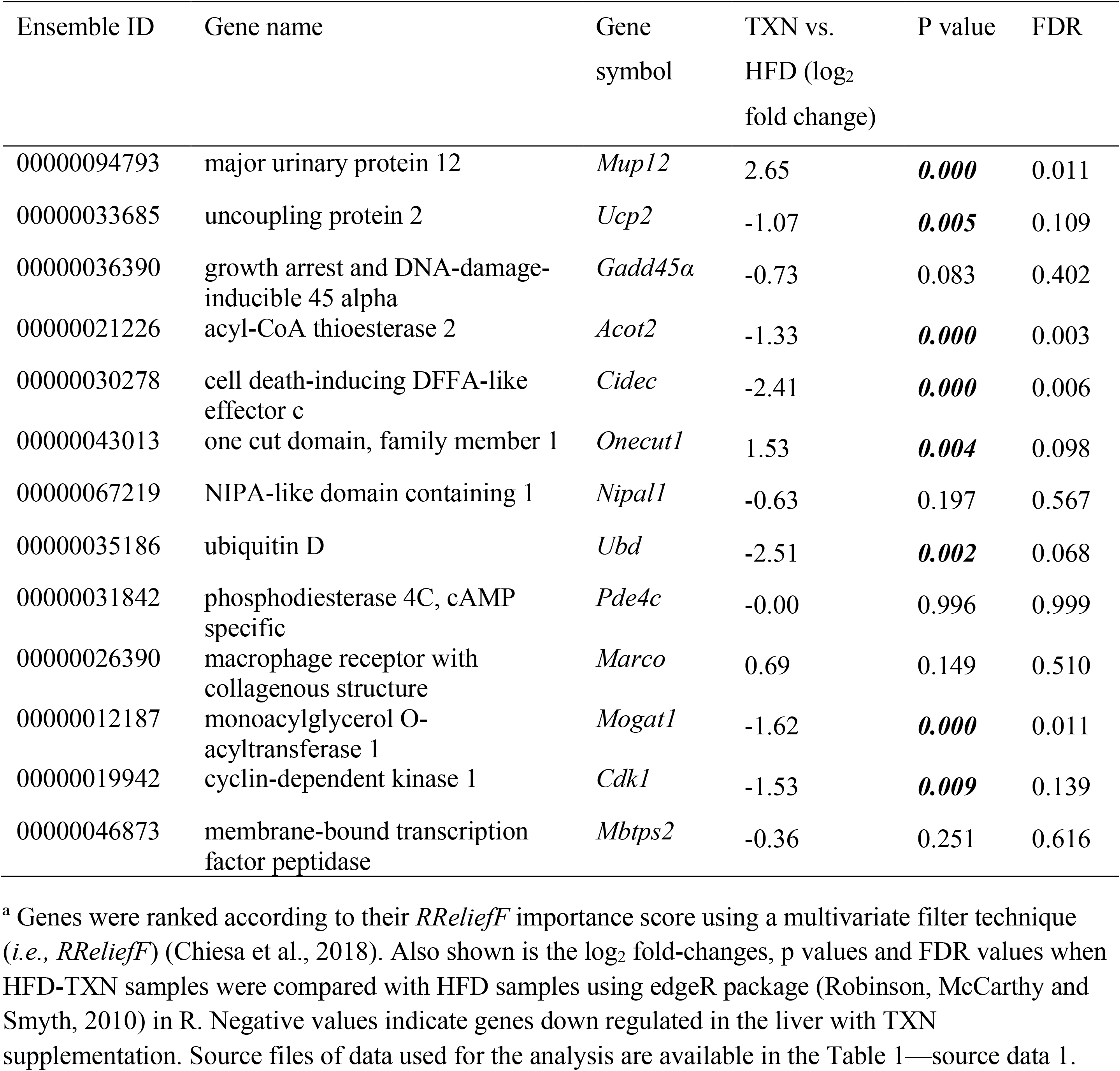
Thirteen genesa used to distinguish TXN transcriptome from HFD transcriptome.

| Ensemble ID | Gene name | Gene symbol | TXN vs. HFD (log<sub>2</sub> fold change) | P value | FDR |
| --- | --- | --- | --- | --- | --- |
| 00000094793 | major urinary protein 12 | *Mup12* | 2.65 | **0.000** | 0.011 |
| 00000033685 | uncoupling protein 2 | *Ucp2* | -1.07 | **0.005** | 0.109 |
| 00000036390 | growth arrest and DNA-damage-inducible 45 alpha | *Gadd45a* | -0.73 | 0.083 | 0.402 |
| 00000021226 | acyl-CoA thioesterase 2 | *Acot2* | -1.33 | **0.000** | 0.003 |
| 00000030278 | cell death-inducing DFFA-like effector c | *Cidec* | -2.41 | **0.000** | 0.006 |
| 00000043013 | one cut domain, family member 1 | *Onecut1* | 1.53 | **0.004** | 0.098 |
| 00000067219 | NIPA-like domain containing 1 | *Nipal1* | -0.63 | 0.197 | 0.567 |
| 00000035186 | ubiquitin D | *Ubd* | -2.51 | **0.002** | 0.068 |
| 00000031842 | phosphodiesterase 4C, cAMP specific | *Pde4c* | -0.00 | 0.996 | 0.999 |
| 00000026390 | macrophage receptor with collagenous structure | *Marco* | 0.69 | 0.149 | 0.510 |
| 00000012187 | monoacylglycerol O-acyltransferase 1 | *Mogat1* | -1.62 | **0.000** | 0.011 |
| 00000019942 | cyclin-dependent kinase 1 | *Cdk1* | -1.53 | **0.009** | 0.139 |
| 00000046873 | membrane-bound transcription factor peptidase | *Mbtps2* | -0.36 | 0.251 | 0.616 |

<sup>a</sup> Genes were ranked according to their *RReliefF* importance score using a multivariate filter technique (*i.e.*, *RReliefF*) (Chiesa et al., 2018). Also shown is the log<sub>2</sub> fold-changes, p values and FDR values when HFD-TXN samples were compared with HFD samples using edgeR package (Robinson, McCarthy and Smyth, 2010) in R. Negative values indicate genes down regulated in the liver with TXN supplementation. Source files of data used for the analysis are available in the Table 1—source data 1.

We then confirmed expression of these genes using RT-qPCR. Consistent with RNAseq results, TXN-treated mice had significantly lower expression of major PPARγ target genes *Cidec*, *Mogat1* and *Pparγ2,* a predicted PPAR target gene (Fang *et al*., 2016) (**Fig. 10** top panel). Moreover, we observed significantly strong positive correlations between the expression of these three genes (**Fig. 10**, bottom panel). The above results suggest TXN treatment inhibits the PPARγ pathway – a key pathway involved in hepatic lipid metabolism.

**Figure 10.**
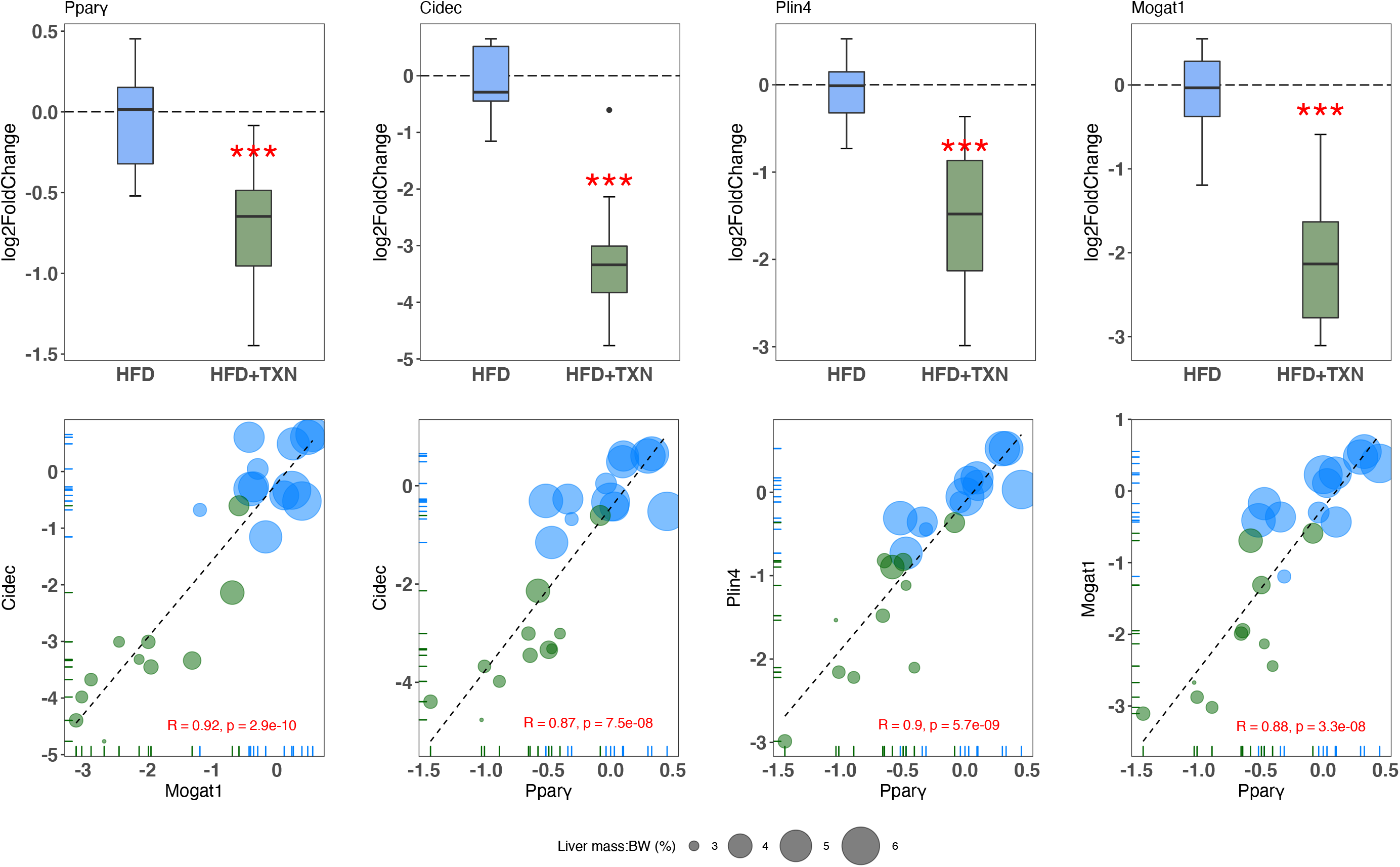
TXN-treated mice show significantly lower expression of PPARγ and target genes. Top panel: reduction of HFD-induced *Pparγ2*, *Cidec*, and *Mogat1* expressions in the liver by TXN administration. Mice were sacrificed after 16-week of HFD (blue, n = 12) or HFD+TXN (dark green, n = 11) feeding. Liver tissues were harvested, and total RNA was extracted. Relative mRNA levels of selected genes were determined by real time PCR. Gene expression is expressed in log_2_ fold change as quartiles. \*\*\**p* ≤ 0.001, t-test. Bottom panel: Pearson correlation between *Pparγ2* and *Cidec* or *Mogat1* expression. Data are presented in log_2_ fold change; bubble size represents liver mass to BW ratio. • indicates sample outside value, which is > 1.5 times the interquartile range beyond upper end of the box. Source files of data used for the analysis are available in the Figure 10—source data 1.

### 7. XN and TXN attenuate intracellular lipid content in 3T3-L1 adipocytes in a dose dependent manner

We hypothesized that TXN and XN antagonizes the PPARγ receptor, which would explain the decreased expression of its target genes. To test our hypothesis, we utilized 3T3-L1 murine fibroblast cells, which depend on PPARγ activity to differentiate into adipocytes (Tamori *et al*., 2002). XN and its derivatives are cytotoxic to some cells and to ensure that we used concentrations that were not cytotoxic to 3T3-L1 adipocytes, we tested an escalating dose of XN and TXN (Strathmann and Gerhauser, 2012). 3T3-L1 cells were treated with 0.1% DMSO, 1 µM rosiglitazone (RGZ), 1 µM GW9662, XN (5, 10 and 25 µM), TXN (5, 10 and 25 µM), 25 µM XN + 1 µM RGZ or 25 µM TXN + 1 µM RGZ for 48 h. After treatments, we determined the number of live cells using an MTT assay. XN and TXN were only significantly cytotoxic for 3T3-L1 cells at a dose of 50 µM (data not shown). While it is difficult to translate *in vivo* doses to *in vitro* doses, based on previous *in vitro* studies (Yang *et al*., 2007; Samuels, Shashidharamurthy and Rayalam, 2018) and our current cell viability data, we selected low (5 µM), medium (10 µM) and high (25 µM) concentrations of XN and TXN for the subsequent experiments where cell viability was greater than 90% (data not shown).

Murine preadipocyte 3T3-L1 differentiation and adipogenesis was induced by addition of dexamethasone, IBMX and insulin which strongly induced intracellular lipid accumulation (**Fig. 11, A2-3**). Addition of XN significantly attenuated intracellular lipid levels in a dose-dependent manner (**B1-3**). Like XN, TXN also strongly inhibit intracellular lipid accumulation (**C1-3**).

**Figure 11.**
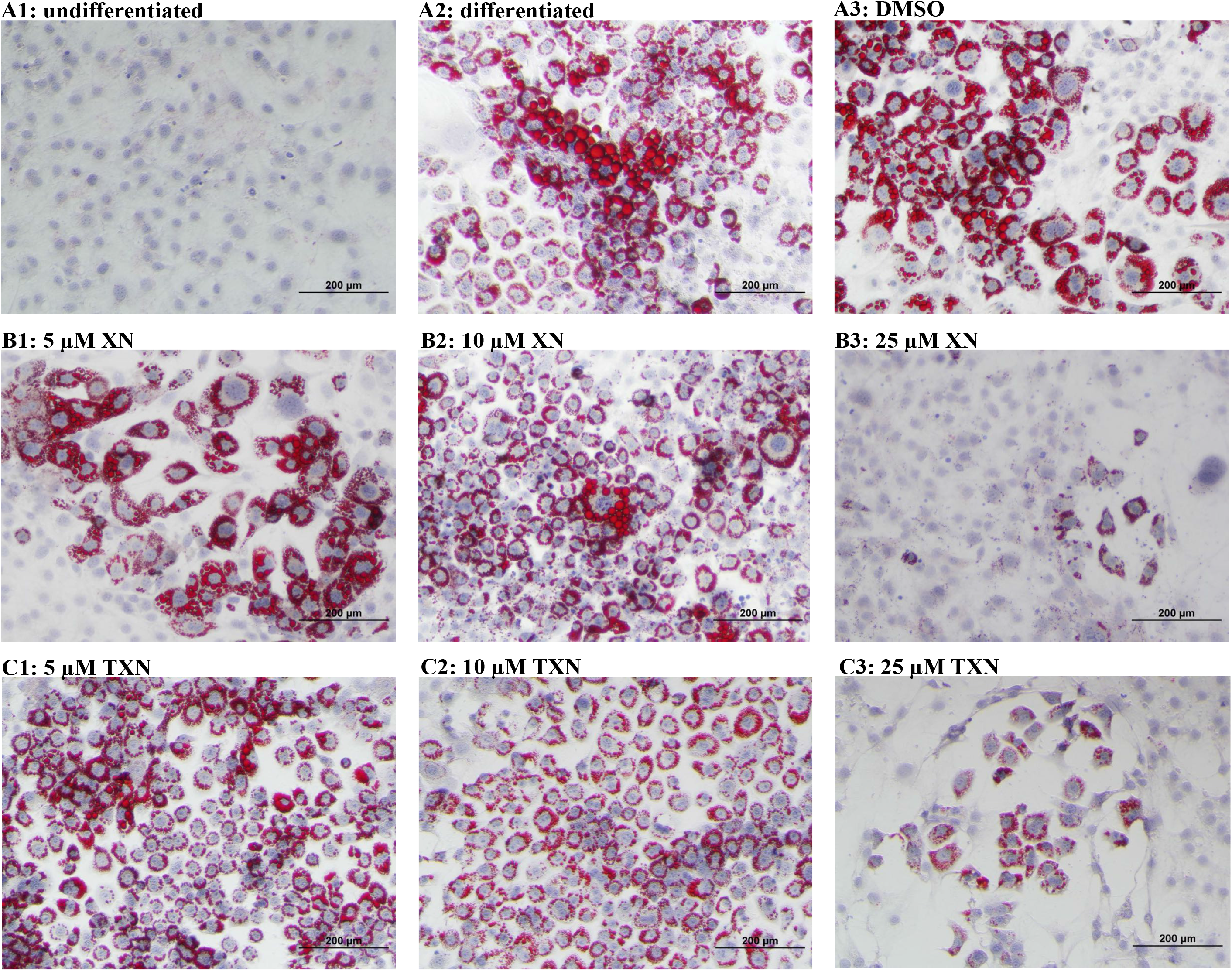
XN and TXN inhibit intracellular lipid accumulation in 3T3-L1 cells. 3T3-L1 cells (1×10^6^ per well) in 12-well plates were cultured with either DMEM (A1), differentiation medium (DM) (A2), DM plus DMSO (A3), DM plus 5 µM XN (B1), DM plus 10 µM XN (B2), DM plus 25 µM XN (B3), DM plus 5 µM TXN (C1), DM plus 10 µM TXN (C2), or DM plus 25 µM TXN (C3). Cells were stained with oil red O to identify lipids at day 7 post-differentiation.

### 8. XN and TXN inhibit RGZ-induced adipocyte differentiation in 3T3-L1 cells in a dose dependent manner

RGZ is a known potent PPARγ agonist used as an insulin-sensitizing agent. To test the hypothesis that XN and TXN may antagonize a known PPARγ ligand, we determined if the compounds would block RGZ-induced PPARγ actions (**Fig. 12**). RGZ strongly induced the differentiation (**Fig. 12**, **A1**), and GW 9662, a potent PPARγ antagonist, inhibited the RGZ-induced differentiation (**Fig. 12**, **A2**). We also observed that both XN (**Fig. 12**, **B1-3**) and TXN (**Fig. 11**, **C1-3**) suppressed RGZ-induced differentiation in a dose dependent manner. At 25 µM concentration, the RGZ-induced differentiation was largely blocked (**Fig. 12**, **B3, C3**), suggesting that XN and TXN may interfere or even compete with binding of RGZ to the PPARγ receptor.

**Figure 12.**
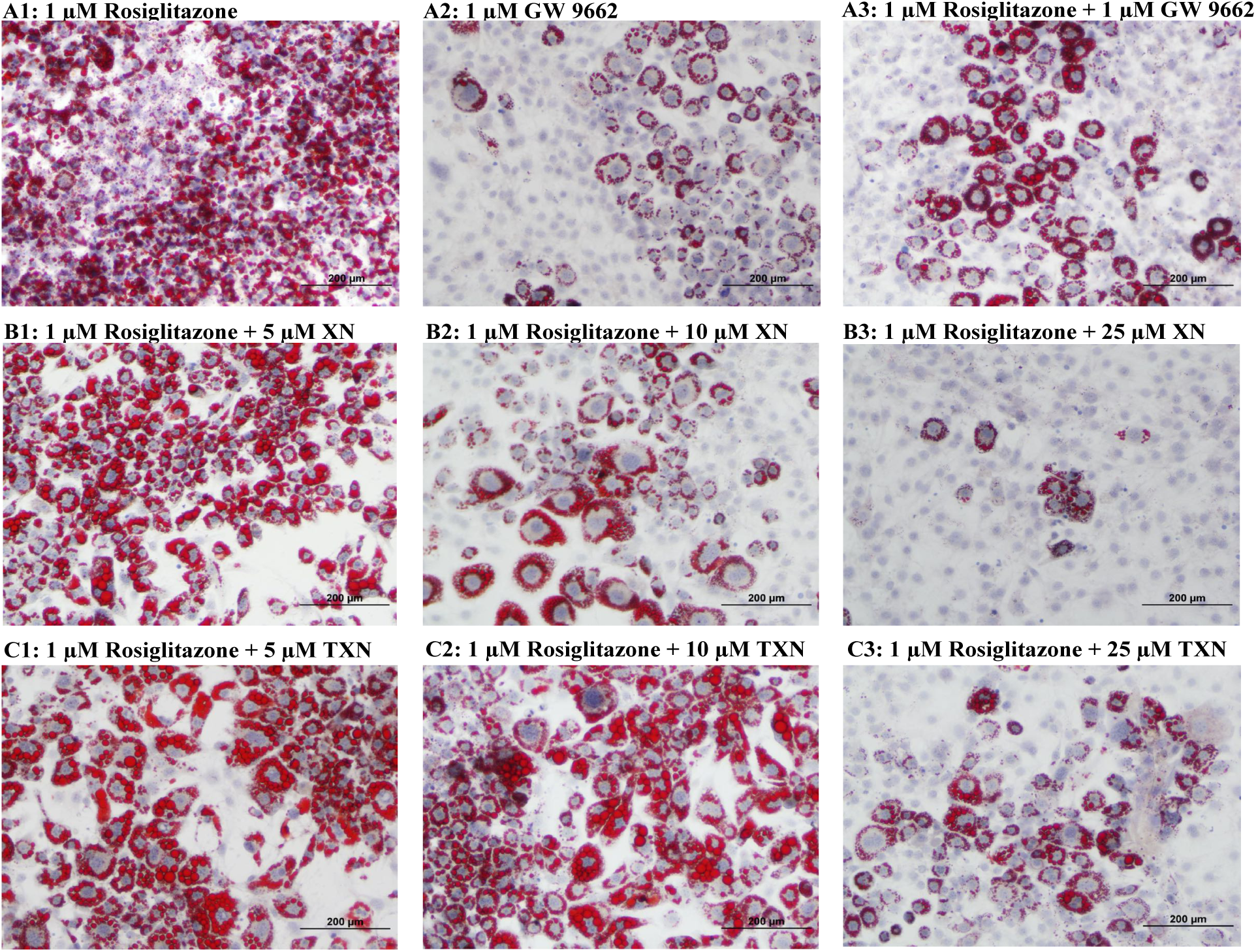
XN and TXN diminished the lipid accumulation in 3T3-L1 cells. 3T3-L1 cells (1×10^6^ per well) in 12-well plates were cultured with either DM plus 1µM rosiglitazone (A1), DM plus 1µM GW 9662 (A2), DM plus 1µM rosiglitazone and 1µM GW9662 (A3), DM plus 1 µM rosiglitazone and 5 µM XN (B1), DM plus 1 µM rosiglitazone and 10 µM XN (B2), DM plus 1 µM rosiglitazone and 25 µM XN (B3), DM plus 1 µM rosiglitazone and 5 µM TXN (C1), DM plus 1 µM rosiglitazone and 10 µM TXN (C2), or DM plus 1 µM rosiglitazone and 25 µM TXN (C3). Cells were stained with oil red O to identify lipids at day 7 post-differentiation.

### 9. XN and TXN downregulate genes regulated by PPARγ in 3T3-L1 cells

To elucidate the effect of XN and TXN on PPARγ action at the transcriptional level, we measured the expression of several known PPARγ target genes using RT-qPCR on samples 7 days post 25 µM XN or TXN treatment. Consistent with the decrease of intracellular lipid content in **Figures 10** and **11**, the expression of PPARγ and its target genes at 7 days post treatment were significantly downregulated by XN and TXN treatments (**Table 2**). Cells treated with 1 μM GW 9662, a PPARγ antagonist, did not significantly reverse the RGZ-induced upregulation of these genes. Cells treated with either 25 µM XN or TXN significantly reversed the RGZ-induced upregulation of *Cd36* (*p* < 0.001, *p* < 0.001), *Fabp4* (*p* < 0.001, *p* < 0.001), *Mogat1* (*p* < 0.001, *p* < 0.01), *Cidec* (*p* < 0.001, *p* < 0.001), *Plin4* (*p* < 0.001, *p* < 0.001), *Fgf21* (*p* < 0.01, *p* < 0.01). Taken together, these data above suggest that XN and TXN antagonize PPARγ at the transcriptional level to block 3T3-L1 differentiation.

**Table 2.** Adipocyte gene expression at day 7 post-differentiation.

| Gene | Log<sub>2</sub> (Fold Change) |  |  |  | p-values vs. RGZ |  |  |
| --- | --- | --- | --- | --- | --- | --- | --- |
|  | RGZ (cont) | RGZ + GW9662 | RGZ + XN | RGZ + TXN | RGZ + GW9662 | RGZ + XN | RGZ + TXN |
| *Pparγ2* | Ref. | -0.11 | -1.93 | -1.53 | 0.30 | < 0.001 | < 0.001 |
| *Cd36* |  | -0.18 | -9.10 | -4.36 | 0.25 | < 0.001 | < 0.001 |
| *Fabp4* |  | -0.12 | -7.94 | -4.08 | 0.43 | < 0.001 | < 0.001 |
| *Mogat1* |  | -0.11 | -4.16 | -3.59 | 0.42 | < 0.001 | < 0.01 |
| *Cidec* |  | -0.18 | -10.10 | -4.46 | 0.40 | < 0.001 | < 0.001 |
| *Plin4* |  | -0.10 | -3.01 | -2.32 | 0.48 | < 0.001 | < 0.001 |
| *Fgf21* |  | 0.03 | -0.99 | -1.08 | 0.40 | < 0.01 | < 0.01 |

3T3-L1 differentiation was induced by IBMX, dexamethasone, insulin and 1 μM RGZ plus the addition of 1 μM GW9662, 25 μM XN, or 25 μM TXN for 48 hours. After 48 hours, the old media was removed and fresh DMEM was replenished for continuing differentiation. Gene expression was measured at day 7 post-differentiation using qRT-PCR. ΔCT = CT(target gene) – CT(reference gene). $\Delta\Delta$ CT = ΔCT(treated sample) – ΔCT(untreated sample/control average). Fold change = $2^{-\Delta\Delta$ CT}. Statistics were performed on $\Delta\Delta$ CT values. Source files of data used for the analysis are available in the Table 2—source data 1.

### 10. XN and TXN antagonize ligand binding to PPARγ

Based on the inhibition of RGZ induced adipocyte differentiation, and expression of PPARγ target genes, we postulated that XN and TXN bind to the PPARγ ligand binding domain and interfere with agonist binding. To test this hypothesis, we first performed a competitive binding assay using a PPARγ TR-FRET assay. Both XN and TXN displaced a labelled pan-PPARγ ligand in a dose-dependent manner with IC_50_ values of 1.97 µM (**Fig. 13B**) and 1.38 µM (Fig. 13C), respectively. Oleic acid, the most abundant FA ligand in the HFD diet (**Table 4**), had an IC_50_ value of 16.6 µM. XN and TXN had similar IC_50_ values as the PPARγ ligand PGZ, a drug used to improve insulin sensitivity and type 2 diabetes, and a natural ligand, arachidonic acid (Chen *et al*., 2012).

**Figure 13.**
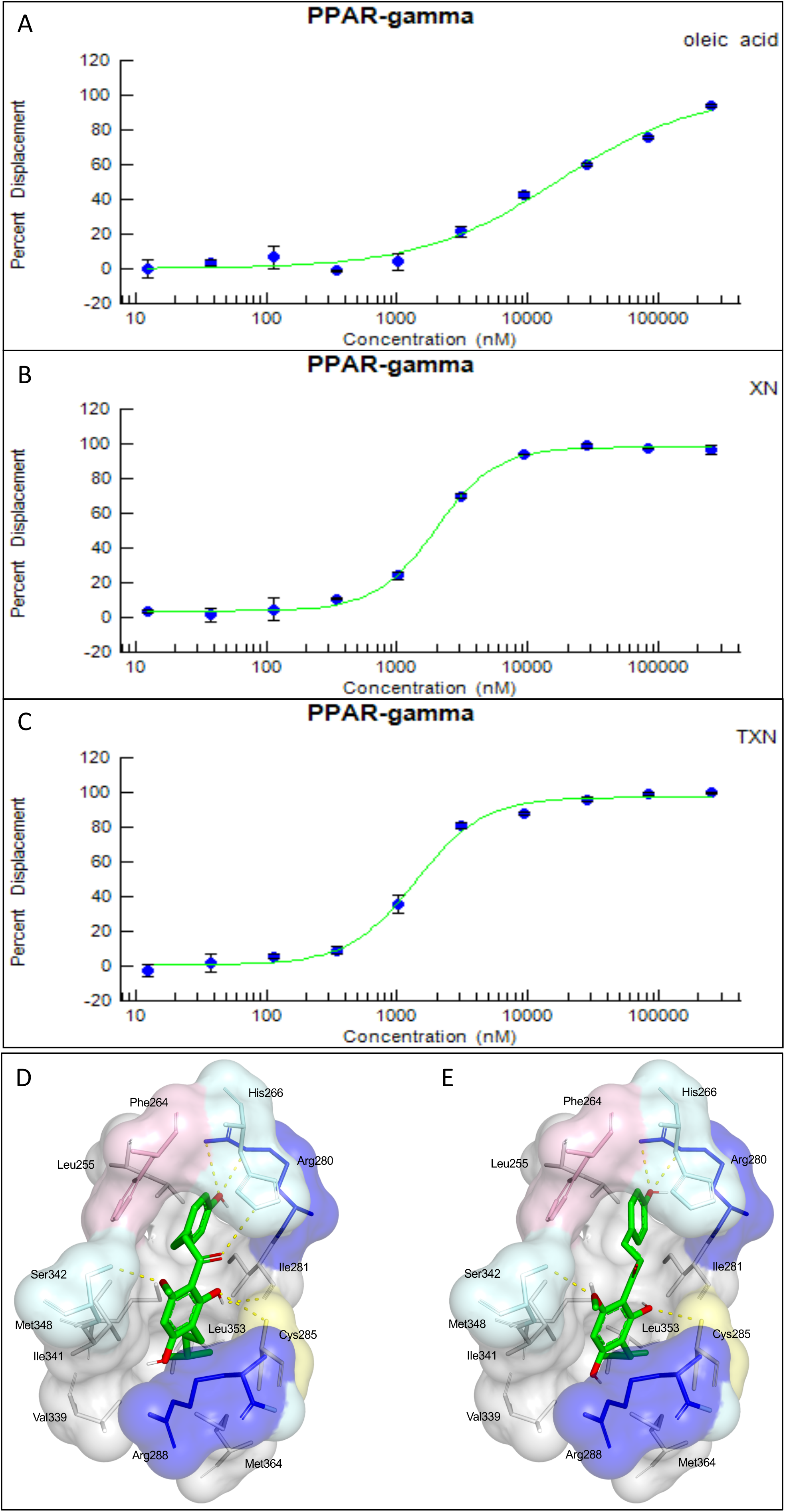
XN and TXN are ligands for PPARγ. A PPARγ nuclear receptor competitive binding assay based on time-resolved fluorescence resonance energy transfer (TR-FRET) was performed. The IC_50_ values for each compound was determined by % displacement of a pan-PPARγ ligand. (A) Oleic acid IC_50_ 16.6 µM. (B) XN IC_50_ 1.97 µM. (C) TXN IC_50_ 1.38 µM. Molecular docking studies show TXN and XN fit into the human PPARγ binding site. PPARγ residues containing atoms involved in hydrophobic interactions are shown. Yellow dashes indicate hydrogen bonds, amino acids colored as hydrophobic (grey), aromatic (pink), polar (cyan), basic (blue), or cysteine (yellow). (D) TXN (E) XN. Source files of data used for the analysis are available in the Figure 13—source data 12.

To obtain further insights into the interaction of XN and TXN with PPARγ, we analyzed the nature of binding between the PPARγ-LBD and XN/TXN using molecular docking to confirm the putative binding pose and position of XN/TXN and to estimate the relative binding affinities of various ligands for PPARγ. To verify the robustness of our docking protocol, resveratrol was re-docked into the bound structure of PPARγ, reproducing the binding pose and orientation found in the crystal structure of the complex (PDB ID: 4JAZ). The best docked position of TXN occupies the binding site of PPARγ, exhibiting many non-bonded interactions involving side chain atoms in Leu255, Phe264, Gly284, Cys 285, Arg288, Val339, Ile 341, Met348, and Met364 (Fig. 13D). The side chains of His266, Arg280, and Ser342 and the main chain carbonyl oxygen atom of Ile281 are well positioned to make electrostatic/hydrogen bonds with the hydroxyl protons and oxygen atoms of the bound TXN molecule. We observed many of the same hydrophobic interactions in the simulated PPARγ-XN (Fig. 13E) and PPARγ-oleic acid complexes, and potential electrostatic interactions between His266 and Glu343, or with Arg280 and XN or oleic acid, respectively. The relative binding affinities, ranked in decreasing value of their negative binding energies were, in order, TXN, XN, and oleic acid, consistent with the TR-FRET binding results.

## Discussion

### XN and TXN are effective in suppressing development of diet-induced steatosis

Low cost natural products like XN are of particular interest for treating obesity and NAFLD due to their availability, safety and efficacy. XN and its derivatives appear to function through multiple mechanisms of action and this polypharmacological effect may enhance their effectiveness. Three studies propose that XN improves diet-induced hepatic steatosis by suppressing SREBP1c mRNA expression and SREBP activation (Yui, Kiyofuji and Osada, 2014; Miyata *et al*., 2015; Takahashi and Osada, 2017). We also observed a decrease in hepatic SREPB1c expression with TXN treatment. Others propose mechanisms include inhibiting pro-inflammatory gene expression (Dorn *et al*., 2010; Mahli *et al*., 2019), inducing AMPK activation in the liver and skeletal muscle (Costa *et al*., 2017), and enhanced FA oxidation (Kirkwood *et al*., 2013). In this study, using a combination of molecular, biochemical, biophysical and bioinformatics approaches, we provide evidence for an additional novel mechanism by which XN and its derivative, TXN, can inhibit diet-induced hepatic steatosis through downregulation of hepatic FA uptake and lipid storage by binding to PPARγ in the liver and effectively antagonizing its actions.

We previously demonstrated that XN and TXN ameliorated DIO in C57Bl6/J mice with no evidence of liver injury (Miranda *et al*., 2018). Using the same animal model, we confirmed the phenotypic outcomes observed in the previous study (**Fig. 2**). In this study and prior studies (Miranda *et al*., 2018), we noted a decrease in weight with treatment in the presence of similar caloric intake. Our metabolic cage data demonstrated energy expenditure increased with body mass, but a treatment effect was not identified. We hypothesize that changes in microbiota composition and bile acid metabolism which can affect nutrient and energy harvesting may explain the reduction in weight (Wahlström *et al*., 2016; Zhang *et al*., 2020) observed by treatment, but requires testing in future work. Furthermore, we demonstrated the effect of XN and TXN on the development and progression of diet-induced hepatic steatosis. Administration of XN (60 mg/kg BW) and TXN (30 mg/kg BW) significantly slowed the development and progression of hepatic steatosis during a 16-week high fat feeding. We observed less macro- and microvesicular steatosis, significantly lower liver mass to BW ratio, decreased TAG accumulation, and significantly lower steatosis scores in the XN and TXN supplemented mice compared to their untreated HFD mice (**Figs. 3B & 6**). Four pathways generally maintain hepatic lipid homeostasis: uptake of circulating lipids, *de novo* lipogenesis (DNL), FA oxidation (FAO) and lipid export in very low-density lipoproteins (VLDL). These pathways are under tight regulation by hormones, nuclear receptors, and other transcription factors (Bechmann *et al*., 2012). Long-term dysregulation of one and/or multiple processes can lead to the development of NAFLD, obesity, type 2 diabetes and other metabolic disorders.

To elucidate the mechanism of XN and TXN, we determined liver transcriptomic changes after 16 weeks of HFD feeding using RNAseq. We observed significant changes in hepatic gene expression with TXN administration (Fig. 7B). GO enrichment analysis of DEGs revealed that several biological processes were significantly downregulated by TXN treatment, including xenobiotic catabolism, FA metabolism, glucose metabolism and regulation of lipid metabolism (**Fig. 8**). Furthermore, KEGG pathway analysis of DEGs revealed multiple biological pathways were downregulated in the livers of TXN-treated mice, including biosynthesis of unsaturated FAs, glutathione metabolism, amino sugar and nucleotide sugar metabolism, glycolysis and gluconeogenesis, FA elongation, and PPAR signaling pathways, suggesting that TXN rewired global hepatic lipid metabolism (**Fig. 8**). There was a paucity of differentially expressed genes in the livers of mice supplemented with a high dose of XN (60 mg/kg BW) even at an FDR cutoff of 0.4. This discrepancy might be due to reduced levels of XN in peripheral tissues as compared with TXN as we previously observed a 12-fold lower level of XN as compared with TXN in the liver (Miranda *et al*., 2018).

To discover signature genes in the liver of mice treated with TXN, we applied a SVM classifier algorithm and extracted the most important features (genes) (**Fig. 9**). Due to the limited number of samples in this study, we did not separate the data into training and testing sets for the construction of SVM. The caveat of this is that the learning model might not generalize well. Consistent with GO analysis, three out of the eight significantly regulated genes – uncoupling protein 2 (*Ucp2*), cell death-inducing DFFA-like effector c (*Cidec*), and monoacylglycerol O-acyltransferase 1 (*Mogat1*) – are involved in lipid metabolism (**Table 1**). Notably, these genes are targets of PPARγ (Bugge *et al*., 2010; Karbowska and Kochan, 2012; Wolf Greenstein *et al*., 2017). PCR confirmed this finding (**Fig. 10**) and suggests that TXN modulates PPARγ actions.

### XN and TXN are novel natural and synthetic PPARγ antagonists

PPARγ belongs to a superfamily of nuclear receptors and just like other members, its activity requires ligand binding. PPARγ is highly expressed in white and brown adipose tissue, and to a lesser extent in the liver, kidney, and heart (Zhu *et al*., 1993; Lee and Ge, 2014). Because of its essential role in regulating adipogenesis and higher expression in the white adipose tissue, PPARγ has been a pharmacological target for drug development (Lehmann *et al*., 1995; Lefterova *et al*., 2014) in combating metabolic diseases such as insulin resistance and type 2 diabetes. Thiazolidinediones (TZDs), which include RGZ and PGZ, are the most widely investigated PPARγ agonists due to their strong insulin-sensitizing ability (Henney, 2000; Soccio, Chen and Lazar, 2014). Studies show that the main action of TZDs occurs in adipocytes (Chao *et al*., 2000). In the liver, PPARγ plays a role in hepatic lipogenesis (Sharma and Staels, 2007). Multiple clinical trials using TZDs have observed significant improvement in hepatic steatosis and inflammation (Ratziu *et al*., 2008, 2010; Sanyal *et al*., 2010), suggesting additional actions of TZDs in non-adipocytes. Interestingly, PGZ is more effective in treating fatty liver disease than RGZ, the more potent PPARγ agonist (Promrat *et al*., 2004; Ratziu *et al*., 2008, 2010), suggesting moderate binding is more effective. Unfortunate side effects of TZDs are weight gain (Fonseca, 2003), bone loss (Schwartz and Sellmeyer, 2007; Schwartz, 2008), edema and increased risk of cardiovascular complications (Nesto *et al*., 2004; Yang and Soodvilai, 2008; Bełtowski, Rachańczyk and Włodarczyk, 2013), due to over-activation of PPARγ. Thus, there is great interest in identifying “ideal” PPARγ modulators that are tissue specific with limited side effects.

An alternative strategy that aims to repress PPARγ has emerged in recent years (Ammazzalorso and Amoroso, 2019). The potential of reducing BW and improving insulin sensitivity suggests a possible clinical role of PPARγ antagonists in treating obesity and type 2 diabetes (Yamauchi *et al*., 2001; Rieusset *et al*., 2002; Nakano *et al*., 2006). Compared to agonists, researchers have identified only a few natural compounds that inhibit PPARγ, all of which have a moderate binding affinity for PPARγ receptor and can inhibit adipogenesis, obesity and/or hepatic steatosis. These include resveratrol (Calleri *et al*., 2014), 7-chloroarctinone-b isolated from the roots of *Rhaponticum uniflorum* (Li *et al*., 2009), tanshinone IIA from the roots of *Salvia miltiorrhiza* (danshen) (Gong *et al*., 2009), astaxanthin from red-colored aquatic organisms (Jia *et al*., 2012), protopanaxatriol (PPT) extracted from *Panax ginseng* roots (Zhang *et al*., 2014), foenumoside B from the herbal plant *Lysimachia foenum-graecum* (Kwak *et al*., 2016), and betulinic acid, a pentacyclic triterpene found in the bark of several plants (Brusotti *et al*., 2017; Ammazzalorso and Amoroso, 2019).

Several lines of evidence presented in this study support the hypothesis that XN and TXN are also PPARγ antagonists. First, using the 3T3-L1 cell model for PPARγ-mediated adipogenesis, we demonstrated that XN and TXN significantly and strongly suppressed RGZ induced adipocyte differentiation and adipogenesis by day 7 (**Fig. 12**). Consistent with a decrease in lipid accumulation, PPARγ target genes were also significantly downregulated in XN and TXN-treated cells (**Table 2**). The PPARγ antagonist, GW9662, did not significantly affect target gene expression of PPARγ, even though it inhibited differentiation (**Fig.12** **A2-3**). In our experiments, we used a significantly lower concentration of GW9662 than used by others that ranged from 3-25 times higher, and this difference could explain our results (Park *et al*., 2008; Kim, Nian and McIntosh, 2011; Sankella, Garg and Agarwal, 2016). Second, the PPARγ nuclear receptor competitive binding assay showed that XN and TXN have a moderate binding affinity of 1.97 µM and 1.38 µM, respectively (**Fig. 13**). Lastly, consistent with the competitive binding assay, simulated molecular docking indicated that XN and TXN can interact with the ligand binding domain of PPARγ like other known ligands and potentially form hydrogen bonds with His266, Arg280, Ser342 and Ile281, in addition to many non-bonded interactions (**Fig. 13D,E**). Moreover, the predicted binding model reveals that the interactions between XN, TXN and the PPARγ ligand binding domain resembles those observed between PPARγ and resveratrol, a dietary polyphenol that is also a PPARγ antagonist (Calleri *et al*., 2014). Our findings are consistent with XN and TXN functioning as PPARγ antagonists and now offer a mechanistic explanation for prior observations that XN impaired adipocyte differentiation (Yang et al., 2007; Mendes et al., 2008; Samuels, Shashidharamurthy and Rayalam, 2018).

One of the many side effects observed from TZD therapy is weight gain. TZDs primarily mediate their effects in adipose tissue by PPARγ activation that stimulates adipocyte differentiation and increases the efficiency of uptake of circulating non-esterified FAs (NEFA) by adipocytes (Rosen and Spiegelman, 2006). Interestingly, in this study, we observed a significant decrease in overall, sWAT, and mWAT fat mass in HXN- and TXN-treated mice (**Figs. 3A**, 5AC), yet a slight increase in the eWAT fat mass (**Fig. 5B**). Prior studies have reported that the expandability of eWAT in male mice is an indicator of metabolic health. Mouse sWAT and mWAT will continue to expand with BW, whereas eWAT expansion diminishes after mouse BW reaches about 40 g (van Beek *et al*., 2015). Our data suggest that HXN- and TXN-treated mice have capacity to expand eWAT, whereas HFD-fed untreated mice do not, which seems to direct the development of metabolic disorders. In our previous study, we demonstrated that XN and TXN accumulates primarily in the liver with significantly lower levels in the muscle (Miranda *et al*., 2018). We could not detect XN or TXN in the WAT of these mice (data not shown). The levels of XN and TXN in the liver (TXN > HXN > LXN) and the absence of both compounds in the WAT suggests that these compounds antagonize PPARγ in the liver and not in the WAT, therefore, minimizing the side effect of weight gain observed with TZDs that are PPARγ agonists.

During a long term HFD feeding, PPARγ and its target genes are upregulated to compensate for the lipid overflow in the liver. Namely, genes associated with lipid uptake and trafficking (*Lpl*, *Cd36*, *Fabp4*), TAG synthesis (*Fasn*, *Scd1*, *Mogat1*), and formation of lipid droplets for storage (*Cidec*/*Fsp27*, *Plin4*) (Supplement_File_B). The result is excessive lipid accumulation in the liver, leading to hepatic steatosis. This was observed with PPARγ overexpression in hepatocytes in *ob/ob* mice (Wolf Greenstein *et al*., 2017). We propose that TXN added to a HFD antagonizes PPARγ action in the liver potentially by physically interacting with PPARγ receptors as indicated in the molecular docking studies (**Fig. 13DE**) and; therefore, reduces PPARγ transcriptional activity and expression of the aforementioned target genes. Several *in vivo* studies support our findings. Liver-specific PPARγ deficiency protects *ob/ob* mice from hepatic steatosis (Matsusue *et al*., 2003); knockdown of *Mogat1* in the liver significantly attenuates hepatic steatosis after 12 weeks HFD feeding (Lee *et al*., 2012); and restoration of CIDEC/FSP27 in *ob/ob* liver-specific PPARγ knockout mice promotes hepatic steatosis (Matsusue *et al*., 2008). The role for PLIN4 in hepatic steatosis is limited, but it may affect TAG accumulation during HFD feeding (Griffin *et al*., 2017).

In conclusion, we demonstrated that TXN is very effective in suppressing the development and progression of diet induced hepatic steatosis in mice. TXN appears more effective in-vivo than XN perhaps due to significantly higher levels of TXN in the liver, but XN can slow progression of the condition at a higher dose. We provide evidence that XN and TXN act as novel, natural and synthetic antagonists of PPARγ that bind with a similar affinity as the agonist PGZ. Our findings support further development of XN and TXN as novel, low-cost therapeutic compounds for diet-linked hepatic steatosis with fewer negative side effects than current drugs (e.g., reduced adipose tissue expansion). Additionally, the structures of XN and TXN could serve as scaffolds for the synthesis of more effective compounds to treat NAFLD and other fatty liver diseases. Although these results are encouraging, further studies are required to clarify possible use in humans for the prevention and treatment of diet-linked hepatic steatosis.

## Materials and Methods

### Animals and diets

Studies were performed using 8-week-old SPF male C57BL/6J mice obtained from The Jackson Laboratory (Bar Harbor, ME, USA). Upon arrival, 60 mice were housed individually in ventilated cages in a controlled environment (23 ± 1°C, 50-60% relative humidity, 12 hours daylight cycle, lights off at 18:25 hours) with food and water *ad libitum*. After acclimating mice for one week on a normal-chow diet (PicoLab Rodent Diet 20, 5053, TX, USA) followed by two-weeks on a low-fat control diet (LFD, Dyets Inc., Bethlehem, PA, USA), they were randomly assigned (restricted) to five groups (n = 12/group). The sample size of 12 mice per treatment group was based on previous published studies (Miranda *et al*., 2016, 2018). The groups were fed either a LFD, HFD, HFD + 30 mg/kg BW/day XN (LXN), HFD + 60 mg/kg BW/day XN (HXN), or HFD + 30 mg/kg BW/day TXN (TXN). The sources and purity of XN and TXN were described previously (Miranda *et al*., 2018). The chemical structures of XN and TXN, a detailed diet composition and FA composition are available in **Figure 1**, **Table 3** and **Table 4**, respectively.

**Table 3.** Composition of diets*^a^*

|  | HFD | HFD+LXN | HFD+HXN | HFD + TXN | LFD |
| --- | --- | --- | --- | --- | --- |
| *Ingredient (g/100g)* |  |  |  |  |  |
| Casein | 2.58 | 2.58 | 2.58 | 2.58 | 1.89 |
| L-Cystine | 0.04 | 0.04 | 0.04 | 0.04 | 0.03 |
| Sucrose | 0.89 | 0.89 | 0.89 | 0.89 | 0.89 |
| Cornstarch | 0.00 | 0.00 | 0.00 | 0.00 | 4.02 |
| Cellulose | 0.54 | 0.54 | 0.54 | 0.54 | 0.47 |
| Dyetrose | 1.62 | 1.62 | 1.62 | 1.62 | 1.62 |
| Soybean Oil | 0.32 | 0.32 | 0.32 | 0.32 | 0.24 |
| Lard | 3.17 | 3.17 | 3.17 | 3.17 | 0.19 |
| Mineral Mix #210088 | 0.13 | 0.13 | 0.13 | 0.13 | 0.10 |
| Dicalcium Phosphate | 0.17 | 0.17 | 0.17 | 0.17 | 0.12 |
| Calcium Carbonate | 0.07 | 0.07 | 0.07 | 0.07 | 0.05 |
| Potassium Citrate H<sub>2</sub>O | 0.21 | 0.21 | 0.21 | 0.21 | 0.16 |
| Vitamin Mix #300050 | 0.13 | 0.13 | 0.13 | 0.13 | 0.10 |
| Choline Bitartrate | 0.03 | 0.03 | 0.03 | 0.03 | 0.02 |
| Test compound | 0.00 | 0.003 | 0.006 | 0.003 | 0.00 |
| OPT | 0.10 | 0.10 | 0.10 | 0.10 | 0.10 |
| *Composition (kcal%)* |  |  |  |  |  |
| Protein | 20 | 20 | 20 | 20 | 20 |
| Carbohydrates | 20 | 20 | 20 | 20 | 70 |
| Lipids | 60 | 60 | 60 | 60 | 10 |
| *Energy Density (kcal/g)* | 5.12 | 5.12 | 5.12 | 5.12 | 3.55 |

<sup>a</sup>LXN provides 30 mg/kg BW xanthohumol (XN), HXN 60 mg/kg XN, and TXN 30 mg/kg Tetrahydro-XN (TXN) per day. The test compounds were dissolved in an isotropic mixture of oleic acid: propylene glycol: Tween 80 (OPT) 0.9:1:1 by weight before incorporation into the diets. All diets were purchased from Dyets Inc., Bethlehem, PA, USA.

**Table 4.**
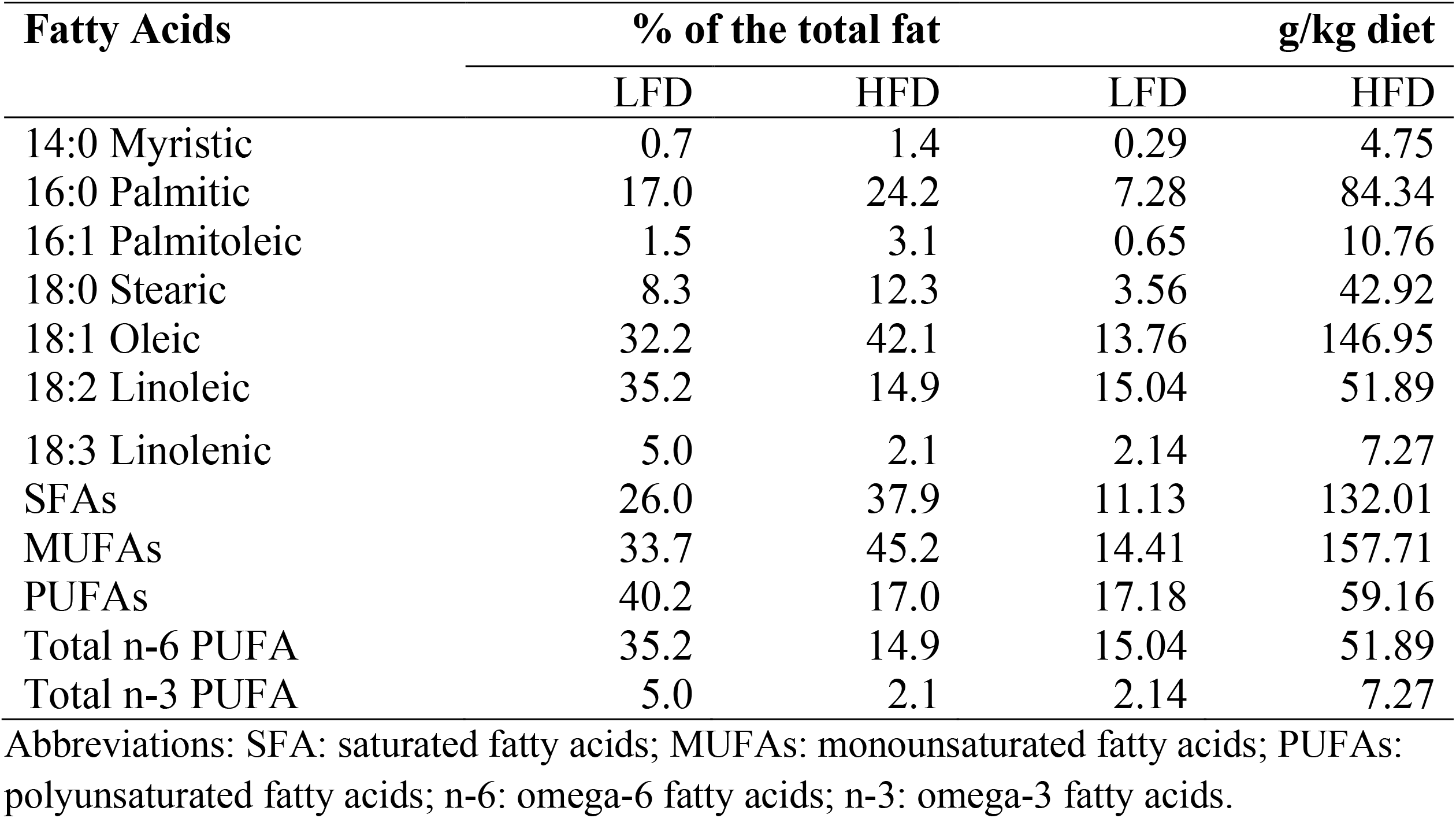
Fatty acid composition (% of the total fat) of the low-fat diet (LFD) and high-fat diet (HFD).

| Fatty Acids | % of the total fat |  | g/kg diet |  |
| --- | --- | --- | --- | --- |
|  | LFD | HFD | LFD | HFD |
| 14:0 Myristic | 0.7 | 1.4 | 0.29 | 4.75 |
| 16:0 Palmitic | 17.0 | 24.2 | 7.28 | 84.34 |
| 16:1 Palmitoleic | 1.5 | 3.1 | 0.65 | 10.76 |
| 18:0 Stearic | 8.3 | 12.3 | 3.56 | 42.92 |
| 18:1 Oleic | 32.2 | 42.1 | 13.76 | 146.95 |
| 18:2 Linoleic | 35.2 | 14.9 | 15.04 | 51.89 |
| 18:3 Linolenic | 5.0 | 2.1 | 2.14 | 7.27 |
| SFAs | 26.0 | 37.9 | 11.13 | 132.01 |
| MUFAs | 33.7 | 45.2 | 14.41 | 157.71 |
| PUFAs | 40.2 | 17.0 | 17.18 | 59.16 |
| Total n-6 PUFA | 35.2 | 14.9 | 15.04 | 51.89 |
| Total n-3 PUFA | 5.0 | 2.1 | 2.14 | 7.27 |

Abbreviations: SFA: saturated fatty acids; MUFAs: monounsaturated fatty acids; PUFAs: polyunsaturated fatty acids; n-6: omega-6 fatty acids; n-3: omega-3 fatty acids.

BW gain and food intake of individual mice were assessed once per week. Body composition was determined at the end of the feeding using a Lunar PIXImus 2 Dual Energy X-ray Absorptiometer (DXA) scan (Madison, WI, USA). After 16 weeks of feeding the control and test diets, mice were fasted for 6 h during the dark cycle, anaesthetized in chambers saturated with isoflurane and then euthanized by cardiac puncture followed with cervical dislocation. Blood was collected in syringes containing 2 IU of heparin and centrifuged to separate plasma from cells. The liver and sWAT, mWAT, and eWAT fat pads were carefully collected and weighed. To avoid batch effect due to difference in hours of fasting, mice were randomized (restricted) and treatment information was masked before sacrifice. The Institutional Animal Care and Use Committee (IACUC) at Oregon State University approved all animal work (ACUP 5053). All animal experiments were performed in accordance with the relevant guidelines and regulations as outlined in the Guide for the Care and Use of Laboratory Animals.

### Liver histology

Liver (∼100 mg) was freshly collected from mice and immediately fixed overnight in 10% neutrally buffered formalin, paraffin embedded, sectioned, and stained with hematoxylin-eosin (Veterinary Diagnostic Laboratory, Oregon State University, OR). Each slide contained two liver sections that were examined using a Leica microscope at 100× magnification. Representative images were taken at 100× magnification from the subjectively least and most severely affected areas ensuring representation of all zones of the hepatic lobule. Steatosis was objectively quantified as percent surface area occupied by lipid vacuoles using ImageJ for image analysis (NIH; imagej.nih.gov/ij/index.html) as previously published (Garcia-Jaramillo *et al*., 2019).

### Energy expenditure

Indirect calorimetry measurements were based on an open respirometer system. From week 10, mice were housed individually in Promethion^®^ Line metabolic phenotyping chambers (Sable Systems International, Las Vegas, NV, USA) and maintained on a standard 12 h light/dark cycle for three days. The system consisted of 10 metabolic cages, each equipped with food and water hoppers connected to inverted laboratory balances for food intake monitoring; both food and water were available *ad-libitum*. Spontaneous physical activity (SPA) was quantified via infrared beam breaks in X and Y axes, and included locomotion, rearing, and grooming behaviors (BXY- R, Sable Systems International). All raw data from all sensors and analyzers were stored every second. Air within the cages was sampled through micro-perforated stainless-steel sampling tubes located around the bottom of the cages, above the bedding. Ambient air was passed through the cages (2 L/min) and gases were sampled continuously for each cage, allowing the simultaneous acquisition of metabolic data every second, for all cages in the system (Lighton and Halsey, 2011). The energy expenditure was estimated from oxygen consumption (VO_2_) and carbon dioxide production (VCO_2_) rates by the Promethion system using the Weir formula (Weir, 1949).

### Liver tissue RNA extraction and library preparation

Freshly dissected liver tissue was flash frozen in liquid N_2_ and then stored at -80°C. Total RNA was isolated using the Direct-zol RNA Miniprep Plus kit as instructed (Zymo Research, Irvine, CA, USA). RNA concentrations were quantified using the Qubit^TM^ 1.0 Fluorometer and the Qubit RNA BR Assay kit (Thermo Fisher Scientific, Waltham, MA, USA). RNA purity and integrity were evaluated using a Bioanalyzer RNA 6000 Nano chip (Agilent Technologies, Santa Clara, CA, USA). Samples ranged from medium to high RNA quality (RIN 5.9-8.3), and samples with different RIN values showed similar RNA-seq qualities.

Each library was prepared with 325 ng total RNA using the Lexogen QuantSeq 3’mRNA-Seq Library Prep Kit-FWD for Illumina sequencing according to the manufacturer’s instructions (Lexogen GmbH, Vienna, Austria). Briefly, library preparation was started by oligo(dT) priming, with primers already containing the Illumina-compatible linker sequence for Read 2. After first-strand synthesis, the RNA was removed before random primers that contained the corresponding Illumina-compatible linker sequence for Read 1 initiated the second-strand synthesis. Second strand synthesis was followed by a magnetic bead-based purification step. The libraries were PCR amplified introducing sequences required for cluster generation and i7 and i5 dual indices (Lexogen i7 6 nt Index Set and Lexogen i5 6 nt Unique Dual Indexing Add-on Kit) for 16-20 PCR cycles with the optimal number predetermined by qPCR with the PCR Add-on Kit for Illumina (Lexogen GmbH). After a second magnetic bead-based purification, libraries were quantified using the Qubit dsDNA HS Assay Kit (Thermo Fisher Scientific) and sized using an Agilent High Sensitive D5000 Screen Tape (Agilent Technologies) to determine molarity. Equal molar amounts of the libraries were multiplexed and then sequenced on an Illumina Hiseq3000 platform (Illumina, San Diego, CA, USA) at the Center for Genome Research and Biocomputing, Oregon State University using single-end sequencing with 100-bp reads. Approximately 6.6 million reads were obtained per liver sample.

### Sequence alignment and gene counts

Adaptors and low quality tails were trimmed and ribosomal rRNA contaminations were removed using BBDuk from the BBTools toolset (Bushnell, 2014). As recommended by the manufacturer (Lexogen GmbH), a Phred score of 10 and a read length of 20 were used as the minimum cutoff prior to data analysis (https://www.lexogen.com/quantseq-data-analysis/). Using a splice-aware aligner STAR (Dobin *et al*., 2013) (version 37.95), cleaned reads were then mapped against the GRCm38 primary assembly of the *Mus musculus* genome (version mm10, M22 release) (ftp://ftp.ebi.ac.uk/pub/databases/gencode/Gencode_mouse/release_M22/GRCm38.primary_assenbly.genome.fa.gz) with the annotation file of the same version (ftp://ftp.ebi.ac.uk/pub/databases/gencode/Gencode_mouse/release_M22/gencode.vM22.annotation.gtf.gz), both from the GENCODE project (Frankish *et al*., 2019). On average, over 81% of the reads were uniquely mapped for each sample. Downstream analyses were based on uniquely aligned reads.

To generate count matrices from bam files, the summarizeOverlaps function from the GenomicAlignments package (v1.26.0) was used (Lawrence *et al*., 2013). The location of the exons for each gene was obtained from a transcript database (TxDb) using the makeTxDbFromGFF function from the GenomicFeatures package (version 1.42.1), with a pre-scanned GTF file used in the mapping step. Genes were then annotated with the R package Mus musculus (version 1.3.1) (Team, 2016).

### Identification of differentially expressed genes (DEGs)

R package edgeR (version 3.26.8) was used to detect differential change in gene expression among mice on different diets (Robinson, McCarthy and Smyth, 2010). Genes expressed in at least nine samples were retained using the filterByExpr function in edgeR. Unannotated genes, pseudogenes and ribosomal RNA genes were also removed from downstream analyses. Gene counts were then normalized with the default TMM (trimmed mean of M-values) method (Robinson and Oshlack, 2010) provided by edgeR. To account for both biological and technical variability, an overdispersed Poisson model and an Empirical Bayes method were used to moderate the degree of overdispersion across transcripts. Genes with a false discovery rate (FDR) threshold < 0.4 were used for heatmap and volcano plot analyses, whereas genes with an FDR threshold < 0.05 were used in gene ontology and pathway enrichment analysis.

### Gene ontology and pathway enrichment analyses

Gene ontology (GO) and KEGG pathway enrichment analysis was conducted using Enrichr (http://amp.pharm.mssm.edu/Enrichr) (Chen *et al*., 2013; Kuleshov *et al*., 2016). Genes with an FDR threshold < 0.05 were analyzed with GO biological process 2018 and KEGG 2019 Mouse databases. Full tables can be found in the supplementary material (Supplement_File_A).

### Classification of RNA-seq data

Gene selection and normalization were performed using the R package DaMiRseq 1.2.0 (Chiesa, Colombo and Piacentini, 2018). To distinguish TXN-fed samples from HFD control samples, we used a correlation cutoff of 0.4 for the partial least-squares feature selection (FSelect), and the default correlation coefficient for the redundant feature removal (FReduct).

### Cell culture

Murine 3T3-L1 pre-adipocytes were obtained from ATCC (Rockville, MD, USA). Prior to treatments, cells were maintained in basic media, which consisted of high glucose DMEM supplemented with 1% penicillin-streptomycin and 10% heat-inactivated FBS (Hyclone, Logan, UT, USA). The cells were allowed to reach full confluence for 2 days. Differentiation was induced by the addition of 0.5 µM IBMX (Sigma-Aldrich, St. Louis, MO, USA), 0.25 µM dexamethasone (Sigma-Aldrich) and 10 µg/ml insulin (Sigma-Aldrich) plus the addition of treatment compounds XN or TXN. After 48 h, media was removed and fresh DMEM was replenished for continuing differentiation. To observe XN and TXN’s effects on 3T3-L1 adipocyte differentiation, different concentrations were selected based on dose-response experiments to identify the dose that maximized effectiveness while minimizing cell toxicity.

### MTT cell viability assay

For cell viability experiments using the MTT assay, 3T3-L1 fibroblasts were seeded in 96-well plates at a density of 15,000 cells per well in 200 µl of DMEM medium supplemented with 10% FBS, 1% glutamine, 1 mM of sodium pyruvate, 100 units/mL penicillin, and 100 µg/mL streptomycin. After incubating 48 h with various concentrations of XN or TXN at 37⁰C in 5% CO_2_ atmosphere, the culture medium was removed and a solution of MTT [3-(4,5- dimethylthiazol-2-yl)-2,5-diphenyltetrazolium bromide], 0.5 mg/mL in complete culture medium, was added to each well. The cells were incubated with MTT for 3 h at 37⁰C and then the MTT medium was removed before adding acidified isopropanol to each well. The cells were shaken for 10 min in an orbital shaker before reading the absorbance at 570 nm using a Microplate Reader (SpectraMax 190, Molecular Devices, Sunnyvale, CA, USA). Cell viability of compound-treated cells was calculated as percent absorbance of vehicle-treated control cells.

### Oil red O staining

Cells were washed twice with phosphate-buffer saline (PBS) and then fixed with 10% formalin for 30 min. Cells were then washed with ddH_2_O followed by 60% isopropanol. A 0.4% stock solution of Oil Red O (Sigma-Aldrich) in isopropanol was diluted 3:2 (Oil red O:ddH_2_O) for a working solution. To determine intracellular lipid accumulation, fixed cells were incubated for 30 – 60 min at room temperature on a rocker with the Oil red O working solution. After incubation, cells were washed with ddH_2_O and imaged using microscopy.

### Adipocyte gene expression by RT-qPCR

Total RNA was isolated as described above, dissolved in RNase-free water and stored at -80 °C. For RT-PCR experiments, cells were grown in 6-well plates and treated with XN and TXN at 25 µM concentration and differentiation medium after confluence for 2 days. Gene expression was measured from cells at 7 d post treatment. RNA (0.25 µg) was converted to cDNA using iScript reverse transcriptase and random hexamer primers (Bio-Rad Laboratories), according to the manufacturer’s recommendations. PCRs were set up as described previously (Gombart, Borregaard and Koeffler, 2005). All the threshold cycle number (CT) were normalized to Ywhaz reference gene. PrimeTime^®^ Std qPCR assays were purchased from IDT (**Table 5**). ΔCT = CT(target gene) – CT(reference gene). ΔΔCT = ΔCT(treated sample) – ΔCT(untreated sample/control average). Statistics were done on ΔΔCT values.

**Table 5.** Primer probe information

| Gene Name | IDT Assay Name | RefSeq Number |
| --- | --- | --- |
| *Cd36* | Mm.PT.58.12375764 | NM_007643 |
| *Cidec/Fsp27* | Mm.PT.58.6462335 | NM_178373 |
| *Fabp4* | Mm.PT.58.43866459 | NM_024406 |
| *Fgf21* | Mm.PT.58.29365871.g | NM_020013 |
| *Il6* | Mm.PT.58.10005566 | NM_031168 |
| *Lpl* | Mm.PT.58.46006099 | NM_008509 |
| *Mogat1* | Mm.PT.58.41635461 | NM_026713 |
| *Pparγ2* | Mm.PT.58.31161924 | NM_011146 |
| *Ppargc1α* | Mm.PT.58.28716430 | NR_027710 |
| *Plin4* | Mm.PT.58.43717773 | NM_020568 |
| *Scd1* | Mm.PT.58.8351960 | NM_009127 |
| *Srebf1* | Mm.PT.58.8508227 | NM_011480 |

### Time-resolved fluorescence resonance energy transfer (TR-FRET)

To determine the binding affinity of XN and TXN to PPARγ, a Lanthascreen^TM^ TR-FRET PPARγ competitive binding assay was performed by Thermo Fisher Scientific as described (cite manual). A terbium-labeled anti-GST antibody binds to a GST-PPARγ-ligand binding domain fusion protein in which the LBD is occupied by a fluorescent pan- PPAR ligand (Fluormone^TM^ Pan-PPAR Green). Energy transfer from the antibody to the ligand occurs and a high TR-FRET ratio (emission signal at 520 nm/495 nm) is detected. When a test compound displaces the ligand from PPARγ-LBD, a decrease in the FRET signal occurs and a lower TR-FRET ratio is detected (Invitrogen Corporation, 2008). For each compound (XN, TXN or oleic acid) a 10-point serial dilution (250,000 to 12.5 nM) was tested. Binding curves were generated by plotting percent displacement versus log concentration (nM), and IC_50_ values were determined using a sigmoidal dose response (variable slope).

### Molecular docking simulations for XN and TXN into the PPARγ ligand-binding domain

To estimate the binding mode of XN and TXN to PPARγ, molecular docking simulations were performed using AutoDock Vina (Trott and Olson, 2010). Structural models of XN and TXN were built using OpenBabel to convert the isometric SMILES descriptor for XN to a PDB formatted file, which was subsequently modified using PyMOL (The PyMOL Molecular Graphics System, Version 1.7.4.5, Schrödinger, LLC) to obtain a PDB file for TXN. The solved structure of PPARγ bound to the antagonist resveratrol (PDB ID: 4JAZ) was used as the receptor model. The PDBQT files for the receptor and the resveratrol, XN, TXN, and oleic acid ligands were generated using MGLTools-1.5.7rc1 (Morris *et al*., 2009). The PPARγ receptor was kept rigid during all docking experiments, and the center and size (20 x 20 x 20 Å^3^) of the docking box was positioned to cover the entire ligand-binding site of PPARγ. All rotatable torsion angles in the ligand models were allowed to be active during the docking simulations. Twenty docking poses were generated for each simulation, and the conformation with the lowest docking energy was chosen as being representative.

### Statistical analysis

Analysis of variance procedures for continuous data and Fisher’s exact test for binary data were used for statistical comparisons. P-values of orthogonal *a priori* comparisons of the HFD control group versus each of the supplement groups are shown in the corresponding tables and figures. Additional details of statistical analyses are described in the corresponding figure legends.

## Acknowledgements

We thank Jamie Pennington, Scott Leonard and Dr. Wenbin Wu for their assistance, Dr. Edward Davis for bioinformatics support and Anne-Marie Girard-Pohjanpelto, Mark Dasenko, Dr. Brent Kronmiller and Matthew Peterson at the Center of Genome Research and Bioinformatics at Oregon State University (OSU) for their assistance with RNA-sequencing. We thank Drs. Russ Turner and Urszula Iwaniec at the School of Biological and Population Health Sciences at OSU for use of the Lunar PIXImus 2 Dual Energy X-ray Absorptiometer (DXA) instrument. The National Institutes of Health (NIH grants 5R01AT009168 to A.F.G., C.S.M., and J.F.S. and 1S10RR027878 to J.F.S.), the Linus Pauling Institute (LPI), the OSU College of Pharmacy, Hopsteiner, Inc., New York, and the OSU Foundation Buhler-Wang Research Fund supported this research. The Marion T. Tsefalas Graduate Fellowship from the LPI, the ZRT Laboratory Fund for the LPI, and the Charley Helen, Nutrition Science and Margy J. Woodburn Fellowships from the School of Biological and Population Health Sciences at OSU supported Y.Z.

---

Figure 2—source data 1

Source files.

This zip archive contains the following:

1) One Comma Separated Values file named “phenome_feeding.csv” contains food intake and weight entries.

2) One Excel workbook named “2019TXN_repeated_measures_YZGB.xlsx” contains repeated measures analyses.

3) The Jupyter Notebook contains scripts used for statistical analysis and generation of Fig. 2.

Figure 3—source data 1 Source files.

This zip archive contains the following:

1) One Comma Separated Values file named “metabolicGasExchange.csv” contains metabolic cage gas exchange data.

2) One Comma Separated Values file named “fig3_table.csv” contains phenotypic data directly pertaining to Fig. 3.

3) A Jupyter Notebook file contains scripts used for statistical analysis and generation of Fig. 3.

4) An R script file “ggplotRegression.R”

5) A folder named “Fig3Sup1” containing Figure 3—figure supplement 1.

a. One Comma Separated Values file named “metabolicGasExchange.csv” contains metabolic cage gas exchange data.

b. One Comma Separated Values file named “supplement1Table.csv” contains phenotypic data directly pertaining to Figure 3—figure supplement 1.

c. An R script file “ggplotRegression.R.

d. A Jupyter Notebook file contains scripts used for statistical analysis and generation of figure supplement 1.

e. A pdf file named “fig3Sup1.pdf”.

f. A word document named “fig3Sup1.docx” containing the figure and figure legend.

6) A folder named “Fig3Sup2” containing Figure 3—figure supplement 2.

a. One Comma Separated Values file named “supplement2Table.csv” contains phenotypic data directly pertaining to Figure 3—figure supplement 2.

b. An R script file “ggplotRegression.R.

c. A Jupyter Notebook file contains scripts used for statistical analysis and generation of figure supplement 2.

d. A pdf file named “fig3Sup2.pdf”.

e. A word document named “fig3Sup2.docx” containing the figure and figure legend.

Figure 4—source data 1 Source files.

This zip archive contains the following:

1) One Comma Separated Values file named “fig4_table.csv” phenotypic data directly pertaining to Fig. 4.

2) An R script file “ggplotRegression.R”.

3) A Jupyter Notebook file contains scripts used for statistical analysis and generation of Fig. 4.

Figure 5—source data 1 Source files.

This zip archive contains the following:

1) One Comma Separated Values file named “fig5_table.csv” phenotypic data directly pertaining to Fig. 5.

2) An R script file “ggplotRegression.R”.

3) A Jupyter Notebook file contains scripts used for statistical analysis and generation of Fig. 5.

Figure 6-source data 1

Source files for histology data.

A folder called “TXN prevents HFD induced liver steatosis in mice” containing histology images in TIFF format (n = 59), used for histology scoring and Excel spreadsheet with scores and sample IDs. https://doi.org.

Figure 6-source data 2

This zip archive contains the following:

1) One Comma Separated Values file named “fig6_table.csv” phenotypic data directly pertaining to Fig. 6.

2) A Jupyter Notebook file contains scripts used for statistical analysis and generation of Fig. 6

3) Two pdf files named “B.pdf” and “C.pdf”.

Figure 7—source data 1 Source files.

This zip archive contains the following:

1) A Jupyter Notebook file contains scripts used for statistical analysis and generation of Fig. 7.

2) A R object file in Rds format named “y_keep.rds”.

3) An R script used to generate the ‘y_keep.rds’ file.

Figure 8—source data 1 Source files.

This zip archive contains the following:

1) A folder named “raw”, containing five Excel workbooks

a. “DEGs_TXN_vs_HFD.xlsx”

b. “DOWN-GO_Biological_Process_2018.xlsx”

c. “UP-GO_Biological_Process_2018.xlsx”

d. “DOWN-KEGG_2019_Mouse.xlsx”

e. “UP-KEGG_2019_Mouse.xlsx”

2) A folder named “processed”, containing two Comma Separated Values files:

a. “BPTerms.csv”

b. “KEGGterms.csv”

3) A Jupyter Notebook file contains scripts used for statistical analysis and generation of Fig. 8.

4) A pdf file named “txnHFDGO.pdf”.

Figure 9—source data 1 Source files.

This zip archive contains the following:

1) A Comma Separated Values file named “colData_hftxn.csv” contains experiment metadata.

2) A Comma Separated Values file named “countMatrix_hftxn.csv” contains raw counts in HFD and HFD+TXN groups.

3) A tab-delimited text file named “dfimportance_hftxn_lgcpm.txt”.

4) A Jupyter Notebook file contains scripts used for statistical analysis and generation of Fig. 9.

5) A pdf file named “leftPanel.pdf”.

6) A pdf file named “rightPanel.pdf”.

7) A PowerPoint file named “fig9.pptx”.

8) A pdf file named “fig9.pdf”.

Figure 10—source data 1 Source files.

This zip archive contains the following:

1) A Comma Separated Values file named “fig10_table.csv” phenotypic data directly pertaining to Fig. 10.

2) A Excel workbook named “PCR_lv_raw.xlsx” contains raw PCR cycle number data, and the calculation of fold change.

3) A Jupyter Notebook file contains scripts used for statistical analysis and generation of Fig. 10.

4) A pdf file named “fig10.pdf”.

Figure 13—source data 12

Source files: an Excel file named “SSBN12209_57828_10-point Titration_Inhibition_Results.xls” containing results from ThermoFisher PPARγ nuclear receptor competitive binding assay.

Table 1—source data 1 Source files.

This zip archive contains the following:

1) An Excel workbook named “DEG_HFD_vs_TXN.xlsx” contains all differentially expressed genes identified. Genes listed in the table were highlighted in yellow in the Excel workbook.

Table 2—source data 1 Source files.

This zip archive contains the following:

2) An Excel workbook named “7days.xlsx” contains raw PCR cycle numbers, fold change, log(2)fold change, p values, and how these are calculated.

**Figure 3-supplement 1.**
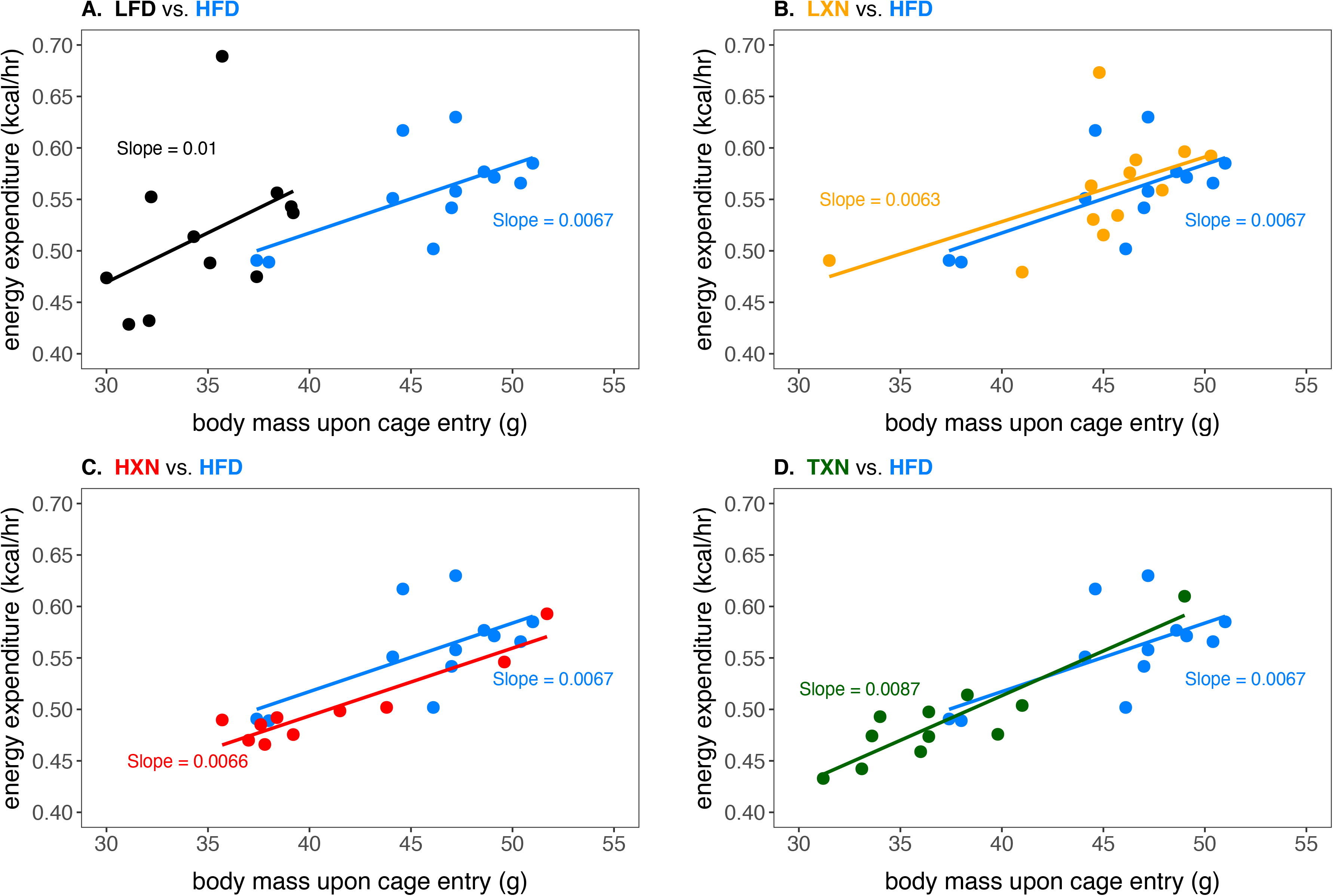
Relationship of body mass and energy expenditure between (A) LFD and HFD; (B) LXN and HFD; (C) HXN and HFD; (D) TXN and HFD. Energy expenditure was measured between weeks 10 to 14. Data was analyzed using analysis of covariance (ANCOVA) of body mass upon entry into the cages and diet. No statistically significant effect from treatments was detected. HFD data are from the same group of mice and are displayed as a reference on all four panels. Source files of data used for the analysis are available in the Figure supplement 1—source data 1.

**Figure 3-supplement 2.**
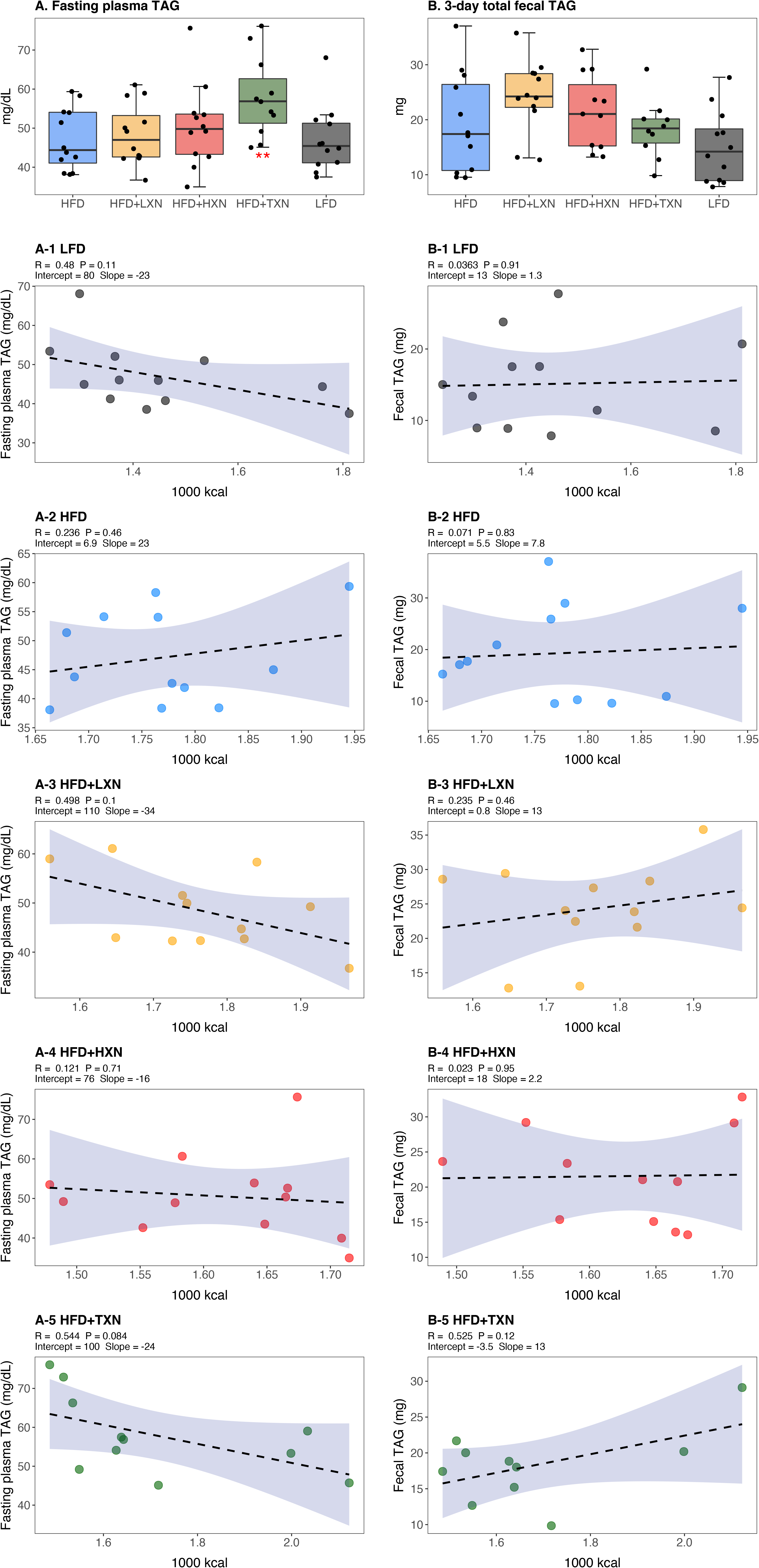
The effect of diet and intervention on fasting plasma and fecal TAG levels. Mice were fed either a LFD (black, n = 12), a HFD (blue, n = 12), HFD+LXN (yellow, n = 12), HFD+HXN (red, n = 12), or HFD+TXN (green, n = 11) for 16 weeks. (A) Fasting plasma TAG levels are expressed as quartiles. (A-1) Relationship between fasting plasma TAG and total caloric intake over 16 weeks of feeding for LFD group; (A-2) for HFD group; (A-3) for HFD+LXN group; (A-4) for HFD+HXN group; (A-5) for HFD+TXN group (B) 3-day total fecal triglyceride (TAG) are expressed as quartiles. (B-1) Relationship between 3-day fecal TAG and total caloric intake over 16 weeks of feeding for LFD group; (B-2) for HFD group; (B-3) for HFD+LXN group; (B-4) for HFD+HXN group; (B-5) for HFD+TXN group. Pre-planned general linear model with contrasts were used to calculate *p*-values in A, B and C. \**p* < 0.05, \*\**p* < 0.01, \*\*\**p* < 0.001. Linear regression analyses of total calories versus fasting plasma TAG (A1-5) or total fecal TAG (B1-5) in mice were done using lm function of stats package version 3.6.2 in R. Blue shading represents 95% CI of the regression line. Absolute value of R, p-value, intercept, and slope for the regression are reported above each corresponding panel.

