## Supplementary material for "TXN, a Xanthohumol Derivative, Attenuates High-Fat Diet Induced Hepatic Steatosis by Antagonizing PPARγ": Figure 2 source data: fig2.ipynb

```
# %% [code]
suppressPackageStartupMessages(library(ggplot2))
suppressPackageStartupMessages(library(dplyr))
suppressPackageStartupMessages(library(tidyverse))
suppressPackageStartupMessages(library(emmeans))
suppressPackageStartupMessages(library(ggpubr))
suppressPackageStartupMessages(library(export))
suppressPackageStartupMessages(library(gridExtra))
R.Version()

# %% [code]
feeding <- read.csv("phenome_feeding.csv", as.is = TRUE); head(feeding)

# %% [code]
feeding <- feeding %>% dplyr::mutate(
    vars = factor(Treatment,levels = c("HFD", "HFD+LXN", "HFD+HXN","HFD+TXN", "LFD"))
)
feeding$Date <- as.Date(feeding$Date, "%m/%d/%y")
feeding$Treatment <- as.factor(feeding$Treatment)
feeding$Week <- as.factor(feeding$Week)
feeding$ID <- ordered(feeding$ID)
feeding$ID <- as.factor(feeding$ID)

# %% [code]
weight_int <- filter(feeding, Date == "2019-02-04")
total_gained <- filter(feeding, Date == "2019-05-28")

# %% [code]
total_food <- feeding %>% group_by(ID, Treatment) %>% summarise(total = sum(Food_Intake_weekly), .groups = 'drop')
total_food <- total_food %>% dplyr::mutate(
    vars = factor(Treatment, levels = c("HFD", "HFD+LXN", "HFD+HXN", "HFD+TXN", "LFD"))
)

# %% [code]
head(total_food)

# %% [code]
total_cal <- feeding %>% group_by(ID, Treatment) %>% summarise(total = sum(Calories_weekly), .groups = 'drop')
total_cal <- total_cal %>% dplyr::mutate(
    vars = factor(Treatment, levels = c("HFD", "HFD+LXN", "HFD+HXN", "HFD+TXN", "LFD"))
); head(total_cal)

# %% [code]
mycolor_solid <- c("#0080ff", "#ffa500", "red", "darkgreen", "black")
mycolor_alpha <- c("#8AB5F9", "#F8CB89", "#F08581", "#87A57E", "#797979")
lines <- c("solid", "solid", "solid", "solid", "dashed")
shapes <- c(24, 4, 15, 25, 1)

# %% [code]
theme_legend_free = function(){
	theme(axis.title.y = element_text(size = 15, 
	margin = margin(t=0, r=5, b=0, l=10)),
	axis.text.y = element_text(size = 13),
	axis.text.x = element_blank(),
	axis.ticks.x = element_blank(),
	legend.position = "none",
	plot.margin = unit(c(0.5,0.5,0,0.5), "cm"),
	panel.grid = element_blank(),
	panel.grid.major = element_blank(),
	panel.grid.minor = element_blank())
}


# %% [code]
F1A <- ggline(feeding, x = "Week", y = "Weight_gain_percent", group = "vars",
              add = "mean_se", color = "vars", palette = mycolor_solid,
              ylab = "%", 
              linetype = "vars", shape = "vars", 
              point.size = 2,
              ggtheme = theme_bw())

A <- F1A + scale_linetype_manual(values=c( "solid", "solid", "solid", "solid", "dashed")) + 
            scale_shape_manual(values = shapes) +
            ggtitle("Weekly Weight Gain (%)") +
            theme(axis.title.y = element_text(size = 15, margin = margin(t=0, r=10, b=0, l=10)),
            axis.text.y = element_text(size = 15),
            axis.text.x = element_text(size = 18),
            legend.position = "top",
            legend.title = element_blank(),
            legend.margin = margin(c(0,0,0,0)),
            plot.margin = unit(c(0.5,0.5,0,0.5), "cm"),
            panel.grid = element_blank())

A <- A + theme(legend.position = c(0.1, 0.6), legend.background = element_rect())

# %% [code]
B <- ggplot(data = total_gained, aes(x = vars, y = Weight_gain_percent, fill = vars)) +
            stat_boxplot(geom = 'errorbar', linetype = 1, width = 0.2) +
            geom_boxplot(outlier.shape = NA) +
            scale_fill_manual(values = mycolor_alpha) +
            geom_jitter(shape = 20, size = 1, width = 0.2, height = 0.1) +
            xlab(" ") + ylab("%") +
            ggtitle("Cumulative Weight Gained") +
            theme_bw() +
            theme_legend_free()

label.df <- data.frame(vars = c("HFD+TXN", "LFD"),
                       Weight_gain_percent = c(25, 25))
B <- B + geom_text(data = label.df, label = c("***", "***"), color='red', size = 6)

# %% [code]
food <- ggline(feeding, x = "Week", y = "Food_Intake_weekly", group = "vars",
               add = "mean_se", color = "vars", palette = mycolor_solid,
               ylab = "g", 
               linetype = "vars", shape = "vars",
               point.size = 2,
               ggtheme = theme_bw())

C <- food + scale_shape_manual(values = shapes) +
            scale_linetype_manual(values=c("solid", "solid", "solid", "solid", "dashed")) + 
            ggtitle("Weekly Food Intake") +
            theme(axis.title.y = element_text(size = 15, margin = margin(t=0, r=10, b=0, l=10)),
            axis.text.y = element_text(size = 15),
            axis.text.x = element_text(size = 18),
            legend.position = "none",
            plot.margin = unit(c(0.5,0.5,0,0.5), "cm"),
            panel.grid = element_blank())

# %% [code]
D <- ggplot(data = total_cal, aes(x = vars, y = total, fill = vars)) +
	stat_boxplot(geom = 'errorbar', linetype = 1, width = 0.2) +
	geom_boxplot(outlier.shape = NA) +
	scale_fill_manual(values = mycolor_alpha) +
	geom_jitter(shape = 20, size = 1, width = 0.2, height = 0.1) +
	xlab("") + ylab("kcal") +
	ggtitle("Cumulative Calorie Intake") +
	theme_bw() +
	theme_legend_free()

label.df <- data.frame(vars = c("HFD+HXN", "LFD"),
                       total = c(1800, 1820))
D <- D + geom_text(data = label.df, label = c("*", "***"), color='red', size = 6)

# %% [code]
par1 <- ggarrange(A, B, align = "hv", 
                  widths = c(2.5,1),
                  labels = c("A", "B"),
                  font.label = list(size = 18, color = "black"),
                  nrow = 1, ncol = 2)

# %% [code]
par2 <- ggarrange(C, D, align = "hv", 
                  widths = c(2.5,1),
                  labels = c("C", "D"),
                  font.label = list(size = 18, color = "black"),
                  nrow = 1, ncol = 2,
                  common.legend = TRUE, legend="none")

# %% [code]
partA <- ggarrange(par1, par2, align = "hv",
                  heights = c(1.5,1.5),
                  ncol = 1, nrow = 2,
                  common.legend = TRUE, legend="bottom")

# %% [code]
#options(repr.plot.width = 14, repr.plot.height = 8, repr.plot.res = 150)
#partA

# %% [code]
#graph2pdf(partA, file = "figure2.pdf", width = 14, height = 8)

# %% [code]
set.seed(12345)
Contrasts <- list(
	'LXN vs HFD' = c(-1, 1, 0, 0, 0),
	'HXN vs HFD' = c(-1, 0, 1, 0, 0),
	'TXN vs HFD' = c(-1, 0, 0, 1, 0),
	'LFD vs HFD' = c(-1, 0, 0, 0, 1),
	'TXN vs LFD' = c(0, 0, 0, 1, -1))

# %% [code]
total_gained %>% group_by(vars) %>% summarise(
        mean_wt_g_pct = sprintf("%0.2f", 
                                mean(Weight_gain_percent), .groups = 'drop'), 
        median_wt_g_pct = sprintf("%0.2f", 
                                  median(Weight_gain_percent), .groups = 'drop'),
        se_wt_g_pct = sprintf("%0.2f", 
                              sd(Weight_gain_percent)/sqrt(n()), .groups = 'drop'),
        mean_wt = sprintf("%0.2f",
                          mean(Weight), .groups = 'drop'), 
        median_wt = sprintf("%0.2f",
                            median(Weight), .groups = 'drop'),
        se_wt = sprintf("%0.2f", 
                        sd(Weight)/sqrt(n())), .groups = 'drop') 

# %% [code]
model_wg <- lm(Weight_gain_percent ~ vars, data = total_gained)
leastsquare_wg <- lsmeans(model_wg, "vars")
output_wg <- contrast(leastsquare_wg, Contrasts, adjust = "none")

print(output_wg)

# %% [code]
total_food %>% group_by(vars) %>% summarise(
        mean_fd = sprintf("%0.2f",
                          mean(total), .groups = 'drop'),
        median_fd = sprintf("%0.2f",
                            median(total), .groups = 'drop'),
        se_fd = sprintf("%0.2f",
                        sd(total)/sqrt(n())), .groups = 'drop')

# %% [code]
model_fd <- lm(total ~ vars, data = total_food)
leastsquare_fd <- lsmeans(model_fd, "vars")
output_fd <- contrast(leastsquare_fd, Contrasts, adjust = "none")

print(output_fd)

# %% [code]
total_cal %>% group_by(vars) %>% summarise(
    mean_cal = sprintf("%0.2f",
                       mean(total), .groups = 'drop'),
    median_cal = sprintf("%0.2f",
                         median(total), .groups = 'drop'),
    se_cal = sprintf("%0.2f",
                     sd(total)/sqrt(n())), .groups = 'drop')

# %% [code]
model_cal <- lm(total ~ vars, data = total_cal)
leastsquare_cal <- lsmeans(model_cal, "vars")
output_cal <- contrast(leastsquare_cal, Contrasts, adjust = "none")

print(output_cal)

# %% [code]
```
