## Supplementary material for "TXN, a Xanthohumol Derivative, Attenuates High-Fat Diet Induced Hepatic Steatosis by Antagonizing PPARγ": Figure 3 source data: fig3Sup1.docx

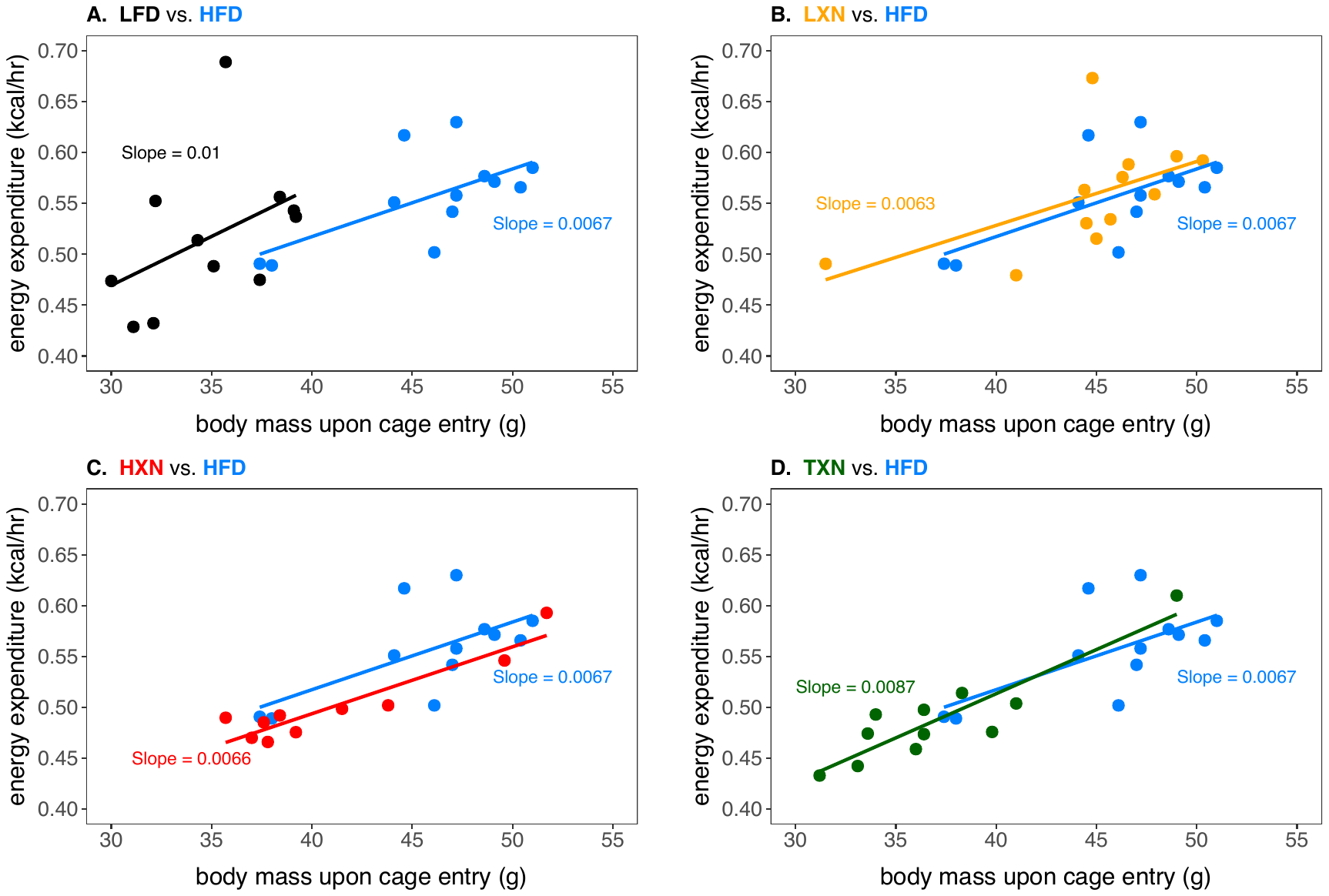

Figure supplement 1. Relationship of body mass and energy expenditure between (A) LFD and HFD; (B) LXN and HFD; (C) HXN and HFD; (D) TXN and HFD. Energy expenditure was measured between weeks 10 to 14. Data was analyzed using analysis of covariance (ANCOVA) of body mass upon entry into the cages and diet. No statistically significant effect from treatments was detected. HFD data are from the same group of mice and are displayed as a reference on all four panels. Source files of data used for the analysis are available in the Figure supplement 1—source data 1.
