## Supplementary material for "TXN, a Xanthohumol Derivative, Attenuates High-Fat Diet Induced Hepatic Steatosis by Antagonizing PPARγ": Figure 3 source data: fig3Sup1.ipynb

```
# %% [code]
# import packages

# %% [code]
suppressPackageStartupMessages(library(dplyr))
suppressPackageStartupMessages(library(ggplot2))
suppressPackageStartupMessages(library(lsmeans))
suppressPackageStartupMessages(library(patchwork))
suppressPackageStartupMessages(library(export))
suppressPackageStartupMessages(library(ggtext))
R.Version()

# %% [code]
# import data

# %% [code]
profile <- read.csv(file = "cageGasExchange.csv", as.is = TRUE)
profile$Diet <- as.factor(profile$Diet)
profile$Diet <- factor(profile$Diet, levels = c("HFD", "HFD+LXN", "HFD+HXN", "HFD+TXN", "LFD"))
profile$MouseID <- ordered(profile$MouseID)
profile$MouseID <- as.factor(profile$MouseID); head(profile)

# %% [code]
data <- read.csv("supplement1Table.csv", header = T); 
data$vars <- factor(data$Treatment, levels = c("LFD", "HFD", "HFD+LXN", "HFD+HXN", "HFD+TXN"))
data$vars2 <- factor(data$Treatment, levels = c("HFD", "HFD+LXN", "HFD+HXN", "HFD+TXN", "LFD"))

txn <- data %>% filter(vars2 == "HFD+TXN"); hfd <- data %>% filter(vars2 == "HFD");
lxn <- data %>% filter(vars2 == "HFD+LXN"); hxn <- data %>% filter(vars2 == "HFD+HXN");
lfd <- data %>% filter(vars2 == "LFD")

# %% [code]
source("ggplotRegression.R")

# %% [code]
theme_set(theme_bw())

# %% [code]
theme_legend1 = function(){
	theme(axis.title.y = element_text(size = 15, 
	margin = margin(t=0, r=5, b=0, l=5)),
	axis.text.y = element_text(size = 13),
	axis.text.x = element_text(size = 13),
	axis.title.x = element_text(size = 15,
                               margin = margin(t=10, r=5, b=0, l=0)),
    #plot.title = element_text(size = 15, face = "bold"),
          plot.title = element_markdown(),
	legend.title = element_text(colour = "white"),
	legend.text = element_text(size = 16),
	legend.position = c(0.8, 0.2),
	plot.margin = unit(c(0.5,0.5,0,0.5), "cm"),
	panel.grid = element_blank())
}

# %% [markdown]
# regression plots

# %% [code]
dataJ <- data %>% filter(Treatment == "HFD" | Treatment == 'LFD')
dataJ$color[dataJ$Treatment == 'HFD'] = '#0080ff'
dataJ$color[dataJ$Treatment == 'LFD'] = 'black'

# %% [markdown]
## HFD vs LFD

# %% [markdown]
### calculate R and slope

# %% [code]
dataJ1 <- data %>% filter(Treatment == "HFD")
dataJ2 <- data %>% filter(Treatment == "LFD")
options(repr.plot.width = 4, repr.plot.height = 4, repr.plot.res = 200)
ggplotRegression(lm(avgEE2 ~ bw_cage, data = dataJ1), "black") + labs(title = "HFD")
ggplotRegression(lm(avgEE2 ~ bw_cage, data = dataJ2), "black") + labs(title = "LFD")

# %% [markdown]
### put together

# %% [code]
options(repr.plot.width = 6, repr.plot.height = 4, repr.plot.res = 200)
parA <- ggplot(dataJ,aes(y = avgEE2, x = bw_cage, colour = color)) +
   geom_point(size=3) + geom_smooth(method = "lm", fill = NA) + 
    scale_y_continuous(limits = c(0.4, 0.7), breaks = seq(0.4, 0.7, 0.05)) +
    scale_x_continuous(limits = c(30, 55), breaks = seq(30, 55, 5)) +
    scale_color_identity() +
    ggtitle("**A.** <span style='color:black'>**LFD**</span> vs. <span style='color:#0080ff'>**HFD**</span>") + 
    ylab('energy expenditure (kcal/hr)') + 
    xlab('body mass upon cage entry (g)') + 
    annotate("text", x = 52, y = 0.53, label = "Slope = 0.0067", color = '#0080ff') +
    annotate("text", x = 33, y = 0.6, label = "Slope = 0.01", color = 'black') +
    theme_legend1()

# %% [markdown]
## HFD vs. LXN

# %% [code]
dataG <- data %>%
filter(Treatment == "HFD" | Treatment == 'HFD+LXN')
dataG$color[dataG$Treatment == 'HFD'] = '#0080ff'
dataG$color[dataG$Treatment == 'HFD+LXN'] = '#ffa500'

# %% [markdown]
### calculate R and slope

# %% [code]
dataG2 <- data %>% filter(Treatment == "HFD+LXN")
options(repr.plot.width = 4, repr.plot.height = 4, repr.plot.res = 200)
ggplotRegression(lm(avgEE2 ~ bw_cage, data = dataG2), "black") + labs(title = "LXN")

# %% [markdown]
### put together

# %% [code]
options(repr.plot.width = 6, repr.plot.height = 4, repr.plot.res = 200)
parB <- ggplot(dataG,aes(y = avgEE2, x = bw_cage, colour = color)) +
   geom_point(size=3) + geom_smooth(method = "lm", fill = NA) + 
    scale_y_continuous(limits = c(0.4, 0.7), breaks = seq(0.4, 0.7, 0.05)) +
    scale_x_continuous(limits = c(30, 55), breaks = seq(30, 55, 5)) +
    scale_color_identity() +
    ggtitle("**B.** <span style='color:#ffa500'>**LXN**</span> vs. <span style='color:#0080ff'>**HFD**</span>") + 
    ylab('energy expenditure (kcal/hr)') + 
    xlab('body mass upon cage entry (g)') + 
    annotate("text", x = 52, y = 0.53, label = "Slope = 0.0067", color = '#0080ff') +
    annotate("text", x = 34, y = 0.55, label = "Slope = 0.0063", color = '#ffa500') +
    theme_legend1()

# %% [markdown]
## HFD vs. HXN

# %% [code]
dataH <- data %>%
filter(Treatment == "HFD" | Treatment == 'HFD+HXN')
dataH$color[dataH$Treatment == 'HFD'] = '#0080ff'
dataH$color[dataH$Treatment == 'HFD+HXN'] = 'red'

# %% [markdown]
### calculate R and slope

# %% [code]
dataH2 <- data %>% filter(Treatment == "HFD+HXN")
options(repr.plot.width = 4, repr.plot.height = 4, repr.plot.res = 200)
ggplotRegression(lm(avgEE2 ~ bw_cage, data = dataH2), "black") + labs(title = "HXN")

# %% [markdown]
### put together

# %% [code]
options(repr.plot.width = 6, repr.plot.height = 4, repr.plot.res = 200)
parC <- ggplot(dataH,aes(y = avgEE2, x = bw_cage, colour = color)) +
   geom_point(size=3) + geom_smooth(method = "lm", fill = NA) + 
    scale_y_continuous(limits = c(0.4, 0.7), breaks = seq(0.4, 0.7, 0.05)) +
    scale_x_continuous(limits = c(30, 55), breaks = seq(30, 55, 5)) +
    scale_color_identity() +
    ggtitle("**C.** <span style='color:red'>**HXN**</span> vs. <span style='color:#0080ff'>**HFD**</span>") + 
    ylab('energy expenditure (kcal/hr)') + 
    xlab('body mass upon cage entry (g)') + 
    annotate("text", x = 52, y = 0.53, label = "Slope = 0.0067", color = '#0080ff') +
    annotate("text", x = 34, y = 0.45, label = "Slope = 0.0066", color = 'red') +
    theme_legend1()

# %% [markdown]
## HFD vs. TXN

# %% [code]
dataF <- data %>%
filter(Treatment == "HFD" | Treatment == 'HFD+TXN')
dataF$color[dataF$Treatment == 'HFD'] = '#0080ff'
dataF$color[dataF$Treatment == 'HFD+TXN'] = 'darkgreen'

# %% [code]
dataF2 <- data %>% filter(Treatment == "HFD+TXN")
options(repr.plot.width = 4, repr.plot.height = 4, repr.plot.res = 200)
ggplotRegression(lm(avgEE2 ~ bw_cage, data = dataF2), "black") + labs(title = "TXN")

# %% [code]
parD <- ggplot(dataF,aes(y = avgEE2, x = bw_cage, colour = color)) +
    geom_point(size=3) + geom_smooth(method = "lm", fill = NA) + 
    scale_y_continuous(limits = c(0.4, 0.7), breaks = seq(0.4, 0.7, 0.05)) +
    scale_x_continuous(limits = c(30, 55), breaks = seq(30, 55, 5)) +
    scale_color_identity() +
    ggtitle("**D.** <span style='color:darkgreen'>**TXN**</span> vs. <span style='color:#0080ff'>**HFD**</span>") + 
    ylab('energy expenditure (kcal/hr)') + 
    xlab('body mass upon cage entry (g)') + 
    annotate("text", x = 52, y = 0.53, label = "Slope = 0.0067", color = '#0080ff') +
    annotate("text", x = 33, y = 0.52, label = "Slope = 0.0087", color = 'darkgreen') +
    theme_legend1()

# %% [code]
options(repr.plot.width = 12, repr.plot.height = 8, repr.plot.res = 200)
figS2 <- (parA | parB | parC | parD) + plot_layout(nrow = 2, ncol=2)

# %% [code]
#graph2pdf(figS2, "fig2Sup1.pdf", width = 12, height = 8)

# %% [markdown]
References:
- https://www.infoworld.com/article/3527449/add-color-to-your-ggplot2-text-in-r.html
- http://www.sthda.com/english/wiki/ggplot2-texts-add-text-annotations-to-a-graph-in-r-software
```
