## Supplementary material for "TXN, a Xanthohumol Derivative, Attenuates High-Fat Diet Induced Hepatic Steatosis by Antagonizing PPARγ": Figure 3 source data: fig3Sup2.docx

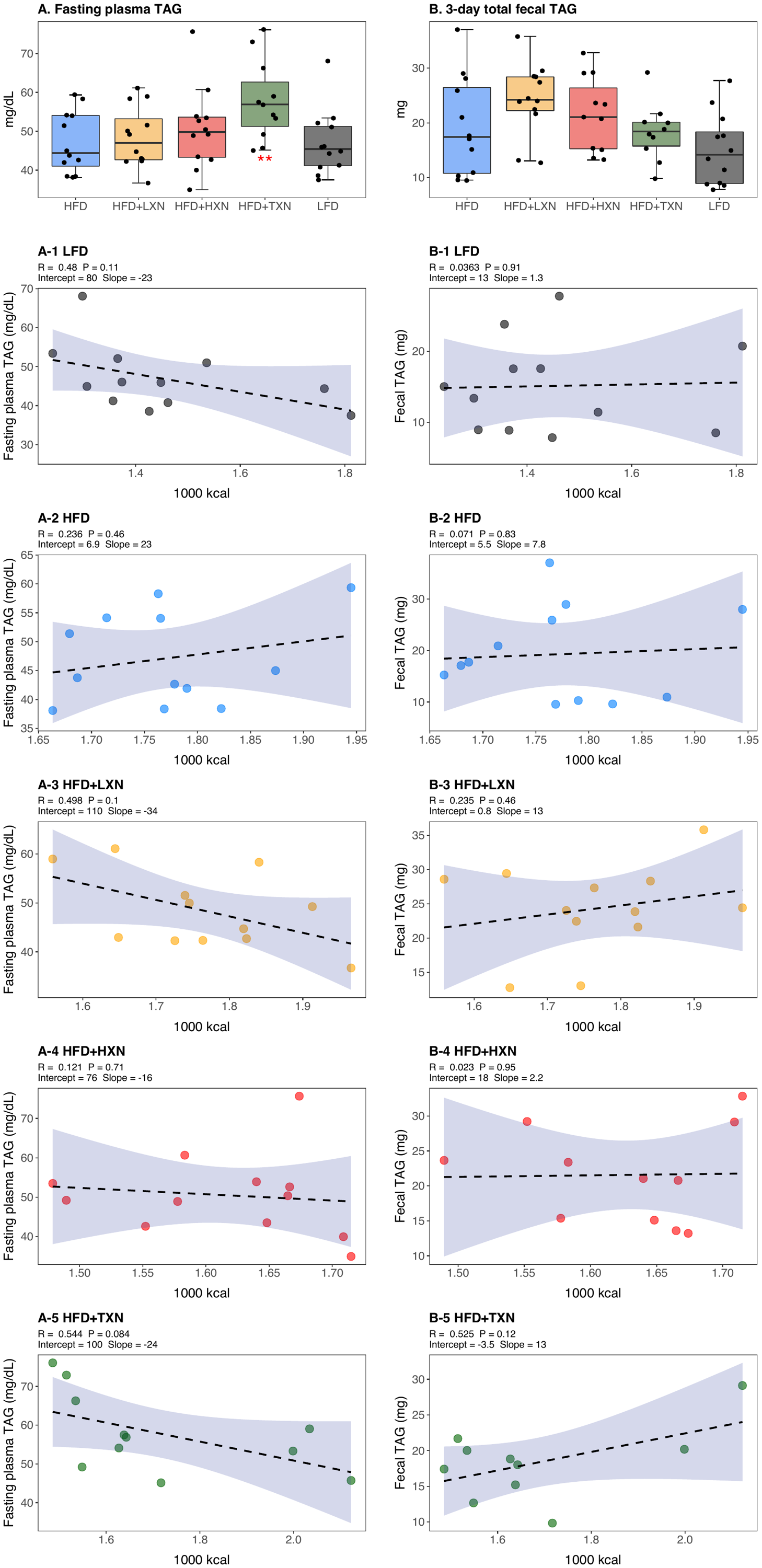

Figure supplement 2. The effect of diet and intervention on fasting plasma and fecal TAG levels. Mice were fed either a LFD (black, n = 12), a HFD (blue, n = 12), HFD+LXN (yellow, n = 12), HFD+HXN (red, n = 12), or HFD+TXN (green, n = 11) for 16 weeks. (A) Fasting plasma TAG levels are expressed as quartiles. (A-1) Relationship between fasting plasma TAG and total caloric intake over 16 weeks of feeding for LFD group; (A-2) for HFD group; (A-3) for HFD+LXN group; (A-4) for HFD+HXN group; (A-5) for HFD+TXN group (B) 3-day total fecal triglyceride (TAG) are expressed as quartiles. (B-1) Relationship between 3-day fecal TAG and total caloric intake over 16 weeks of feeding for LFD group; (B-2) for HFD group; (B-3) for HFD+LXN group; (B-4) for HFD+HXN group; (B-5) for HFD+TXN group. Pre-planned general linear model with contrasts were used to calculate *p*-values in A, B and C. **p* < 0.05, ***p* < 0.01, ****p* < 0.001. Linear regression analyses of total calories versus fasting plasma TAG (A1-5) or total fecal TAG (B1-5) in mice were done using lm function of stats package version 3.6.2 in R. Blue shading represents 95% CI of the regression line. Absolute value of R, p-value, intercept, and slope for the regression are reported above each corresponding panel.
