## Supplementary material for "TXN, a Xanthohumol Derivative, Attenuates High-Fat Diet Induced Hepatic Steatosis by Antagonizing PPARγ": Figure 3 source data: fig3Sup2.ipynb

```
# %% [code]
suppressPackageStartupMessages(library(dplyr))
suppressPackageStartupMessages(library(ggplot2))
suppressPackageStartupMessages(library(lsmeans))
suppressPackageStartupMessages(library(patchwork))
suppressPackageStartupMessages(library(export))
suppressPackageStartupMessages(library(ggtext))
R.Version()

# %% [markdown]
# Import data

# %% [markdown]
## - Phenotypes

# %% [code]
data <- read.csv("supplement2Table.csv", header = T); 
data$vars <- factor(data$Treatment, levels = c("LFD", "HFD", "HFD+LXN", "HFD+HXN", "HFD+TXN"))
data$vars2 <- factor(data$Treatment, levels = c("HFD", "HFD+LXN", "HFD+HXN", "HFD+TXN", "LFD"))
data <- data %>% mutate(kkcal = Cal_cum/1000,
                        fecal_tot_tag = TG_feces2*fecal_tot_output); head(data)

# %% [code]
txn <- data %>% filter(vars2 == "HFD+TXN"); hfd <- data %>% filter(vars2 == "HFD");
lxn <- data %>% filter(vars2 == "HFD+LXN"); hxn <- data %>% filter(vars2 == "HFD+HXN");
lfd <- data %>% filter(vars2 == "LFD")

# %% [markdown]
# Set theme, import functions

# %% [code]
theme_set(theme_bw())

# %% [code]
theme_legend2 = function(){
	theme(axis.title.y = element_text(size = 15, 
	margin = margin(t=0, r=5, b=0, l=5)),
	axis.text.y = element_text(size = 13),
	axis.text.x = element_text(size = 13),
	axis.title.x = element_text(size = 15,
                               margin = margin(t=10, r=5, b=0, l=0)),
    plot.title = element_text(size = 15, face = "bold"),
	legend.title = element_text(colour = "white"),
	legend.text = element_text(size = 16),
	legend.position = "none",
	plot.margin = unit(c(0.5,0.5,0,0.5), "cm"),
	panel.grid = element_blank())
}

# %% [code]
source("ggplotRegression.R")

# %% [code]
mycolor <- c("#0080ff", "#ffa500", "red", "darkgreen", "black")
mycolor_alpha <- c("#8AB5F9", "#F8CB89", "#F08581", "#87A57E", "#797979")

# %% [code]
Contrasts <- list(
    'LFD vs HFD' = c(-1, 0, 0, 0, 1),
    'LXN vs HFD' = c(-1, 1, 0, 0, 0),
    'HXN vs HFD' = c(-1, 0, 1, 0, 0),
    'TXN vs HFD' = c(-1, 0, 0, 1, 0))

# %% [markdown]
# Figures

# %% [markdown]
## - A. fat mass boxplot

# %% [code]
theme_legend1 = function(){
	theme(axis.title.y = element_text(size = 15, 
	margin = margin(t=0, r=5, b=0, l=5)),
	axis.text.y = element_text(size = 13),
	axis.text.x = element_text(size = 13),
	axis.title.x = element_text(size = 15,
                               margin = margin(t=10, r=5, b=0, l=0)),
    #plot.title = element_text(size = 15, face = "bold"),
          plot.title = element_markdown(),
	legend.title = element_text(colour = "white"),
	legend.text = element_text(size = 16),
	legend.position = c(0.8, 0.2),
	plot.margin = unit(c(0.5,0.5,0,0.5), "cm"),
	panel.grid = element_blank())
}

# %% [code]
model <- lm(plasma_TAG ~ vars2, data = data)
leastsquare <- lsmeans(model, "vars2")
contrast(leastsquare, Contrasts, adjust = "none")

# %% [code]
A <- ggplot(data = data, aes(x = vars2, y = plasma_TAG, fill = vars2)) +
	stat_boxplot(geom = 'errorbar', linetype = 1, width = 0.2) +
	geom_boxplot(outlier.shape = NA) +
	scale_fill_manual(values = mycolor_alpha) +
	geom_jitter(shape = 20, size = 3, width = 0.2, height = 0.1) +
    labs(x="",y="mg/dL",title="A. Fasting plasma TAG") +
	theme_legend2()

label.df <- data.frame(vars2 = c("HFD+TXN"),
                       plasma_TAG = c(42))
A <- A + geom_text(data = label.df, label = c("**"), color='red', size = 8)

# %% [markdown]
## - B. average energy expenditure

# %% [code]
model <- lm(fecal_tot_tag ~ vars2, data = data)
leastsquare <- lsmeans(model, "vars2")
contrast(leastsquare, Contrasts, adjust = "none")

# %% [code]
C <- ggplot(data = data, aes(x = vars2, y = fecal_tot_tag, fill = vars2)) +
	stat_boxplot(geom = 'errorbar', linetype = 1, width = 0.2) +
	geom_boxplot(outlier.shape = NA) +
	scale_fill_manual(values = mycolor_alpha) +
	geom_jitter(shape = 20, size = 3, width = 0.2, height = 0.1) +
    labs(x="",y="mg",title="B. 3-day total fecal TAG") +
	theme_legend2()

# %% [code]
options(repr.plot.width = 20, repr.plot.height = 4, repr.plot.res = 150)
row1 <- (A | C) + plot_layout(nrow = 1, ncol=2)

# %% [code]
lfd_plasma_TAG_cal <- ggplotRegression(lm(plasma_TAG ~ kkcal, data = lfd), "black") + 
                        labs(title = "A-1 LFD", x = "1000 kcal", y = "Fasting plasma TAG (mg/dL)")+ theme_legend2()
hfd_plasma_TAG_cal <- ggplotRegression(lm(plasma_TAG ~ kkcal, data = hfd), "#0080ff")+ 
                        labs(title = "A-2 HFD", x = "1000 kcal", y = "Fasting plasma TAG (mg/dL)")+ theme_legend2()
lxn_plasma_TAG_cal <- ggplotRegression(lm(plasma_TAG ~ kkcal, data = lxn), "#ffa500")+ 
                        labs(title = "A-3 HFD+LXN", x = "1000 kcal", y = "Fasting plasma TAG (mg/dL)")+ theme_legend2()
hxn_plasma_TAG_cal <- ggplotRegression(lm(plasma_TAG ~ kkcal, data = hxn), "red")+ ggtitle("HXN")+ 
                        labs(title = "A-4 HFD+HXN", x = "1000 kcal", y = "Fasting plasma TAG (mg/dL)")+ theme_legend2()
txn_plasma_TAG_cal <- ggplotRegression(lm(plasma_TAG ~ kkcal, data = txn), "darkgreen")+ 
                        labs(title = "A-5 HFD+TXN", x = "1000 kcal", y = "Fasting plasma TAG (mg/dL)")+ theme_legend2()

# %% [code]
lfd_fecalTAG_cal <- ggplotRegression(lm(fecal_tot_tag ~ kkcal, data = lfd), "black") + 
                        labs(title = "B-1 LFD", x = "1000 kcal", y = "Fecal TAG (mg)")+ theme_legend2()
hfd_fecalTAG_cal <- ggplotRegression(lm(fecal_tot_tag ~ kkcal, data = hfd), "#0080ff")+ 
                        labs(title = "B-2 HFD", x = "1000 kcal", y = "Fecal TAG (mg)")+ theme_legend2()
lxn_fecalTAG_cal <- ggplotRegression(lm(fecal_tot_tag ~ kkcal, data = lxn), "#ffa500")+ 
                        labs(title = "B-3 HFD+LXN", x = "1000 kcal", y = "Fecal TAG (mg)")+ theme_legend2()
hxn_fecalTAG_cal <- ggplotRegression(lm(fecal_tot_tag ~ kkcal, data = hxn), "red")+ ggtitle("HXN")+ 
                        labs(title = "B-4 HFD+HXN", x = "1000 kcal", y = "Fecal TAG (mg)")+ theme_legend2()
txn_fecalTAG_cal <- ggplotRegression(lm(fecal_tot_tag ~ kkcal, data = txn), "darkgreen")+ 
                        labs(title = "B-5 HFD+TXN", x = "1000 kcal", y = "Fecal TAG (mg)")+ theme_legend2()

# %% [code]
options(repr.plot.width = 20, repr.plot.height = 4, repr.plot.res = 150)
row2 <- (lfd_plasma_TAG_cal | lfd_fecalTAG_cal) + plot_layout(nrow = 1, ncol = 2)

# %% [code]
options(repr.plot.width = 20, repr.plot.height = 4, repr.plot.res = 150)
row3 <- (hfd_plasma_TAG_cal | hfd_fecalTAG_cal) + plot_layout(nrow = 1, ncol = 2)

# %% [code]
options(repr.plot.width = 20, repr.plot.height = 4, repr.plot.res = 150)
row4 <- (lxn_plasma_TAG_cal | lxn_fecalTAG_cal) + plot_layout(nrow = 1, ncol = 2)

# %% [code]
options(repr.plot.width = 20, repr.plot.height = 4, repr.plot.res = 150)
row5 <- (hxn_plasma_TAG_cal | hxn_fecalTAG_cal) + plot_layout(nrow = 1, ncol = 2)

# %% [code]
options(repr.plot.width = 20, repr.plot.height = 4, repr.plot.res = 150)
row6 <- (txn_plasma_TAG_cal | txn_fecalTAG_cal) + plot_layout(nrow = 1, ncol = 2)

# %% [markdown]
## Put together

# %% [code]
options(repr.plot.width = 12.5, repr.plot.height = 25, repr.plot.res = 200)
figure3.2 <- wrap_plots(row1, row2, row3, row4, row5, row6, nrow=6)

# %% [code]
#graph2pdf(figure3.2, "fig3Sup2.pdf", width = 12.5, height = 25)

# %% [markdown]
## Correlations

# %% [code]
.ci <-
function(x,conf.level=0.95) {
  tri <- sort(na.omit(x))
  if (any(!is.finite(tri))) {tri <- tri[-which(!is.finite(tri))]}
  repet <- length(tri)
  int <- (1-conf.level)/2
  if(repet*int<1) {
    int.inf <- ceiling(repet*int)
  } else {
    int.inf <- floor(repet*int)
  }
  int.sup <- ceiling(repet*(1-int))
  result <- c("Inf"=tri[int.inf],"Sup"=tri[int.sup])
  return(result)
}

spearman.ci <-
function(var1,var2,nrep=1000,conf.level=0.95){
  if (length(var1)!=length(var2)) {stop(paste("'",deparse(substitute(var1)),"' and '",
    deparse(substitute(var2)),"' lengths differ",sep=""))}
  data.name <- paste(deparse(substitute(var1))," and ",deparse(substitute(var2)),"\n",nrep," replicates",sep="")
  nul <- as.numeric(row.names(table(c(which(is.na(var1)),which(is.na(var2))))))
  var1.2 <- if(length(nul)>0) {var1[-nul]} else {var1}
  var2.2 <- if(length(nul)>0) {var2[-nul]} else {var2}
  cor.fun <- function(data,ind) {
    as.numeric(suppressWarnings(cor.test(data[ind,1],data[ind,2],method="spearman")$estimate))
  }
  simul <- boot::boot(data.frame(var1.2,var2.2),cor.fun,R=nrep)
  interval <- .ci(simul$t,conf.level=conf.level)
  attr(interval,"conf.level") <- conf.level
  coeff <- as.numeric(suppressWarnings(cor.test(var1,var2,method="spearman")$estimate))
  names(coeff) <- "rho"
  result <- list(method="Spearman's rank correlation",conf.level=conf.level,rep=nrep,data.name=data.name,
    estimate=coeff,conf.int=interval)
  class(result) <- "htest"
  return(result)
}

pearson.ci <-
function(var1,var2,nrep=1000,conf.level=0.95){
  if (length(var1)!=length(var2)) {stop(paste("'",deparse(substitute(var1)),"' and '",
    deparse(substitute(var2)),"' lengths differ",sep=""))}
  data.name <- paste(deparse(substitute(var1))," and ",deparse(substitute(var2)),"\n",nrep," replicates",sep="")
  nul <- as.numeric(row.names(table(c(which(is.na(var1)),which(is.na(var2))))))
  var1.2 <- if(length(nul)>0) {var1[-nul]} else {var1}
  var2.2 <- if(length(nul)>0) {var2[-nul]} else {var2}
  cor.fun <- function(data,ind) {
    as.numeric(suppressWarnings(cor.test(data[ind,1],data[ind,2],method="pearson")$estimate))
  }
  simul <- boot::boot(data.frame(var1.2,var2.2),cor.fun,R=nrep)
  interval <- .ci(simul$t,conf.level=conf.level)
  attr(interval,"conf.level") <- conf.level
  coeff <- as.numeric(suppressWarnings(cor.test(var1,var2,method="pearson")$estimate))
  names(coeff) <- "rho"
  result <- list(method="pearson correlation",conf.level=conf.level,rep=nrep,data.name=data.name,
    estimate=coeff,conf.int=interval)
  class(result) <- "htest"
  return(result)
}

# %% [markdown]
## A

# %% [code]
data %>%
    summarise(r = pearson.ci(kkcal, plasma_TAG)$estimate,
             pvalue = cor.test(kkcal, plasma_TAG, method = "pearson")$p.value,
             lower = pearson.ci(kkcal, plasma_TAG)$conf.int[1],
             upper = pearson.ci(kkcal, plasma_TAG)$conf.int[2])

# %% [code]
data %>%
    summarise(r = spearman.ci(kkcal, plasma_TAG)$estimate,
             pvalue = cor.test(kkcal, plasma_TAG, method = "spearman")$p.value,
             lower = spearman.ci(kkcal, plasma_TAG)$conf.int[1],
             upper = spearman.ci(kkcal, plasma_TAG)$conf.int[2])

# %% [markdown]
## B

# %% [code]
data %>%
    summarise(r = pearson.ci(kkcal, fecal_tot_tag)$estimate,
             pvalue = cor.test(kkcal, fecal_tot_tag, method = "pearson")$p.value,
             lower = pearson.ci(kkcal, fecal_tot_tag)$conf.int[1],
             upper = pearson.ci(kkcal, fecal_tot_tag)$conf.int[2])

# %% [code]
data %>%
    summarise(r = spearman.ci(kkcal, fecal_tot_tag)$estimate,
             pvalue = cor.test(kkcal, fecal_tot_tag, method = "spearman")$p.value,
             lower = spearman.ci(kkcal, fecal_tot_tag)$conf.int[1],
             upper = spearman.ci(kkcal, fecal_tot_tag)$conf.int[2])

# %% [markdown]
## A-1. LFD

# %% [code]
data_lfd <- data %>% filter(Treatment == "LFD")

# %% [code]
data_lfd %>%
    summarise(r = pearson.ci(kkcal, plasma_TAG)$estimate,
             pvalue = cor.test(kkcal, plasma_TAG, method = "pearson")$p.value,
             lower = pearson.ci(kkcal, plasma_TAG)$conf.int[1],
             upper = pearson.ci(kkcal, plasma_TAG)$conf.int[2])

# %% [code]
data_lfd %>%
    summarise(r = spearman.ci(kkcal, plasma_TAG)$estimate,
             pvalue = cor.test(kkcal, plasma_TAG, method = "spearman")$p.value,
             lower = spearman.ci(kkcal, plasma_TAG)$conf.int[1],
             upper = spearman.ci(kkcal, plasma_TAG)$conf.int[2])

# %% [markdown]
## A-2. HFD

# %% [code]
data_hfd <- data %>% filter(Treatment == "HFD")

# %% [code]
data_hfd %>%
    summarise(r = pearson.ci(kkcal, plasma_TAG)$estimate,
             pvalue = cor.test(kkcal, plasma_TAG, method = "pearson")$p.value,
             lower = pearson.ci(kkcal, plasma_TAG)$conf.int[1],
             upper = pearson.ci(kkcal, plasma_TAG)$conf.int[2])

# %% [code]
data_hfd %>%
    summarise(r = spearman.ci(kkcal, plasma_TAG)$estimate,
             pvalue = cor.test(kkcal, plasma_TAG, method = "spearman")$p.value,
             lower = spearman.ci(kkcal, plasma_TAG)$conf.int[1],
             upper = spearman.ci(kkcal, plasma_TAG)$conf.int[2])

# %% [markdown]
## A-3.LXN

# %% [code]
dataLXN <- data %>% filter(Treatment == "HFD+LXN")

# %% [code]
dataLXN %>%
    summarise(r = pearson.ci(kkcal, plasma_TAG)$estimate,
             pvalue = cor.test(kkcal, plasma_TAG, method = "pearson")$p.value,
             lower = pearson.ci(kkcal, plasma_TAG)$conf.int[1],
             upper = pearson.ci(kkcal, plasma_TAG)$conf.int[2])

# %% [code]
dataLXN %>%
    summarise(r = spearman.ci(kkcal, plasma_TAG)$estimate,
             pvalue = cor.test(kkcal, plasma_TAG, method = "spearman")$p.value,
             lower = spearman.ci(kkcal, plasma_TAG)$conf.int[1],
             upper = spearman.ci(kkcal, plasma_TAG)$conf.int[2])

# %% [markdown]
## A-4.HXN

# %% [code]
dataHXN <- data %>% filter(Treatment == "HFD+HXN")

# %% [code]
dataHXN %>%
    summarise(r = pearson.ci(kkcal, plasma_TAG)$estimate,
             pvalue = cor.test(kkcal, plasma_TAG, method = "pearson")$p.value,
             lower = pearson.ci(kkcal, plasma_TAG)$conf.int[1],
             upper = pearson.ci(kkcal, plasma_TAG)$conf.int[2])

# %% [code]
dataHXN %>%
    summarise(r = spearman.ci(kkcal, plasma_TAG)$estimate,
             pvalue = cor.test(kkcal, plasma_TAG, method = "spearman")$p.value,
             lower = spearman.ci(kkcal, plasma_TAG)$conf.int[1],
             upper = spearman.ci(kkcal, plasma_TAG)$conf.int[2])

# %% [markdown]
## A-5. TXN

# %% [code]
dataTXN <- data %>% filter(Treatment == "HFD+TXN")

# %% [code]
dataTXN %>%
    summarise(r = pearson.ci(kkcal, plasma_TAG)$estimate,
             pvalue = cor.test(kkcal, plasma_TAG, method = "pearson")$p.value,
             lower = pearson.ci(kkcal, plasma_TAG)$conf.int[1],
             upper = pearson.ci(kkcal, plasma_TAG)$conf.int[2])

# %% [code]
dataTXN %>%
    summarise(r = spearman.ci(kkcal, plasma_TAG)$estimate,
             pvalue = cor.test(kkcal, plasma_TAG, method = "spearman")$p.value,
             lower = spearman.ci(kkcal, plasma_TAG)$conf.int[1],
             upper = spearman.ci(kkcal, plasma_TAG)$conf.int[2])
```
