## Supplementary material for "TXN, a Xanthohumol Derivative, Attenuates High-Fat Diet Induced Hepatic Steatosis by Antagonizing PPARγ": Figure 3 source data: figure3.ipynb

```
# %% [code]
## Load packages

# %% [code]
suppressPackageStartupMessages(library(dplyr))
suppressPackageStartupMessages(library(ggplot2))
suppressPackageStartupMessages(library(lsmeans))
suppressPackageStartupMessages(library(patchwork))
suppressPackageStartupMessages(library(export))
R.Version()

# %% [markdown]
# Import data

# %% [markdown]
## - Promethion

# %% [code]
profile <- read.csv(file = "fig2_stat_corrected.csv", as.is = TRUE)
profile$Diet <- as.factor(profile$Diet)
profile$Diet <- factor(profile$Diet, levels = c("HFD", "HFD+LXN", "HFD+HXN", "HFD+TXN", "LFD"))
profile$MouseID <- ordered(profile$MouseID)
profile$MouseID <- as.factor(profile$MouseID); head(profile)

# %% [markdown]
## - Phenotypes

# %% [code]
data <- read.csv("fig2_table.csv", header = T); 
data$vars <- factor(data$Treatment, levels = c("LFD", "HFD", "HFD+LXN", "HFD+HXN", "HFD+TXN"))
data$vars2 <- factor(data$Treatment, levels = c("HFD", "HFD+LXN", "HFD+HXN", "HFD+TXN", "LFD"))
data <- data %>% mutate(kkcal = Cal_cum/1000); head(data)
txn <- data %>% filter(vars2 == "HFD+TXN"); hfd <- data %>% filter(vars2 == "HFD");
lxn <- data %>% filter(vars2 == "HFD+LXN"); hxn <- data %>% filter(vars2 == "HFD+HXN");
lfd <- data %>% filter(vars2 == "LFD")

# %% [markdown]
# Set theme, import functions

# %% [code]
theme_set(theme_bw())

# %% [code]
theme_legend2 = function(){
	theme(axis.title.y = element_text(size = 15, 
	margin = margin(t=0, r=5, b=0, l=5)),
	axis.text.y = element_text(size = 13),
	axis.text.x = element_text(size = 13),
	axis.title.x = element_text(size = 15,
                               margin = margin(t=10, r=5, b=0, l=0)),
    plot.title = element_text(size = 15, face = "bold"),
	legend.title = element_text(colour = "white"),
	legend.text = element_text(size = 16),
	legend.position = "none",
	plot.margin = unit(c(0.5,0.5,0,0.5), "cm"),
	panel.grid = element_blank())
}

# %% [code]
source("ggplotRegression.R")

# %% [code]
mycolor <- c("#0080ff", "#ffa500", "red", "darkgreen", "black")
mycolor_alpha <- c("#8AB5F9", "#F8CB89", "#F08581", "#87A57E", "#797979")

# %% [code]
Contrasts <- list(
    'LFD vs HFD' = c(-1, 0, 0, 0, 1),
    'LXN vs HFD' = c(-1, 1, 0, 0, 0),
    'HXN vs HFD' = c(-1, 0, 1, 0, 0),
    'TXN vs HFD' = c(-1, 0, 0, 1, 0))

# %% [markdown]
# Figures

# %% [markdown]
## - A. fat mass boxplot

# %% [code]
model <- lm(Fat_mass ~ vars2, data = data)
leastsquare <- lsmeans(model, "vars2")
contrast(leastsquare, Contrasts, adjust = "none")

# %% [code]
A <- ggplot(data = data, aes(x = vars2, y = Fat_mass, fill = vars2)) +
	stat_boxplot(geom = 'errorbar', linetype = 1, width = 0.2) +
	geom_boxplot(outlier.shape = NA) +
	scale_fill_manual(values = mycolor_alpha) +
	geom_jitter(shape = 20, size = 3, width = 0.2, height = 0.1) +
    labs(x="",y="g",title="A. Fat mass") +
	theme_legend2()

label.df <- data.frame(vars2 = c("HFD+HXN", "HFD+TXN", "LFD"),
                       Fat_mass = c(21.5, 20, 15))
A <- A + geom_text(data = label.df, label = c("*", "***", "***"), color='red', size = 8)

# %% [markdown]
## - B. Hepatic lipidosis area percent

# %% [code]
model <- lm(lipidosis_area_pct ~ vars2, data = data)
leastsquare <- lsmeans(model, "vars2")
contrast(leastsquare, Contrasts, adjust = "none")

# %% [code]
B <- ggplot(data = data, aes(x = vars2, y = lipidosis_area_pct, fill = vars2)) +
	stat_boxplot(geom = 'errorbar', linetype = 1, width = 0.2) +
	geom_boxplot(outlier.shape = NA) +
	scale_fill_manual(values = mycolor_alpha) +
	geom_jitter(shape = 20, size = 3, width = 0.2, height = 0.1) +
    labs(x="",y="%",title="B. Hepatic lipidosis area percent") +
	theme_legend2()
label.df <- data.frame(vars2 = c("HFD+HXN", "HFD+TXN"),
                       lipidosis_area_pct = c(22, 20))
B <- B + geom_text(data = label.df, label = c("*", "**"), color='red', size = 8)

# %% [markdown]
## - C. average energy expenditure

# %% [code]
model <- lm(avgEE2 ~ vars2, data = data)
leastsquare <- lsmeans(model, "vars2")
contrast(leastsquare, Contrasts, adjust = "none")

# %% [code]
C <- ggplot(data = data, aes(x = vars2, y = avgEE2, fill = vars2)) +
	stat_boxplot(geom = 'errorbar', linetype = 1, width = 0.2) +
	geom_boxplot(outlier.shape = NA) +
	scale_fill_manual(values = mycolor_alpha) +
	geom_jitter(shape = 20, size = 3, width = 0.2, height = 0.1) +
    labs(x="",y="kcal/hr",title="C. Average energy expenditure") +
	theme_legend2()

label.df <- data.frame(vars2 = c("HFD+HXN", "HFD+TXN", "LFD"),
                       avgEE2 = c(0.6, 0.55, 0.58))
C <- C + geom_text(data = label.df, label = c("*", "**", "*"), color='red', size = 8)

# %% [markdown]
# Regression plots

# %% [code]
lfd_Fat_mass_cal <- ggplotRegression(lm(Fat_mass ~ kkcal, data = lfd), "black") + 
                        labs(title = "A-1 LFD", x = "1000 kcal", y = "fat mass (g)")+ theme_legend2()
hfd_Fat_mass_cal <- ggplotRegression(lm(Fat_mass ~ kkcal, data = hfd), "#0080ff")+ 
                        labs(title = "A-2 HFD", x = "1000 kcal", y = "fat mass (g)")+ theme_legend2()
lxn_Fat_mass_cal <- ggplotRegression(lm(Fat_mass ~ kkcal, data = lxn), "#ffa500")+ 
                        labs(title = "A-3 HFD+LXN", x = "1000 kcal", y = "fat mass (g)")+ theme_legend2()
hxn_Fat_mass_cal <- ggplotRegression(lm(Fat_mass ~ kkcal, data = hxn), "red")+ ggtitle("HXN")+ 
                        labs(title = "A-4 HFD+HXN", x = "1000 kcal", y = "fat mass (g)")+ theme_legend2()
txn_Fat_mass_cal <- ggplotRegression(lm(Fat_mass ~ kkcal, data = txn), "darkgreen")+ 
                        labs(title = "A-5 HFD+TXN", x = "1000 kcal", y = "lean mass (g)")+ theme_legend2()

# %% [code]
lfd_lipidosis_area_pct_cal <- ggplotRegression(lm(lipidosis_area_pct ~ kkcal, data = lfd), "black") + 
                        labs(title = "B-1 LFD", x = "1000 kcal", y = "hepatic lipidosis area percent")+ theme_legend2()
hfd_lipidosis_area_pct_cal <- ggplotRegression(lm(lipidosis_area_pct ~ kkcal, data = hfd), "#0080ff")+ 
                        labs(title = "B-2 HFD", x = "1000 kcal", y = "hepatic lipidosis area percent")+ theme_legend2()
lxn_lipidosis_area_pct_cal <- ggplotRegression(lm(lipidosis_area_pct ~ kkcal, data = lxn), "#ffa500")+ 
                        labs(title = "B-3 HFD+LXN", x = "1000 kcal", y = "hepatic lipidosis area percent")+ theme_legend2()
hxn_lipidosis_area_pct_cal <- ggplotRegression(lm(lipidosis_area_pct ~ kkcal, data = hxn), "red")+ ggtitle("HXN")+ 
                        labs(title = "B-4 HFD+HXN", x = "1000 kcal", y = "hepatic lipidosis area percent")+ theme_legend2()
txn_lipidosis_area_pct_cal <- ggplotRegression(lm(lipidosis_area_pct ~ kkcal, data = txn), "darkgreen")+ 
                        labs(title = "B-5 HFD+TXN", x = "1000 kcal", y = "hepatic lipidosis area percent")+ theme_legend2()

# %% [code]
lfd_EE_cal <- ggplotRegression(lm(avgEE ~ kkcal, data = lfd), "black") + 
                        labs(title = "C-1 LFD", x = "1000 kcal", y = "energy expenditure")+ theme_legend2()
hfd_EE_cal <- ggplotRegression(lm(avgEE ~ kkcal, data = hfd), "#0080ff")+ 
                        labs(title = "C-2 HFD", x = "1000 kcal", y = "energy expenditure")+ theme_legend2()
lxn_EE_cal <- ggplotRegression(lm(avgEE ~ kkcal, data = lxn), "#ffa500")+ 
                        labs(title = "C-3 HFD+LXN", x = "1000 kcal", y = "energy expenditure")+ theme_legend2()
hxn_EE_cal <- ggplotRegression(lm(avgEE2 ~ kkcal, data = hxn), "red")+ ggtitle("HXN")+ 
                        labs(title = "C-4 HFD+HXN", x = "1000 kcal", y = "energy expenditure")+ theme_legend2()
txn_EE_cal <- ggplotRegression(lm(avgEE ~ kkcal, data = txn), "darkgreen")+ 
                        labs(title = "C-5 HFD+TXN", x = "1000 kcal", y = "energy expenditure")+ theme_legend2()

# %% [markdown]
## Put together

# %% [code]
options(repr.plot.width = 20, repr.plot.height = 4, repr.plot.res = 150)
row1 <- (A | B | C) + plot_layout(nrow = 1, ncol=3)

# %% [code]
options(repr.plot.width = 20, repr.plot.height = 4, repr.plot.res = 150)
row2 <- (lfd_Fat_mass_cal | lfd_lipidosis_area_pct_cal | lfd_EE_cal) + plot_layout(nrow = 1, ncol = 3)

# %% [code]
options(repr.plot.width = 20, repr.plot.height = 4, repr.plot.res = 150)
row3 <- (hfd_Fat_mass_cal | hfd_lipidosis_area_pct_cal | hfd_EE_cal) + plot_layout(nrow = 1, ncol = 3)

# %% [code]
options(repr.plot.width = 20, repr.plot.height = 4, repr.plot.res = 150)
row4 <- (lxn_Fat_mass_cal | lxn_lipidosis_area_pct_cal | lxn_EE_cal) + plot_layout(nrow = 1, ncol = 3)

# %% [code]
options(repr.plot.width = 20, repr.plot.height = 4, repr.plot.res = 150)
row5 <- (hxn_Fat_mass_cal | hxn_lipidosis_area_pct_cal | hxn_EE_cal) + plot_layout(nrow = 1, ncol = 3)

# %% [code]
options(repr.plot.width = 20, repr.plot.height = 4, repr.plot.res = 150)
row6 <- (txn_Fat_mass_cal | txn_lipidosis_area_pct_cal | txn_EE_cal) + plot_layout(nrow = 1, ncol = 3)

# %% [code]
options(repr.plot.width = 20, repr.plot.height = 25, repr.plot.res = 200)
figure2 <- wrap_plots(row1, row2, row3, row4, row5, row6, nrow=6)

# %% [code]
#graph2pdf(figure2, "figure2.pdf", width = 20, height = 25)
```
