## Supplementary material for "TXN, a Xanthohumol Derivative, Attenuates High-Fat Diet Induced Hepatic Steatosis by Antagonizing PPARγ": Figure 3 source data: ggplotRegression.R

```
ggplotRegression <- function (fit, cols) {
    
    ggplot(fit$model, aes_string(x = names(fit$model)[2], y = names(fit$model)[1])) + 
          geom_point(size = 4, color = cols, alpha = 0.6) +
          stat_smooth(method = "lm", col = "black", geom = "smooth", fill = "#303F9F", alpha = 0.2, linetype = "dashed") +
          labs(subtitle = paste("R = ",signif(sqrt(summary(fit)$r.squared), 3),
                                " P =",signif(summary(fit)$coef[2,4], 2),
                               "\nIntercept =",signif(fit$coef[[1]],2 ),
                                 " Slope =",signif(fit$coef[[2]], 2))) #+
          #theme(axis.title = element_text(size= 20, face = 'bold'),
          #axis.text = element_text(size = 18))
}
```
