## Supplementary figures and images for "TXN, a Xanthohumol Derivative, Attenuates High-Fat Diet Induced Hepatic Steatosis by Antagonizing PPARγ"

### 5 uM TXN+Rosi2.jpg

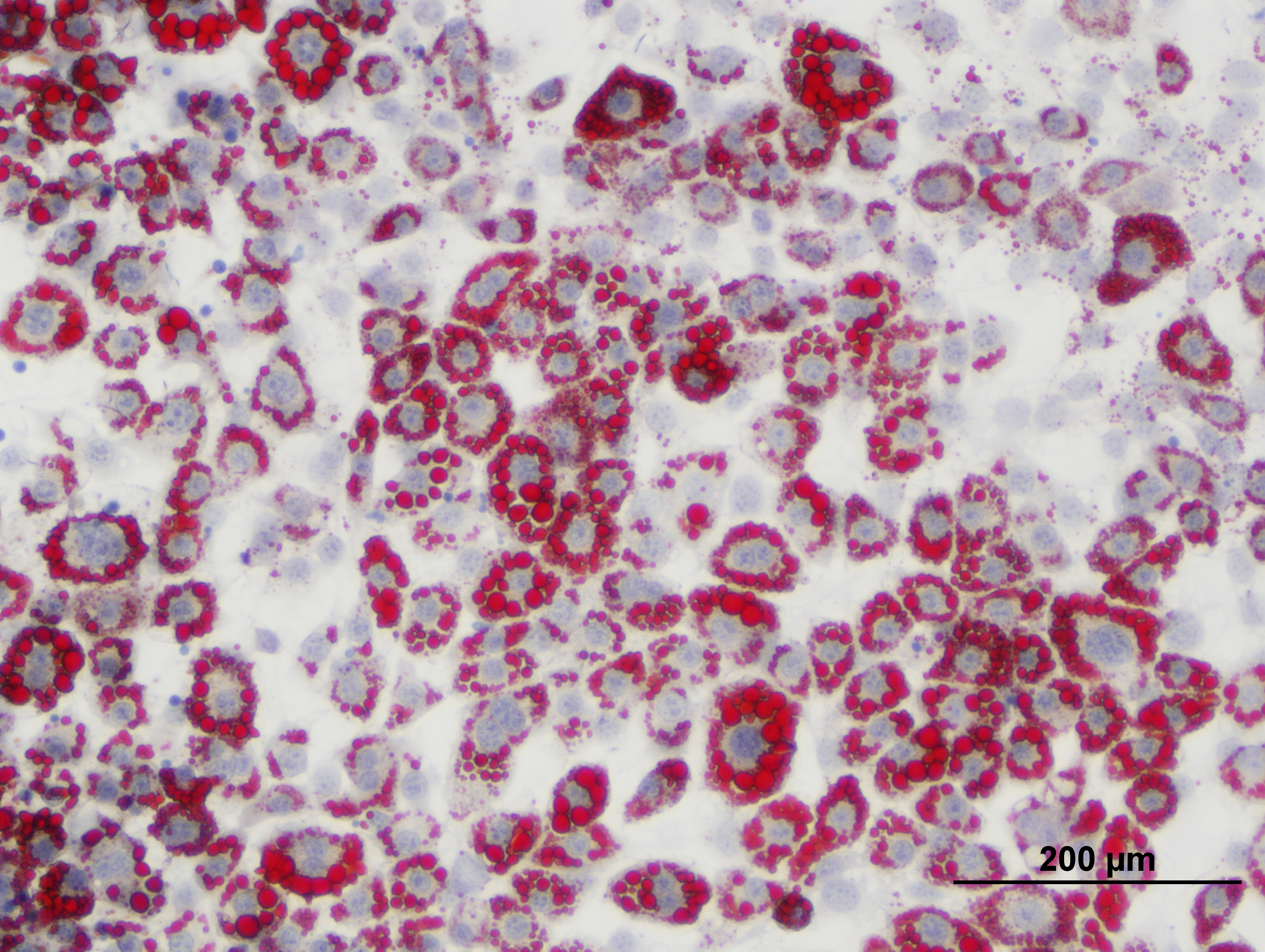

### 5 uM TXN+Rosi3.jpg

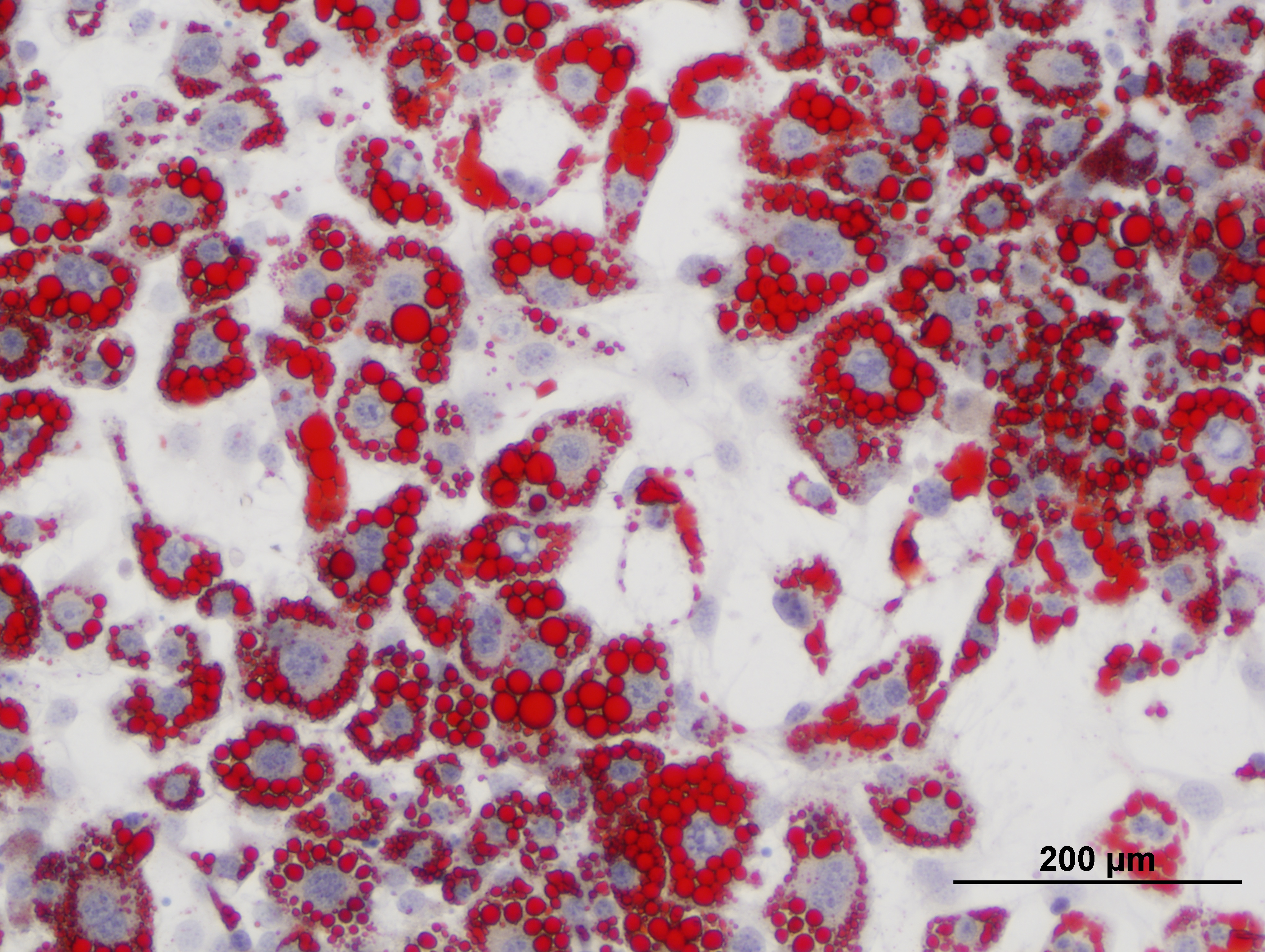

### 5 uM TXN+Rosi.jpg

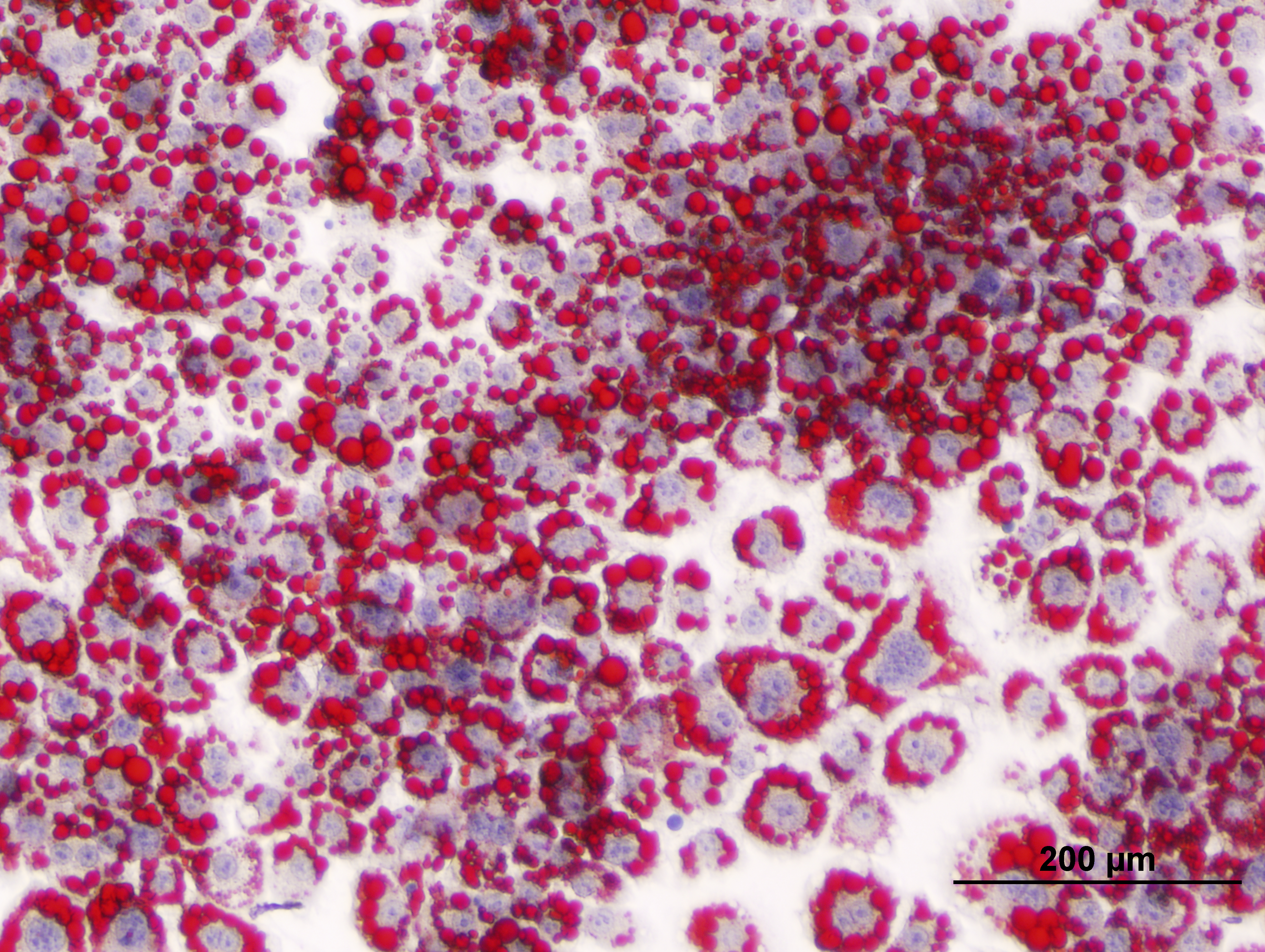

### 5 uM XN+Rosi2.jpg

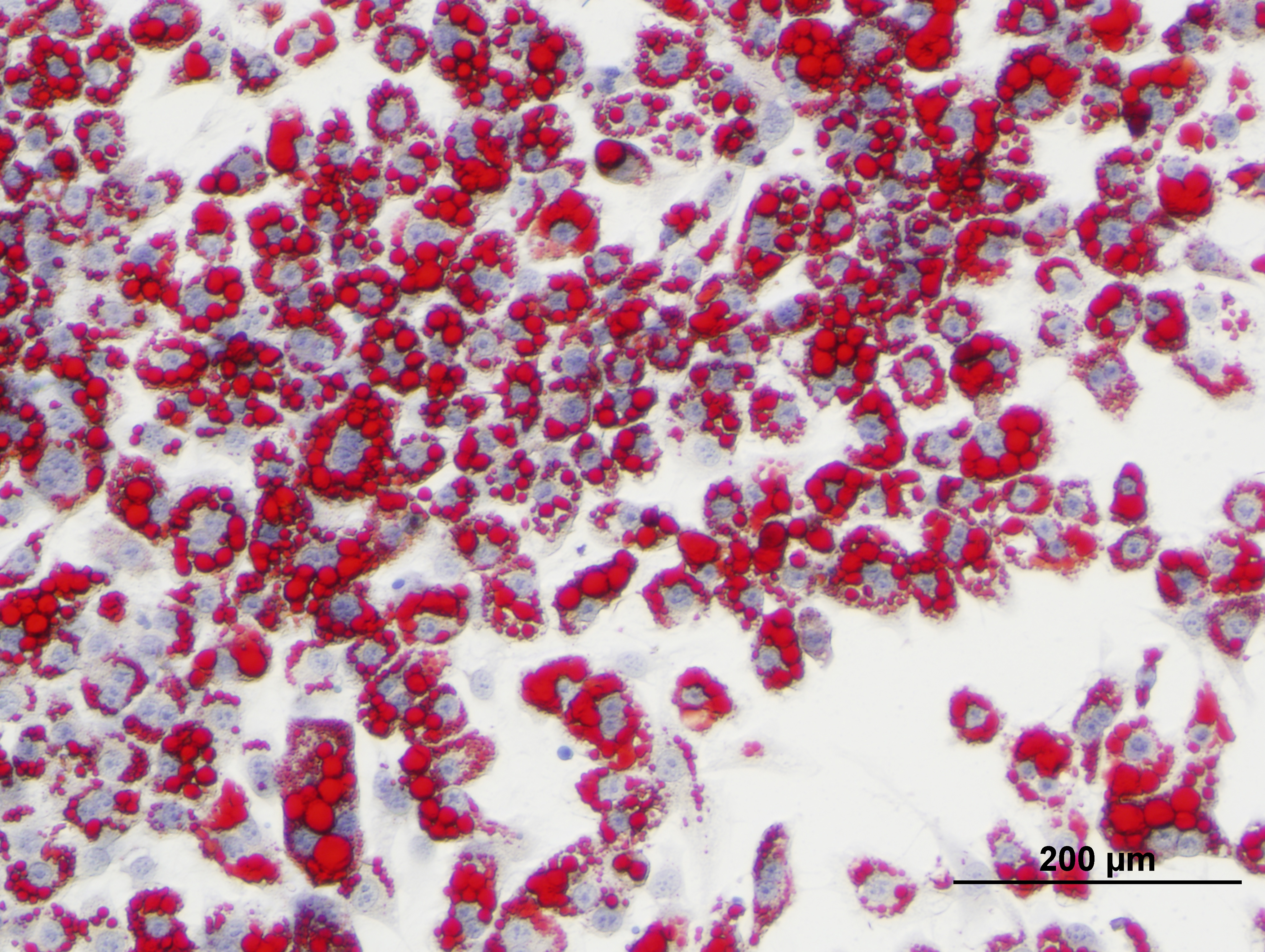

### 5 uM XN+Rosi3.jpg

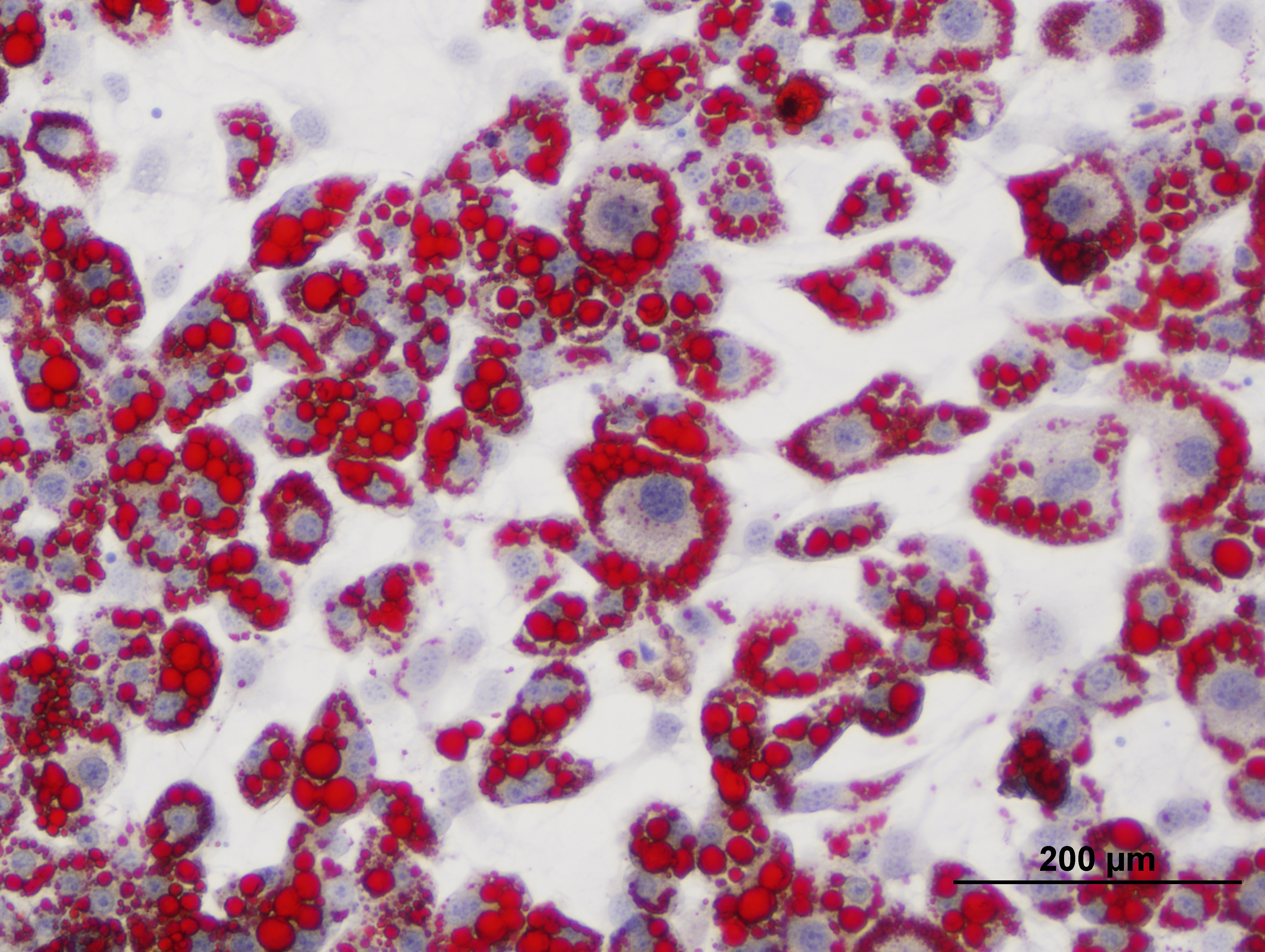

### 5 uM XN+Rosi.jpg

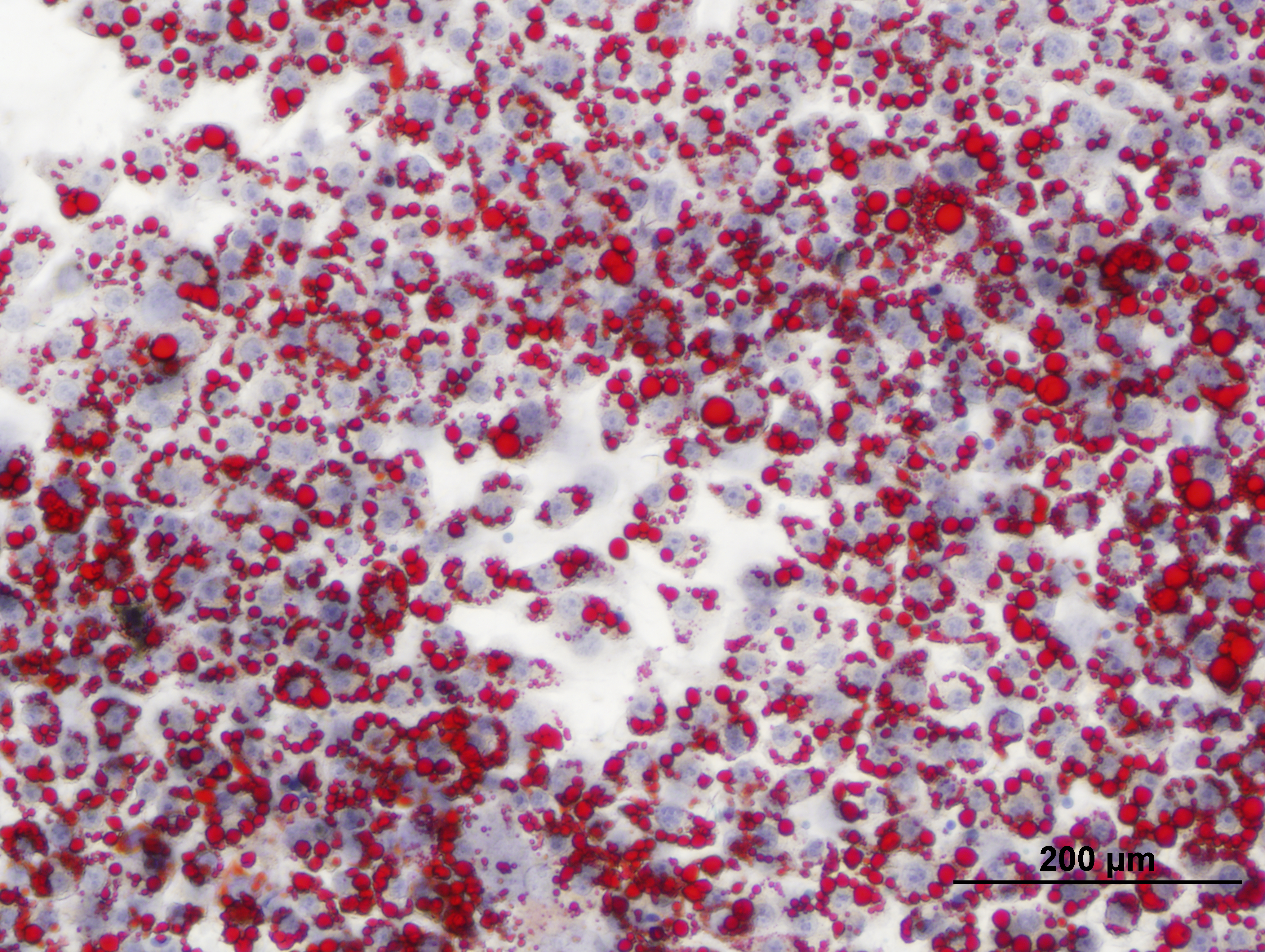

### 10 uM TXN+Rosi2.jpg

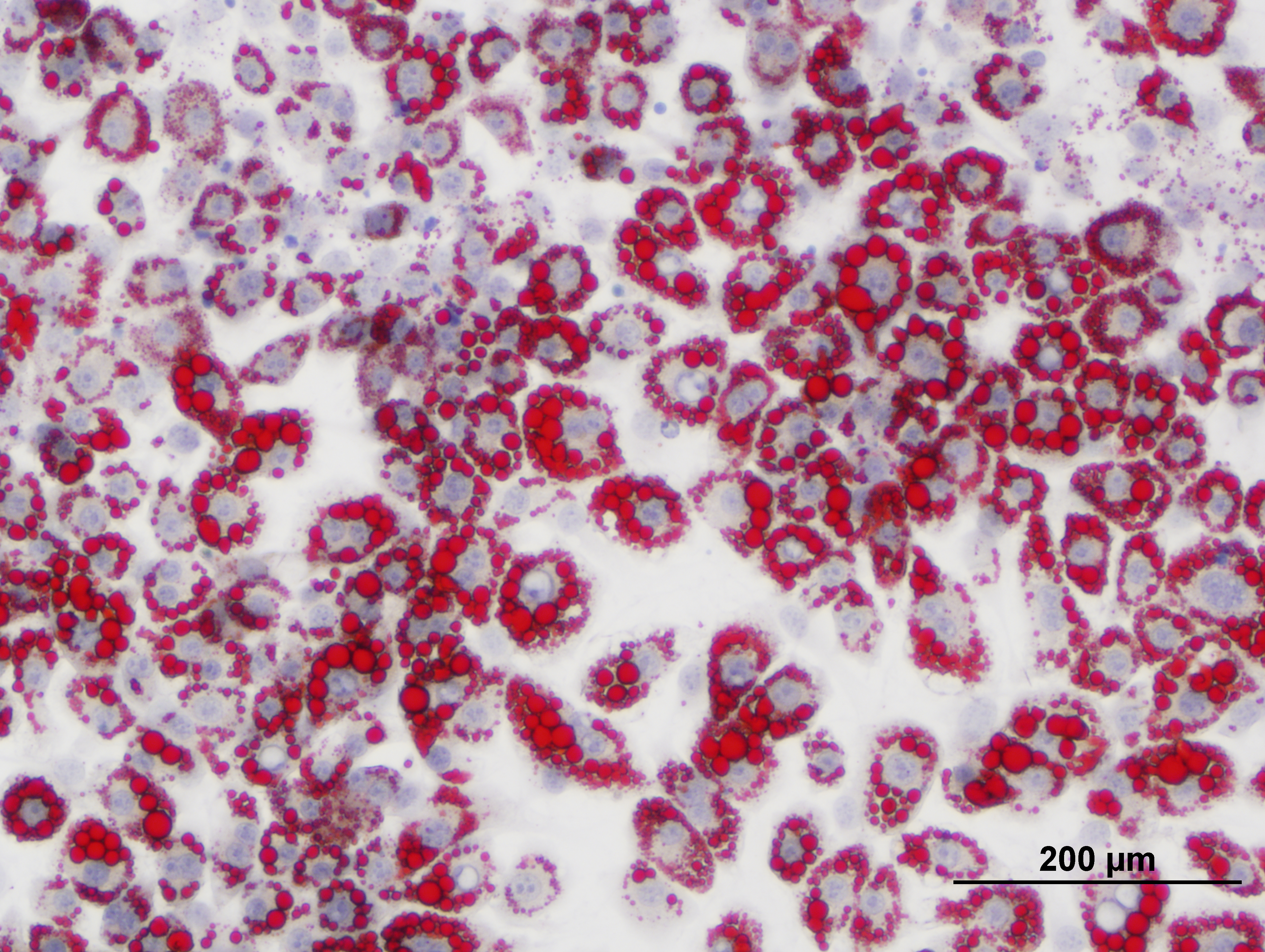

### 10 uM TXN+Rosi3.jpg

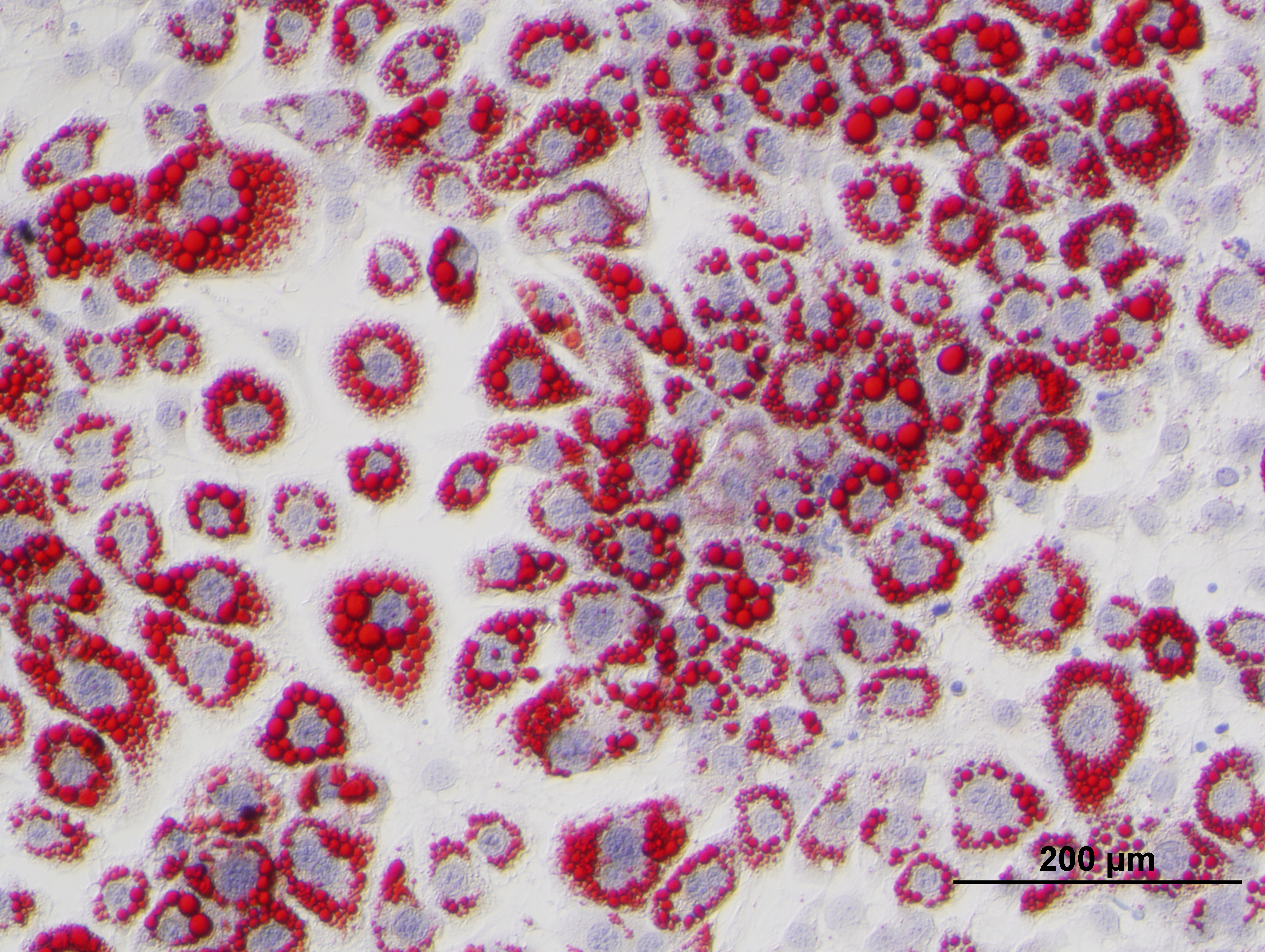

### 10 uM TXN+Rosi.jpg

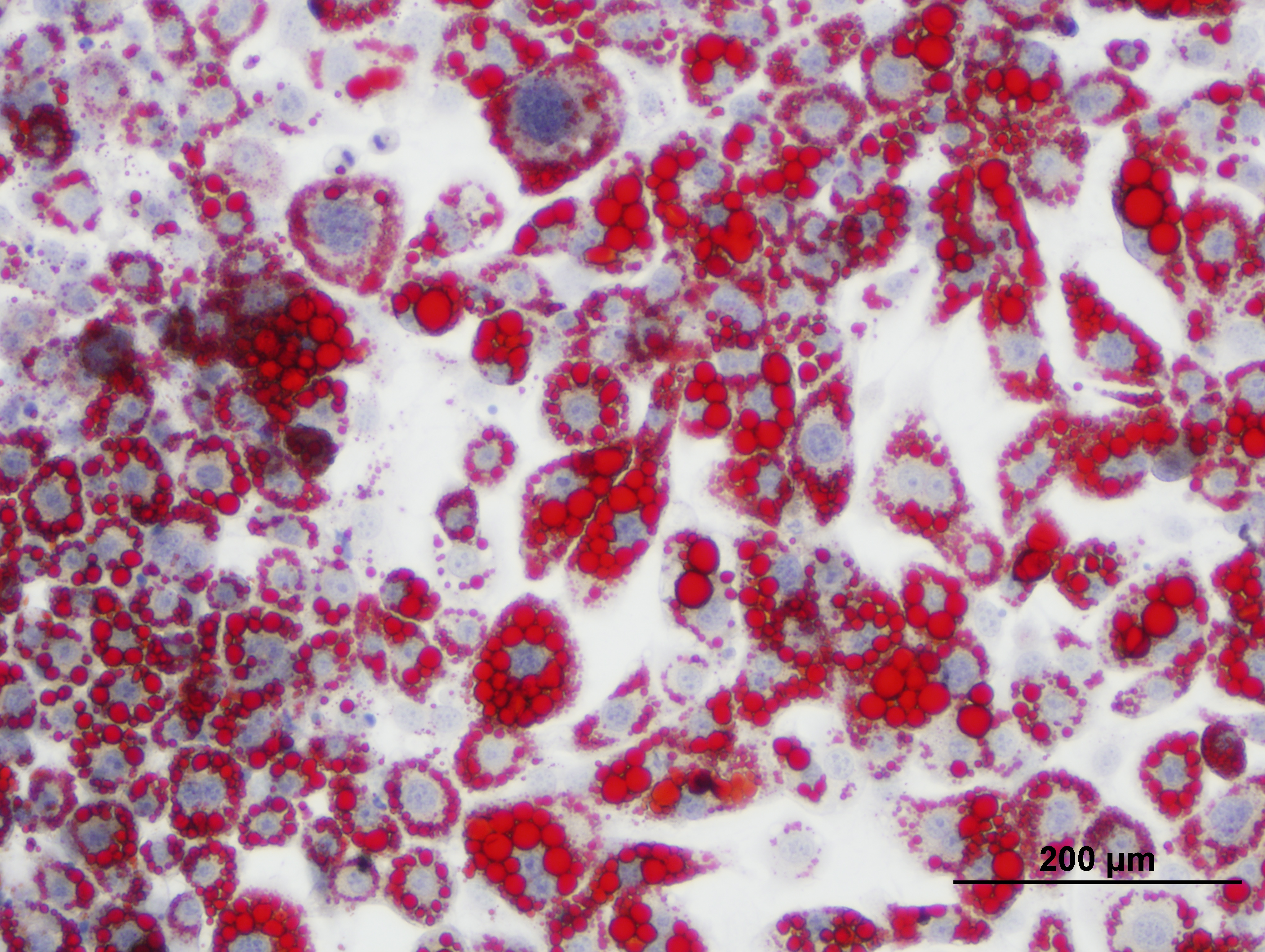

### 10 uM XN+Rosi2.jpg

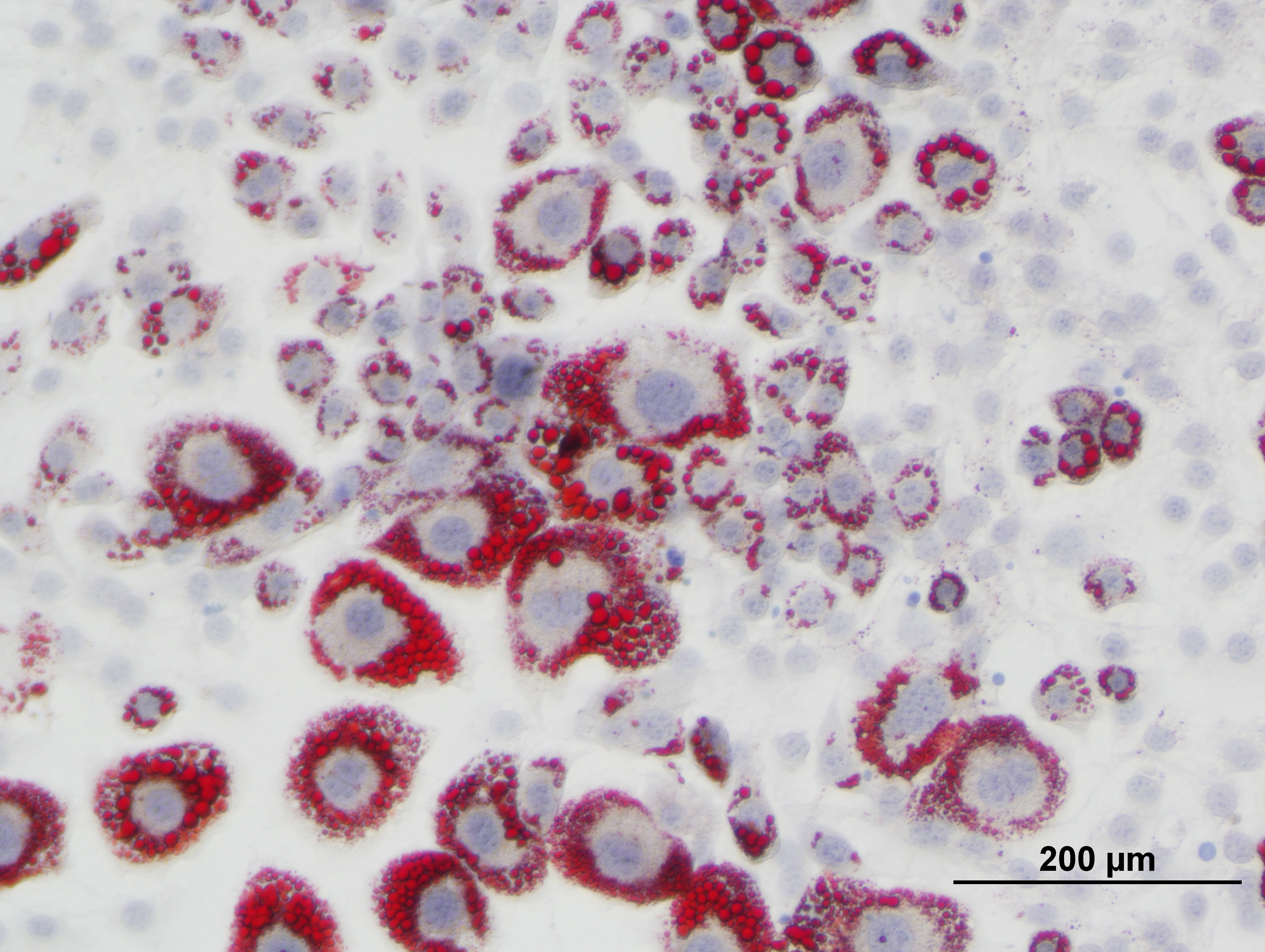

### 10 uM XN+Rosi3.jpg

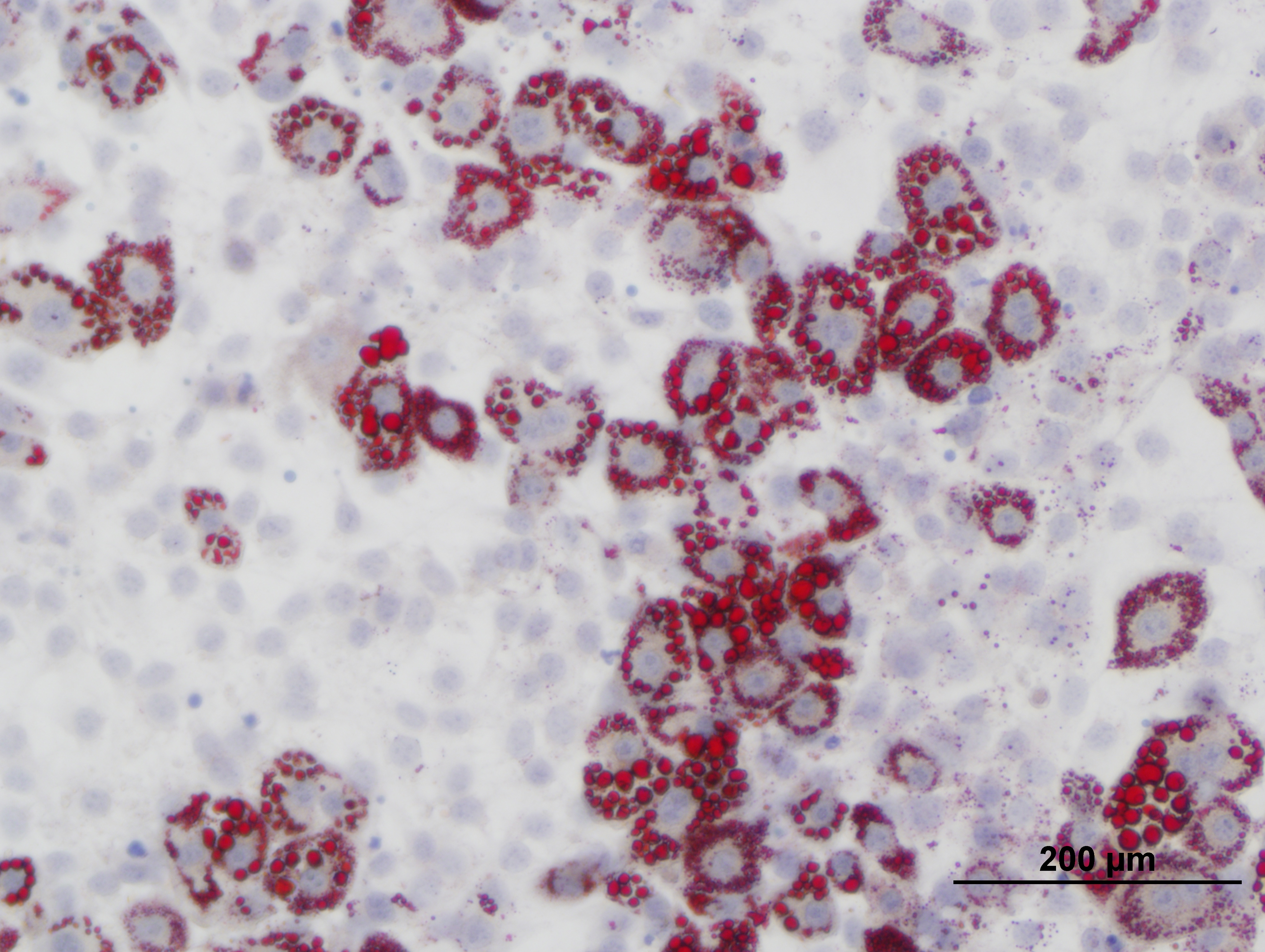

### 10 uM XN+Rosi.jpg

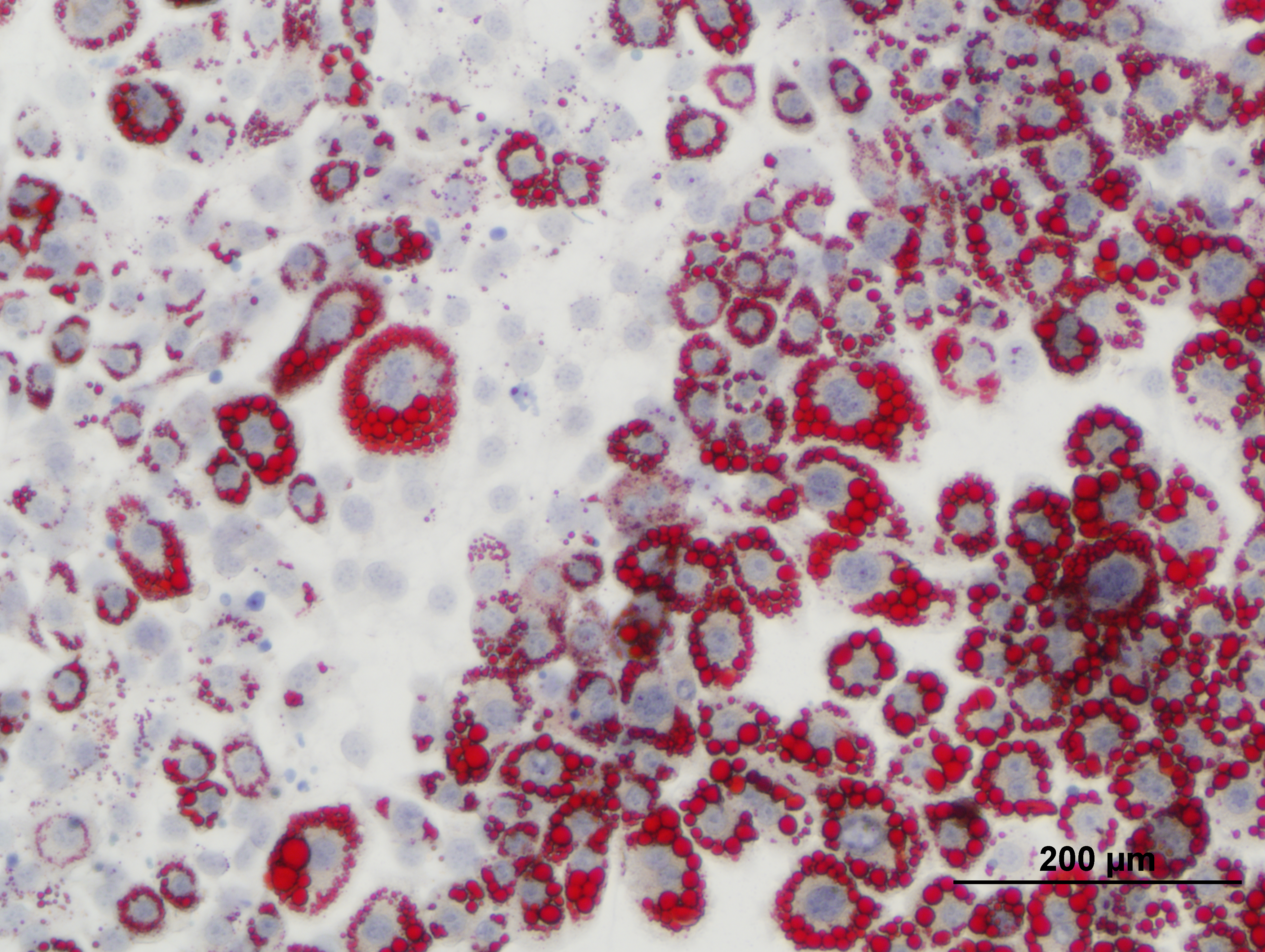

### 25 uM TXN+Rosi2.jpg

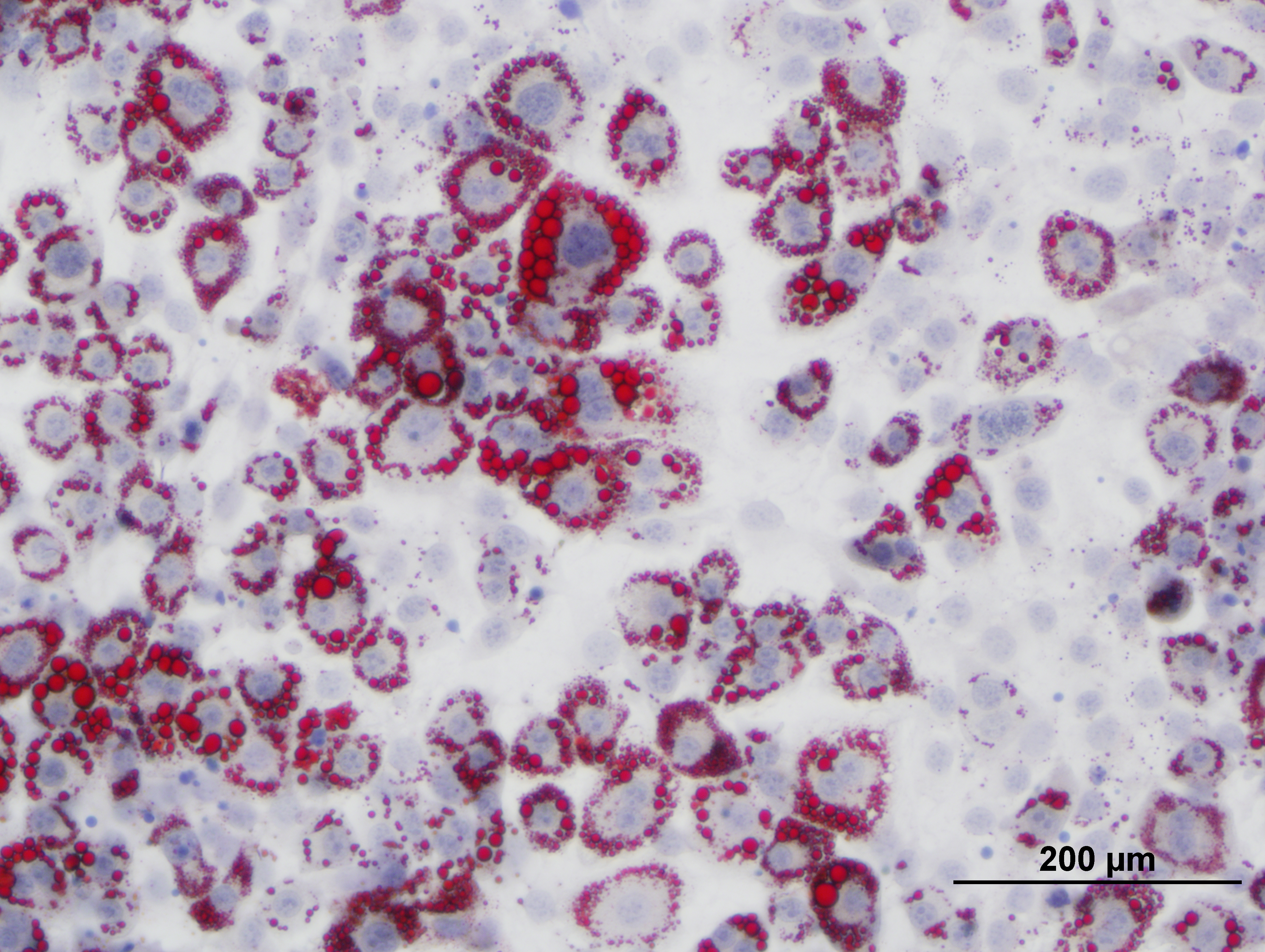

### 25 uM TXN+Rosi3.jpg

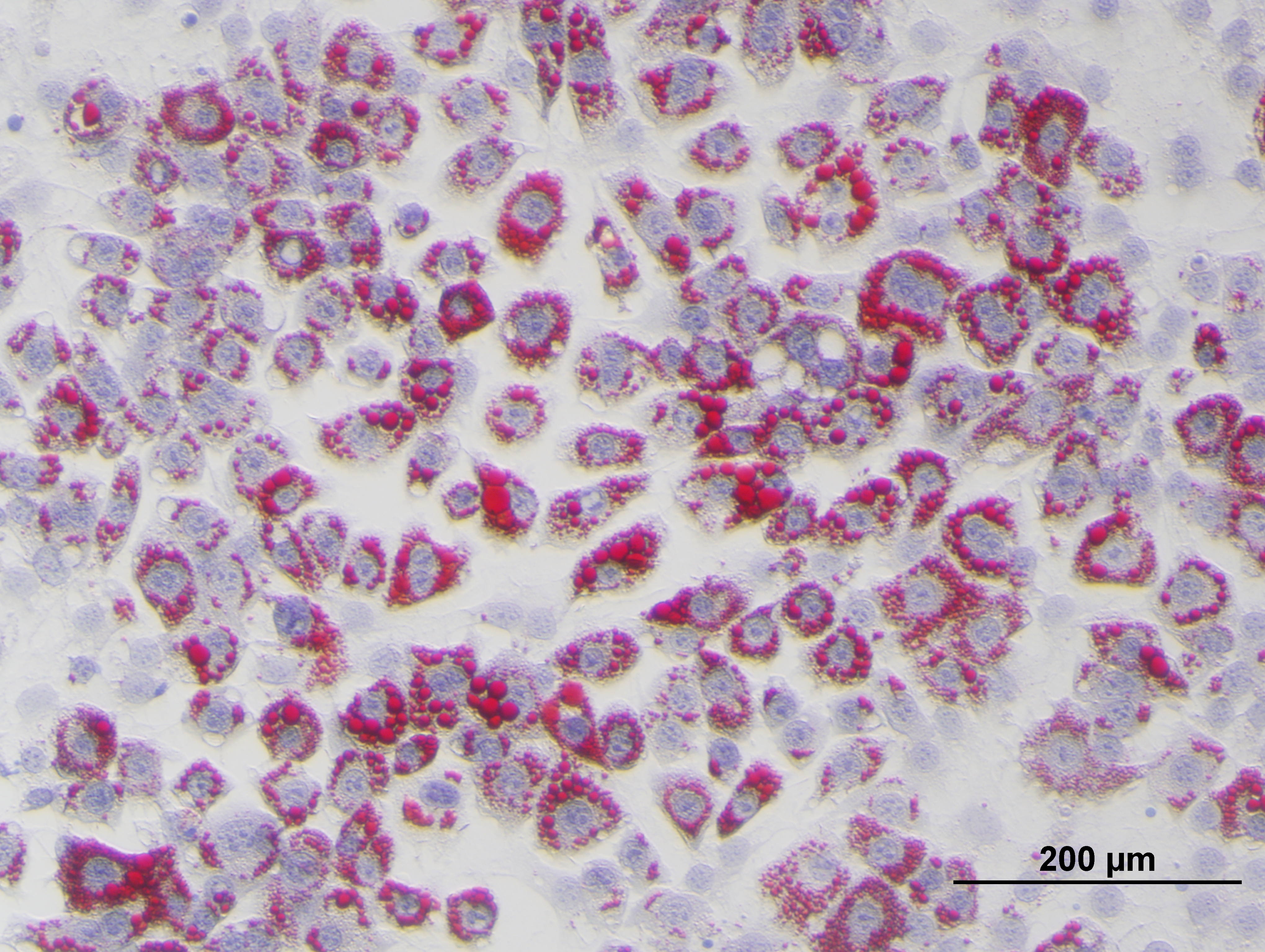

### 25 uM TXN+Rosi.jpg

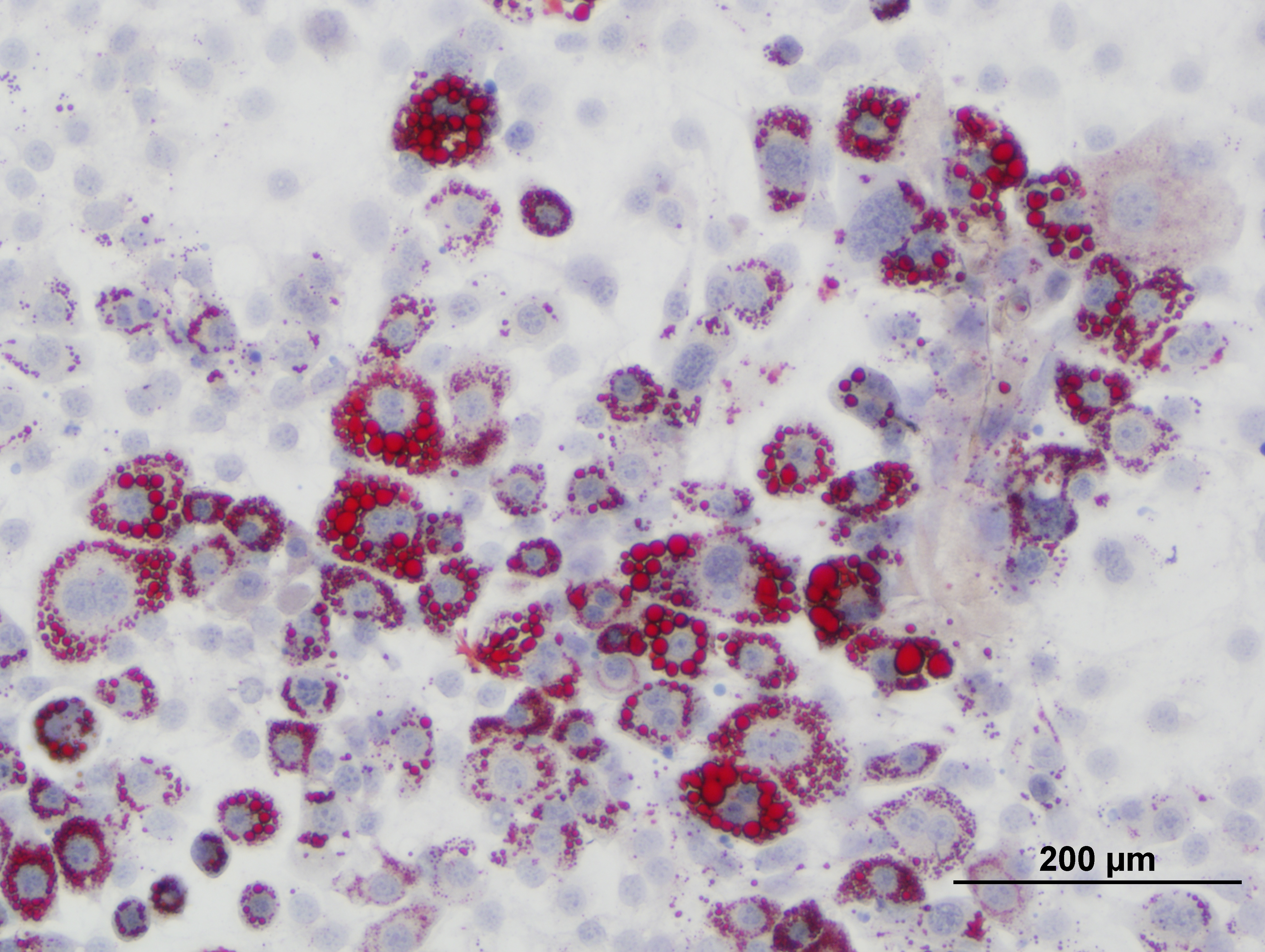

### 25 uM XN+Rosi2.jpg

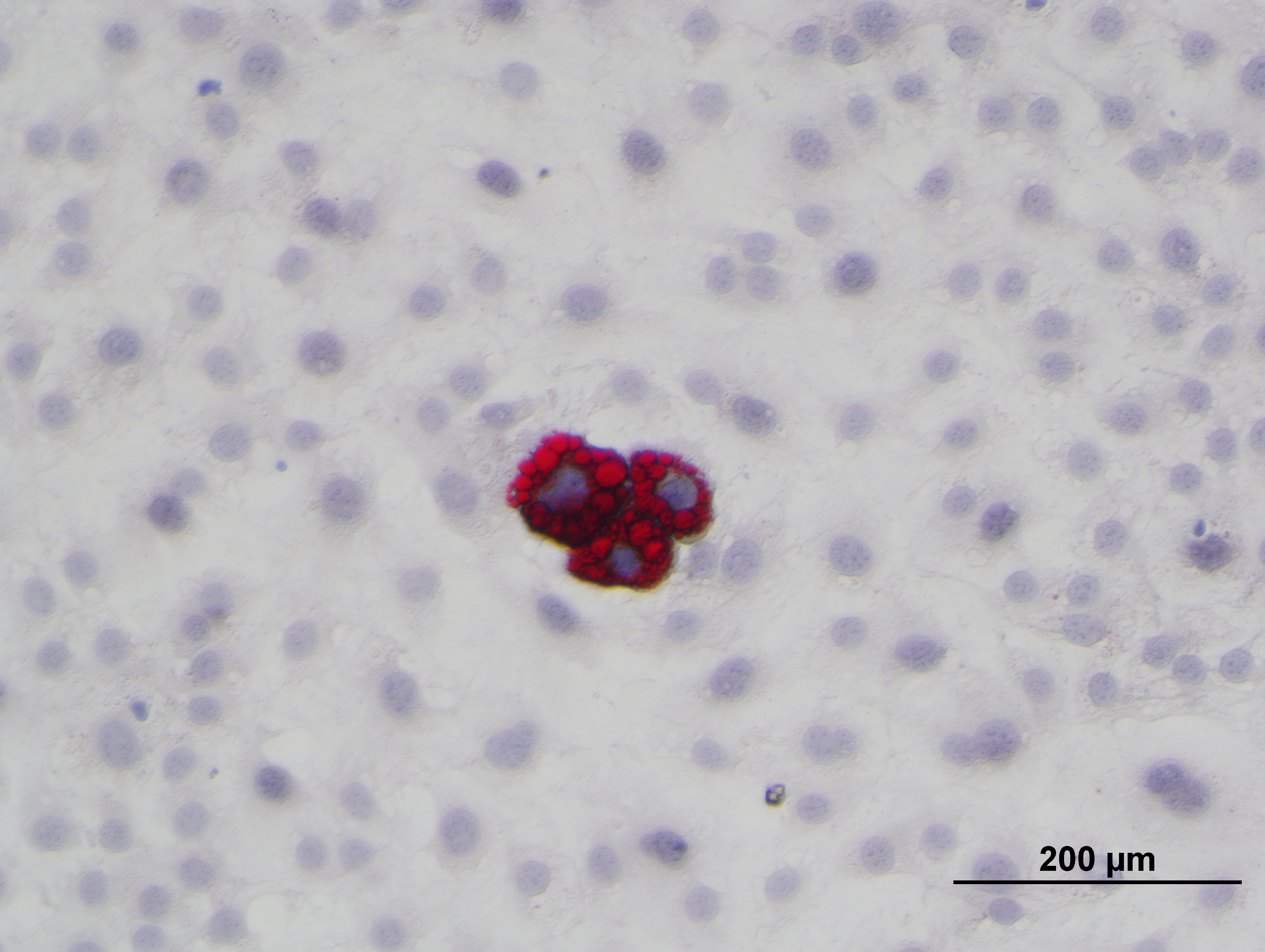

### 25 uM XN+Rosi3.jpg

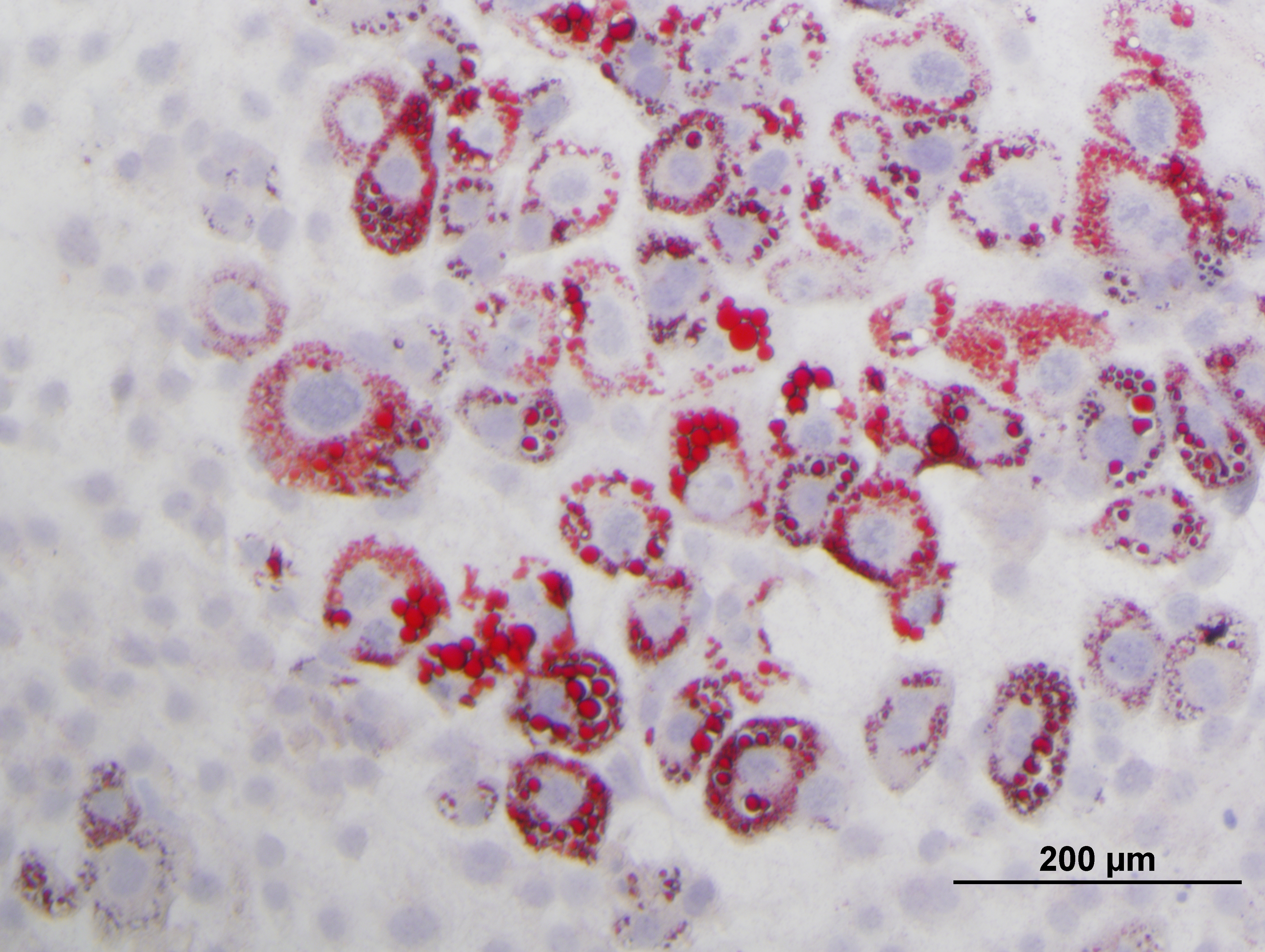

### 25 uM XN+Rosi.jpg

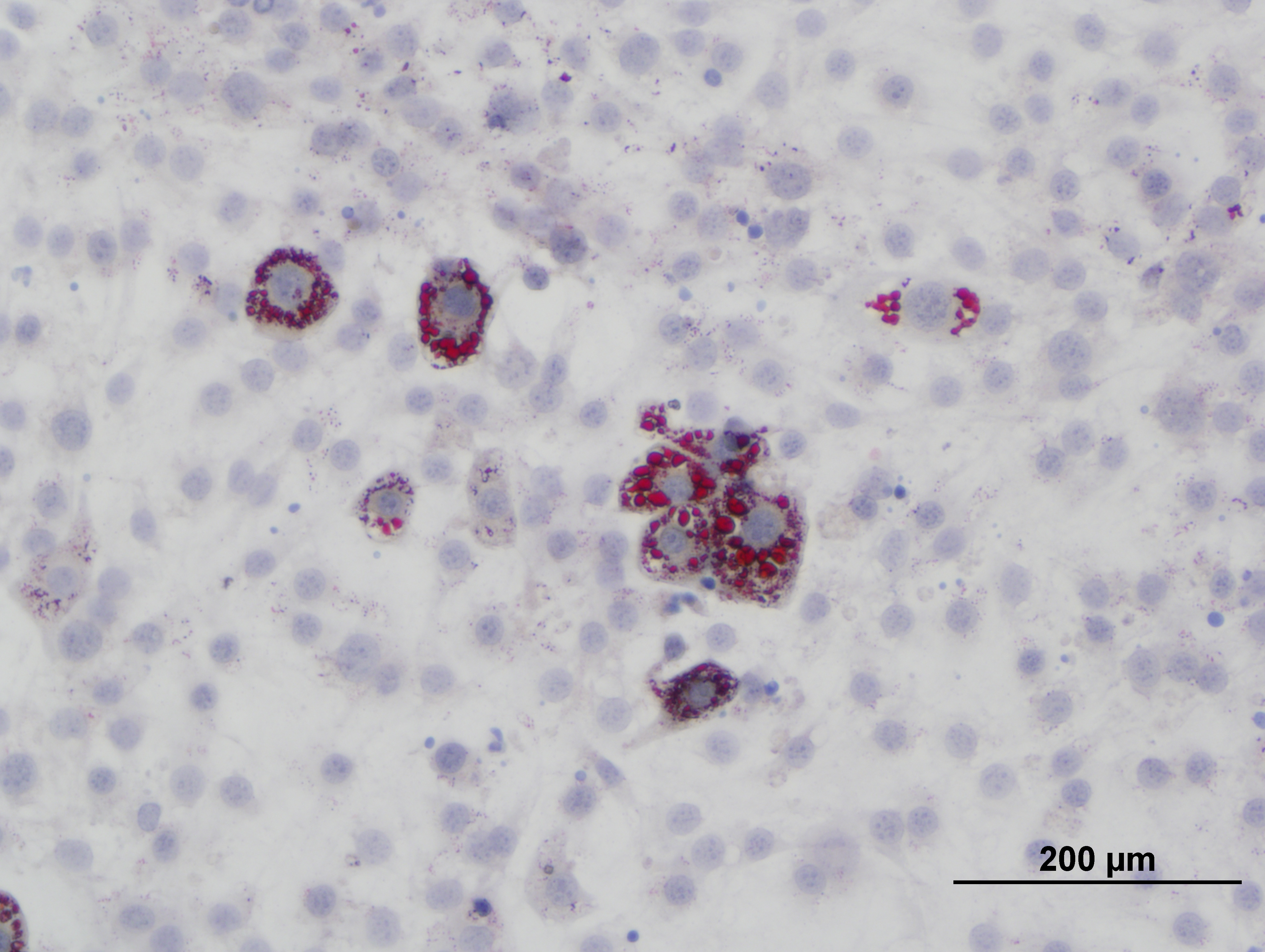

### B.pdf

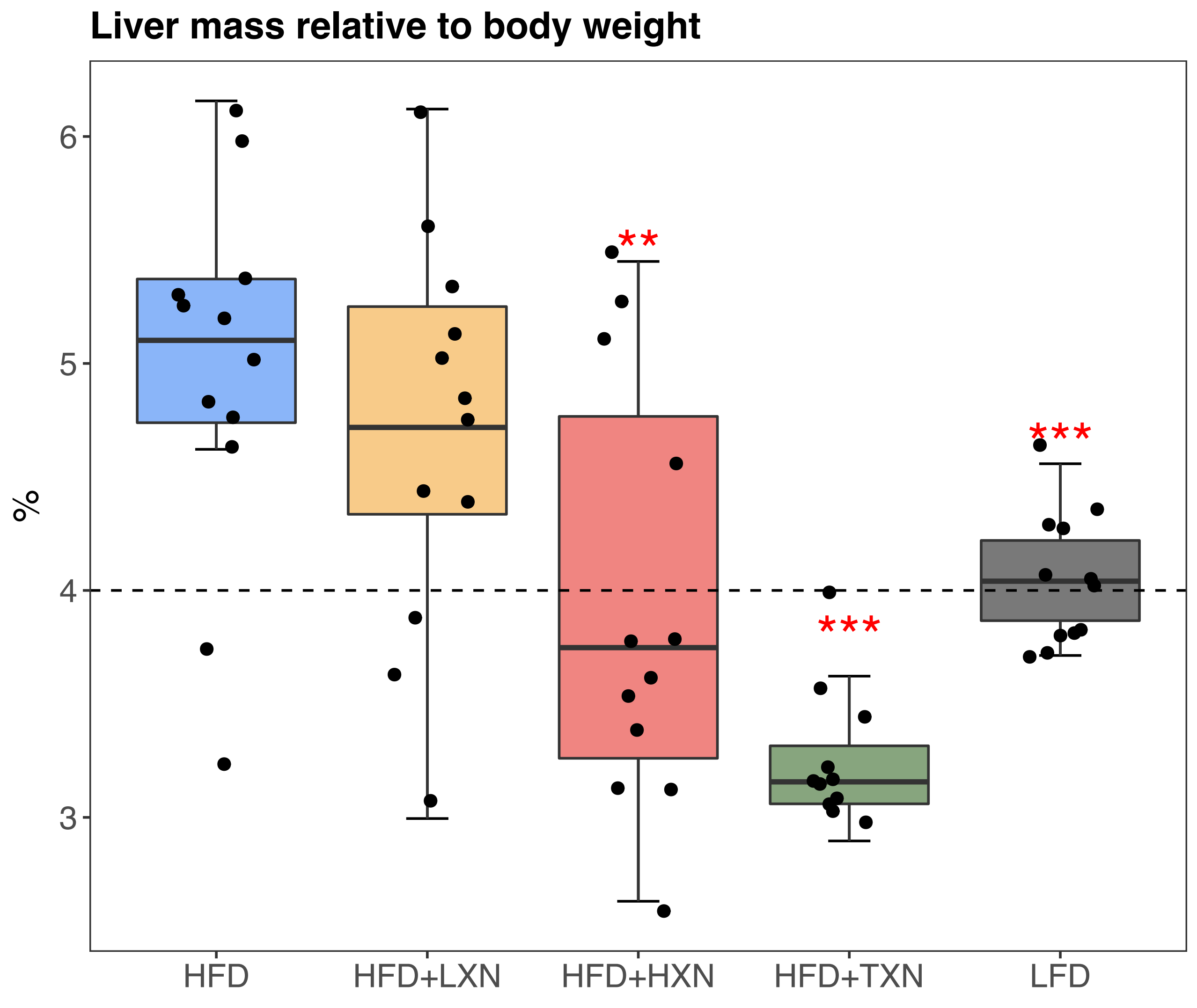

### C.pdf

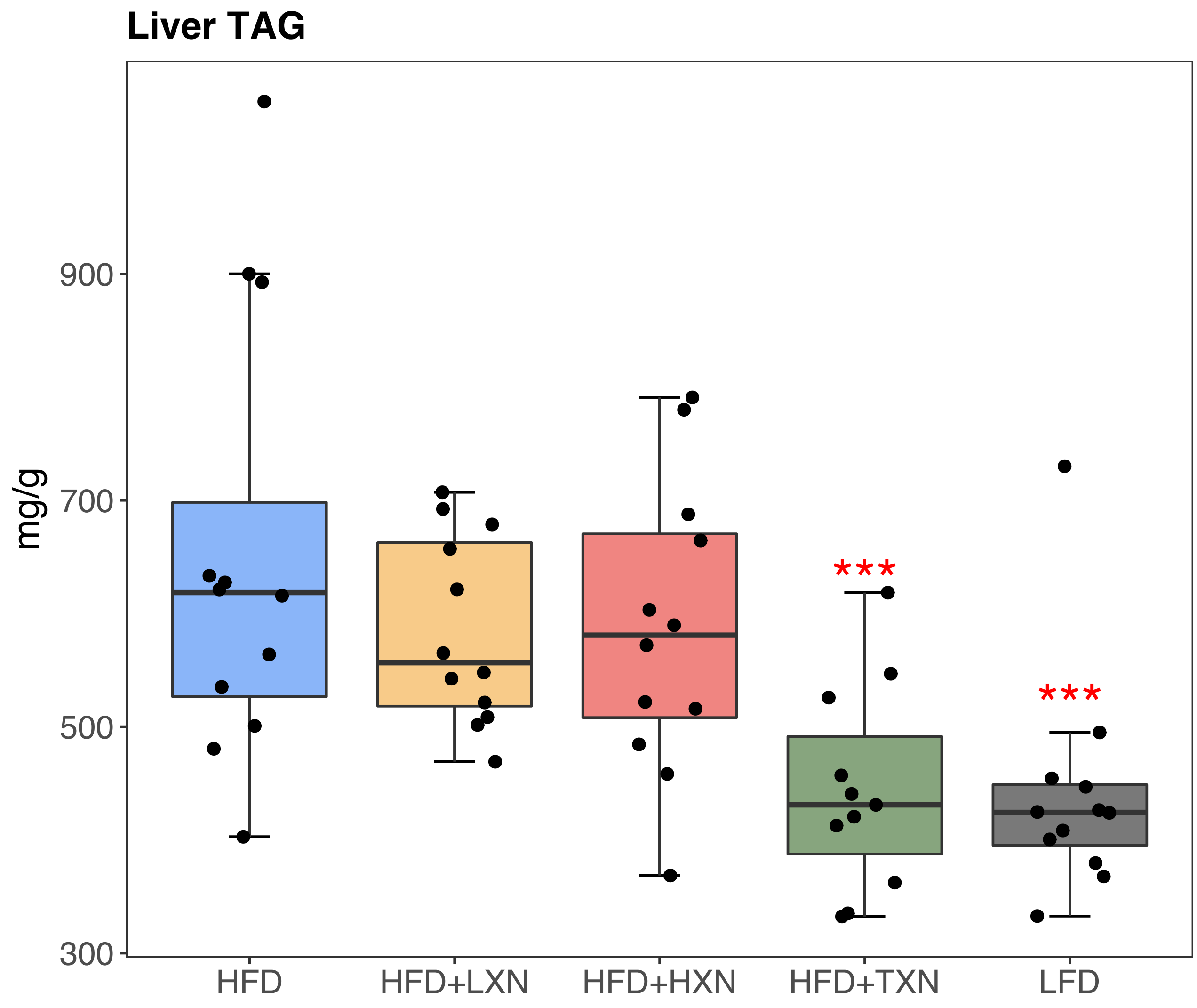

### fig3Sup1.pdf

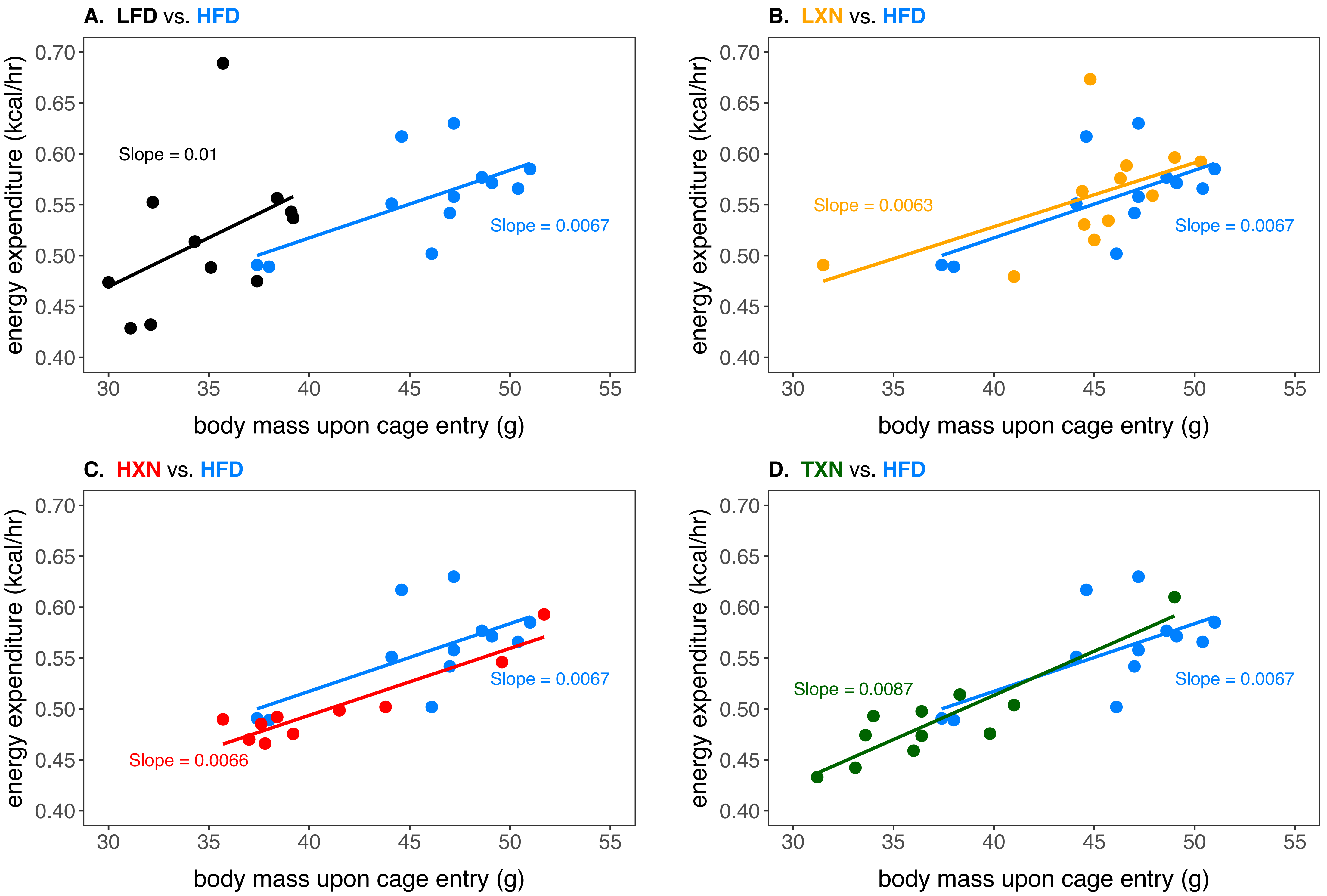

### fig3Sup2.pdf

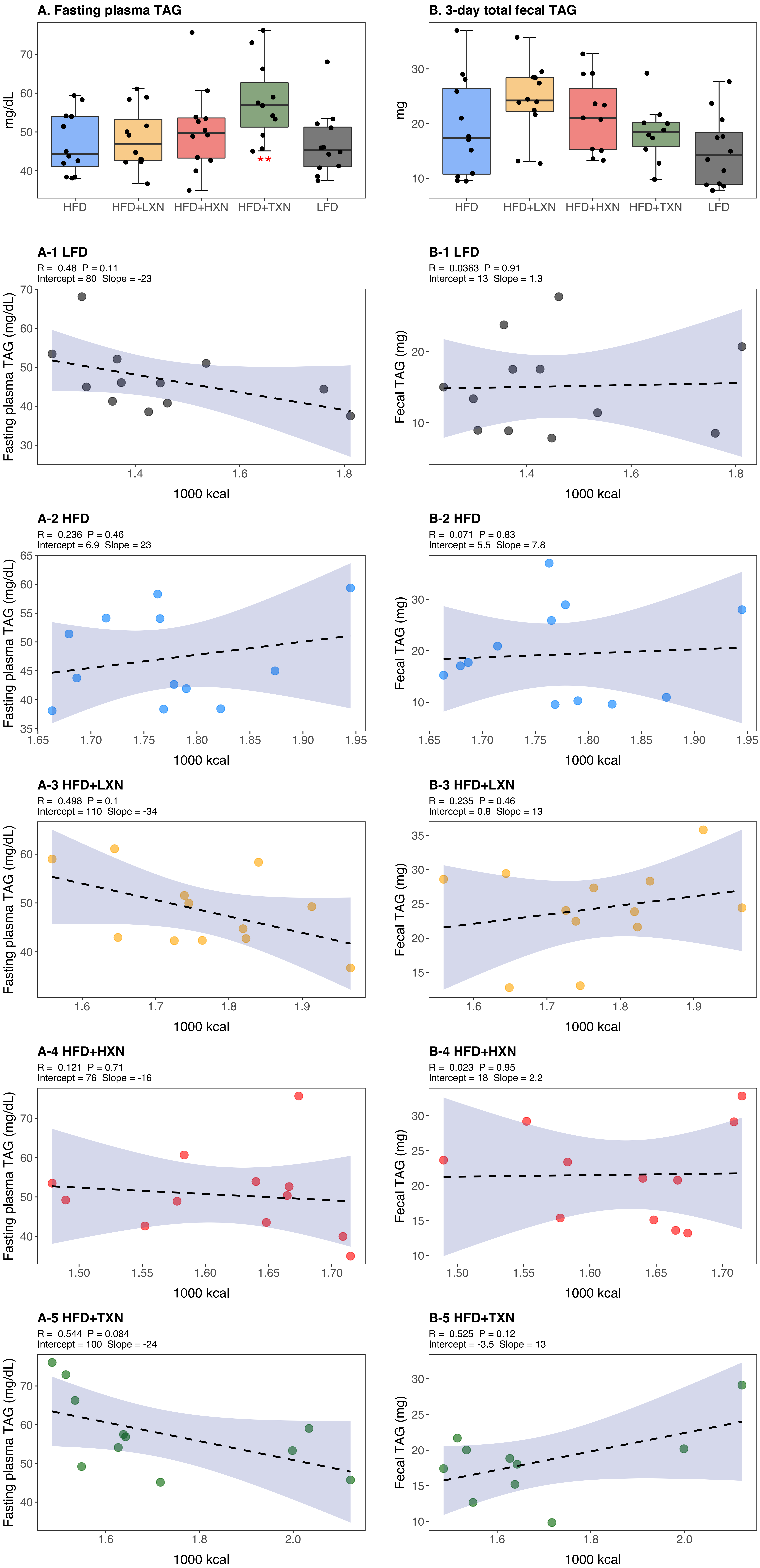

### fig9.pdf

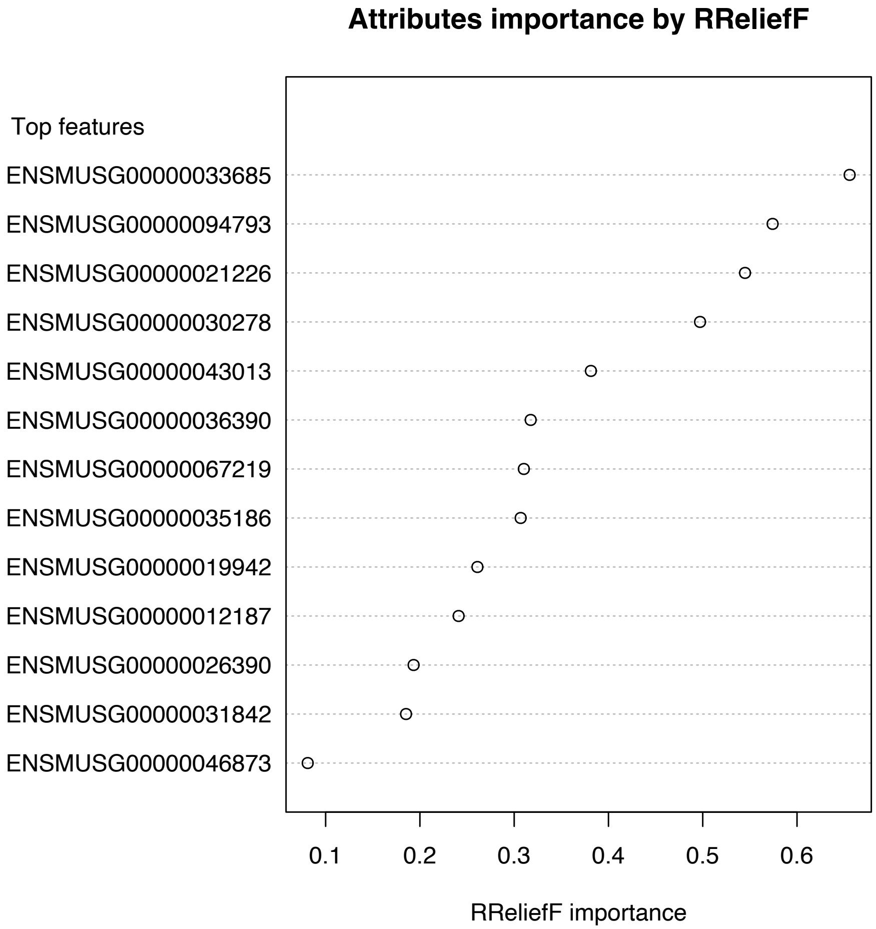

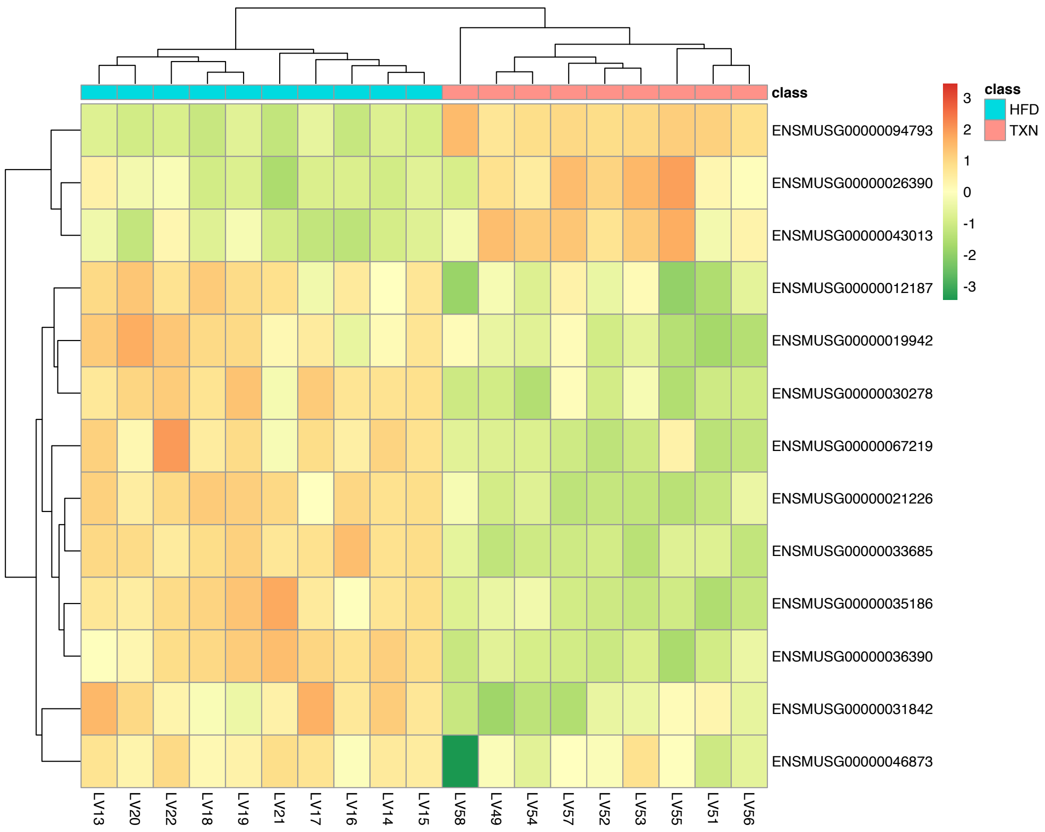

### fig9.pptx

#### Slide 1
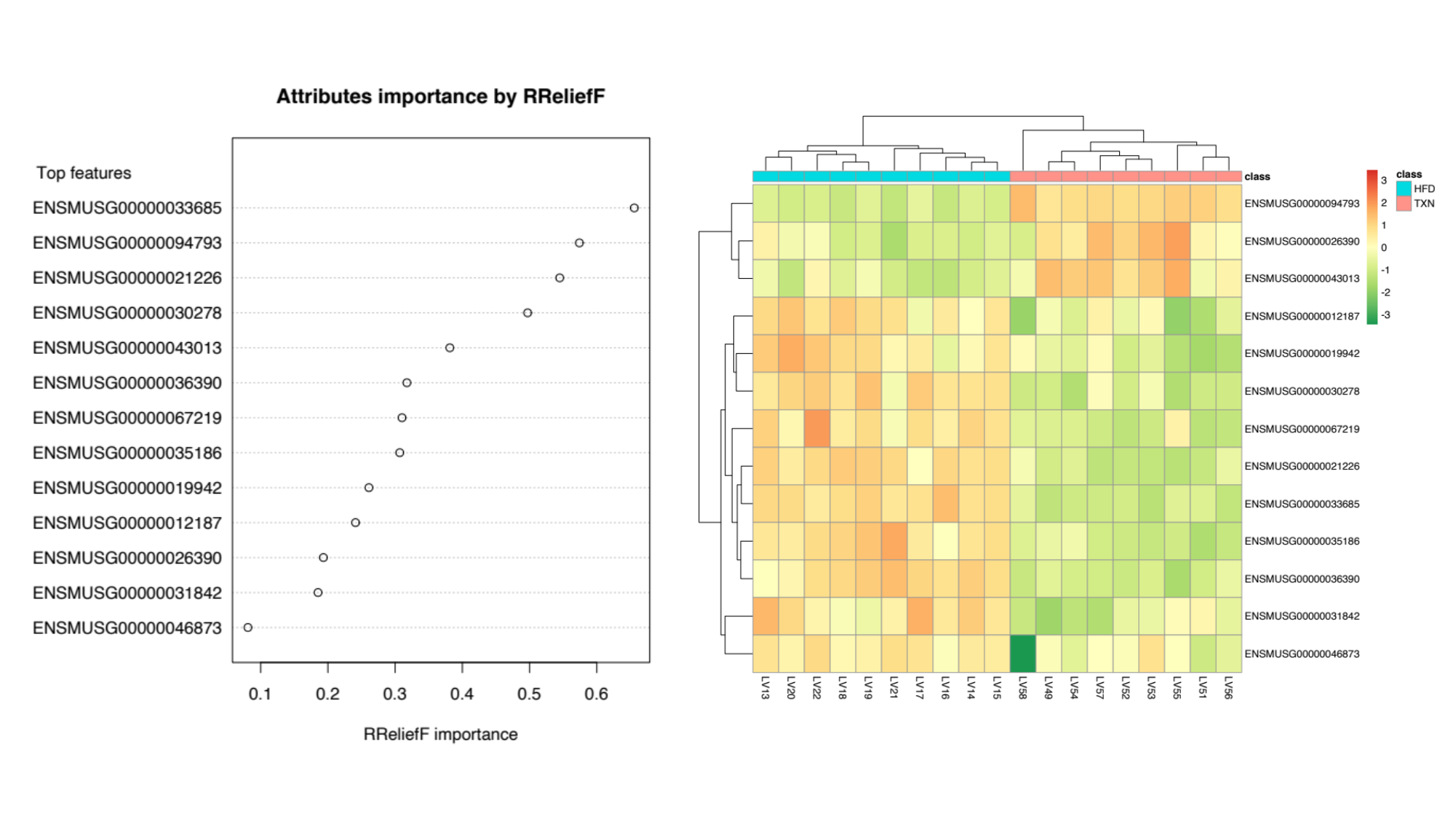

### figure10.pdf

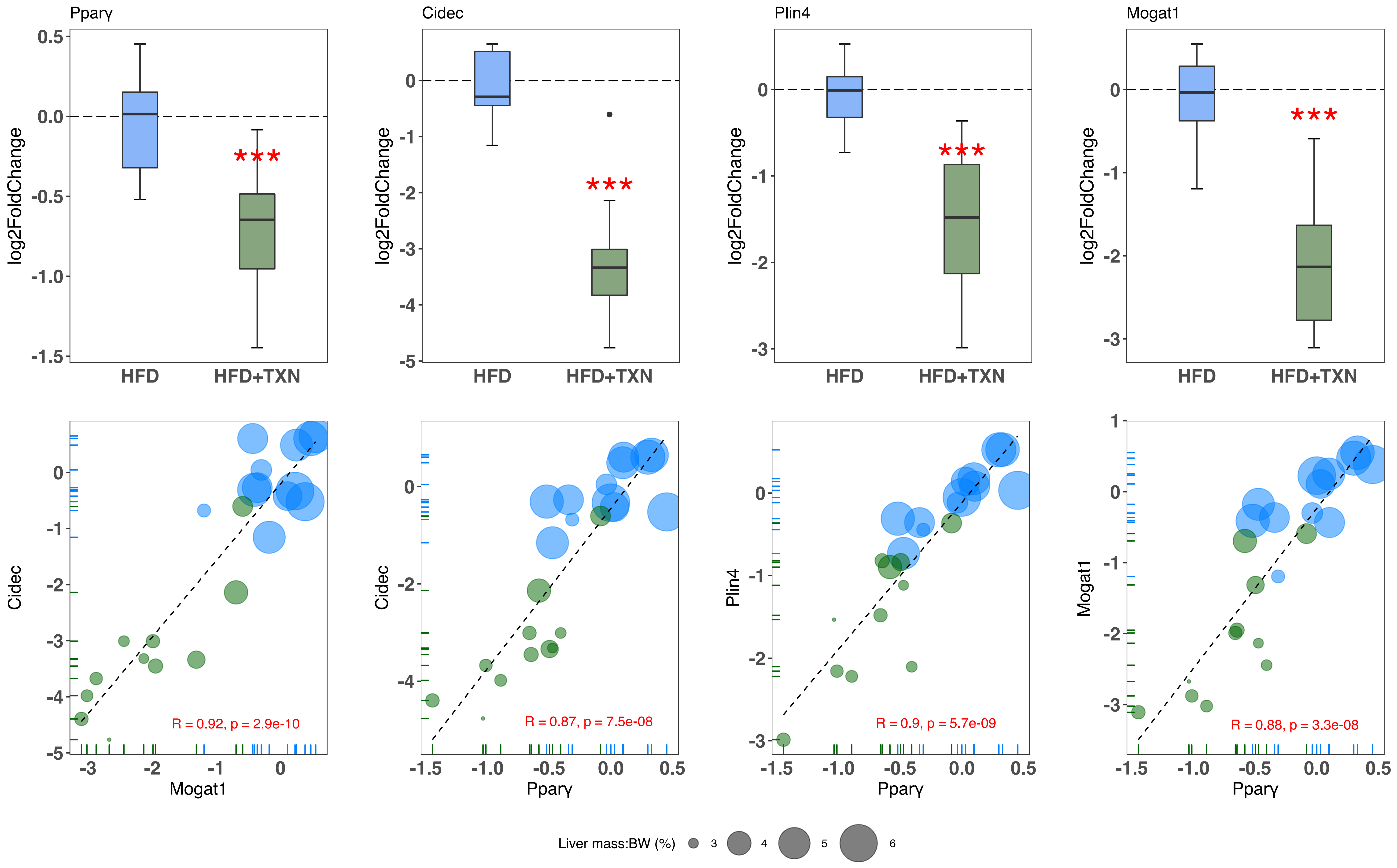

### GW2.jpg

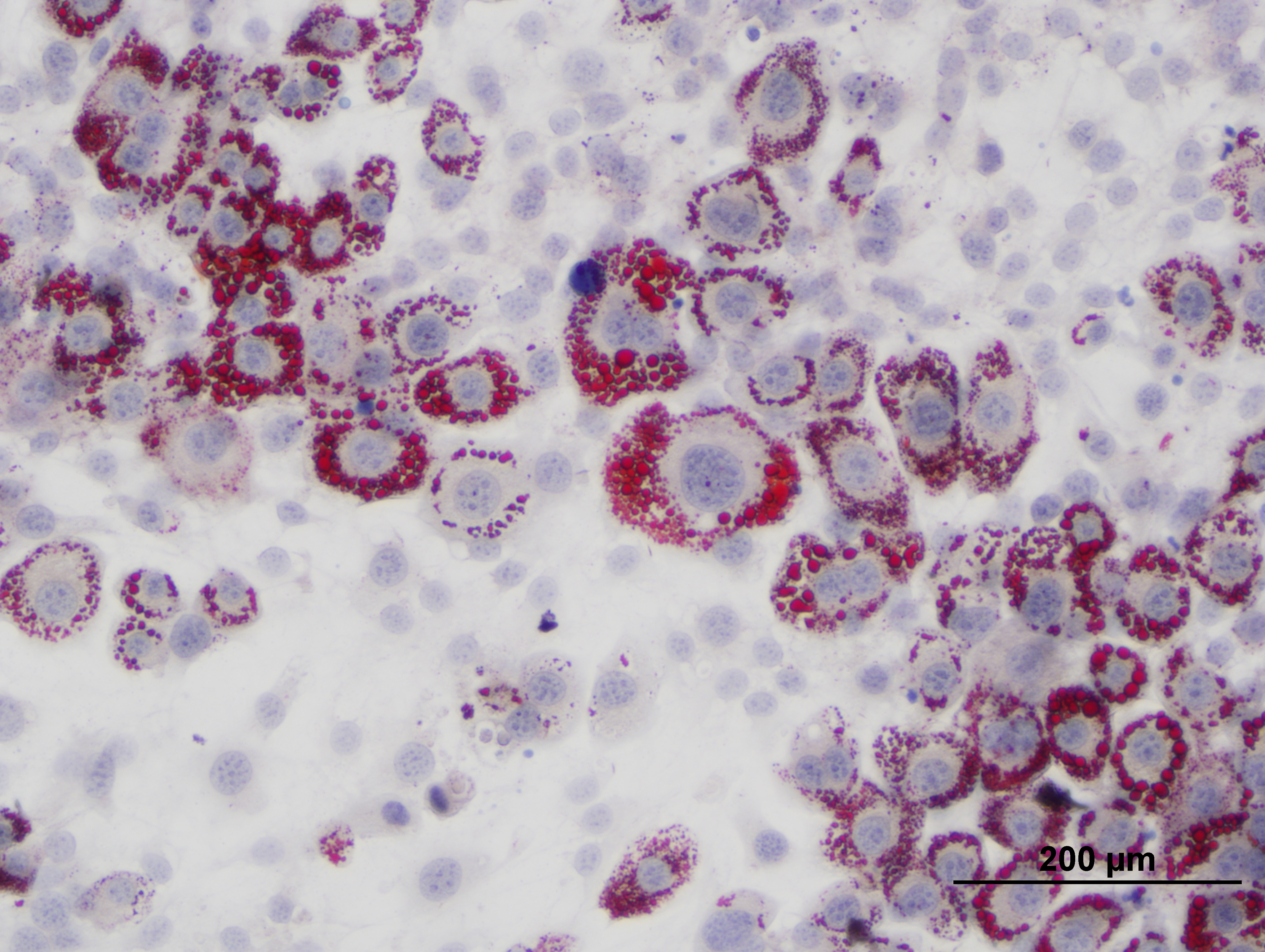

### GW.jpg

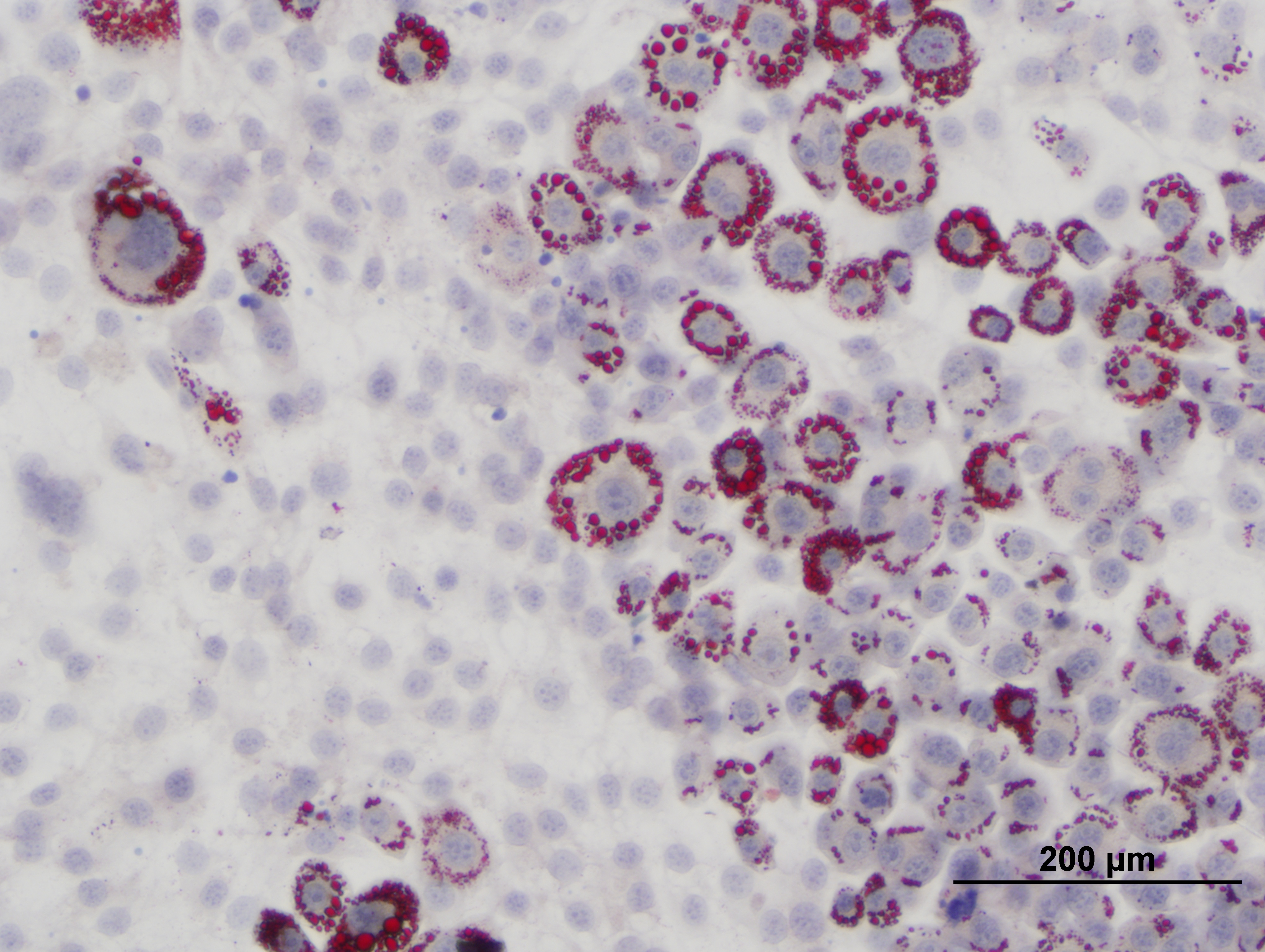

### heatmap.pdf

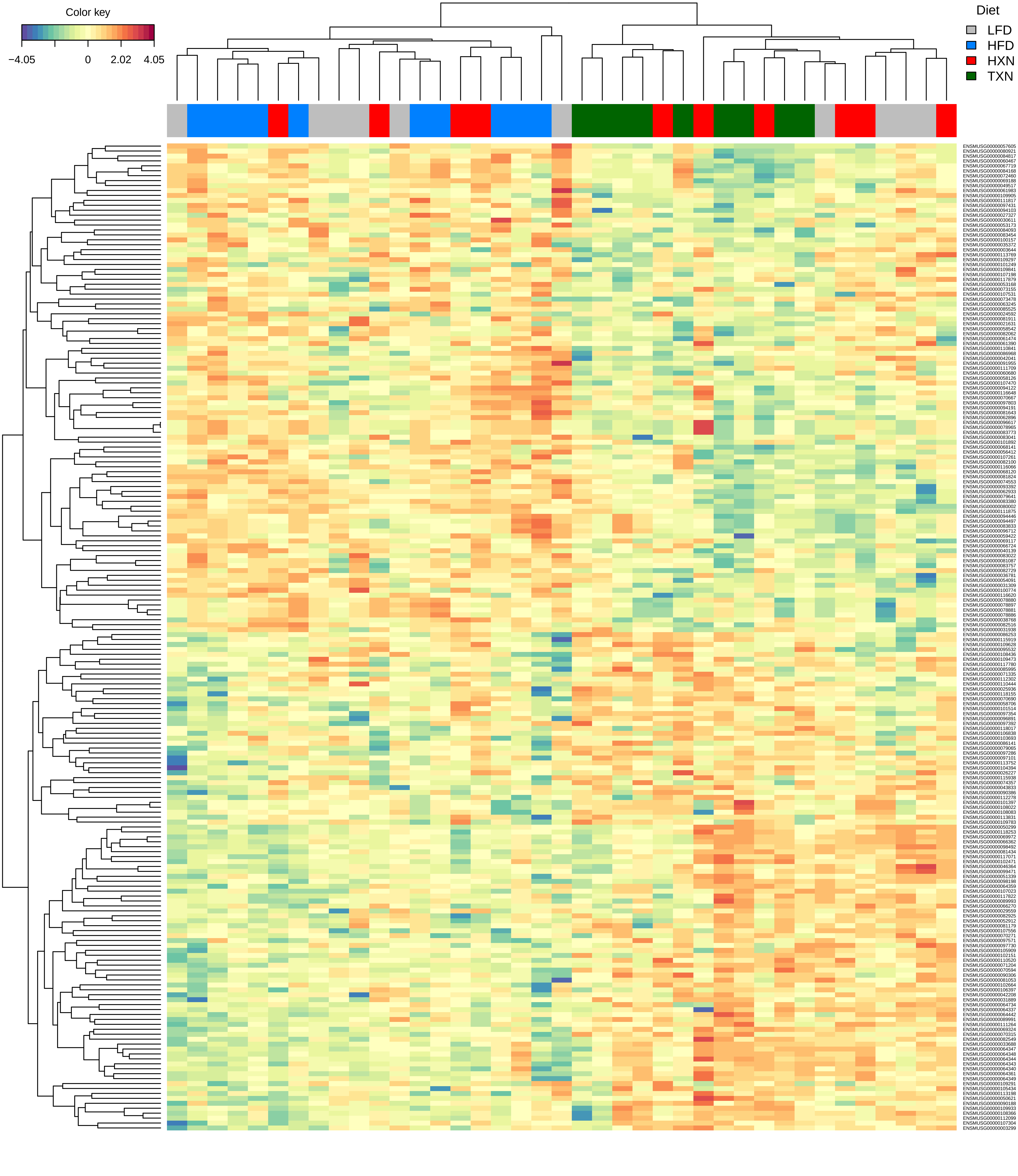

### leftPanel.pdf

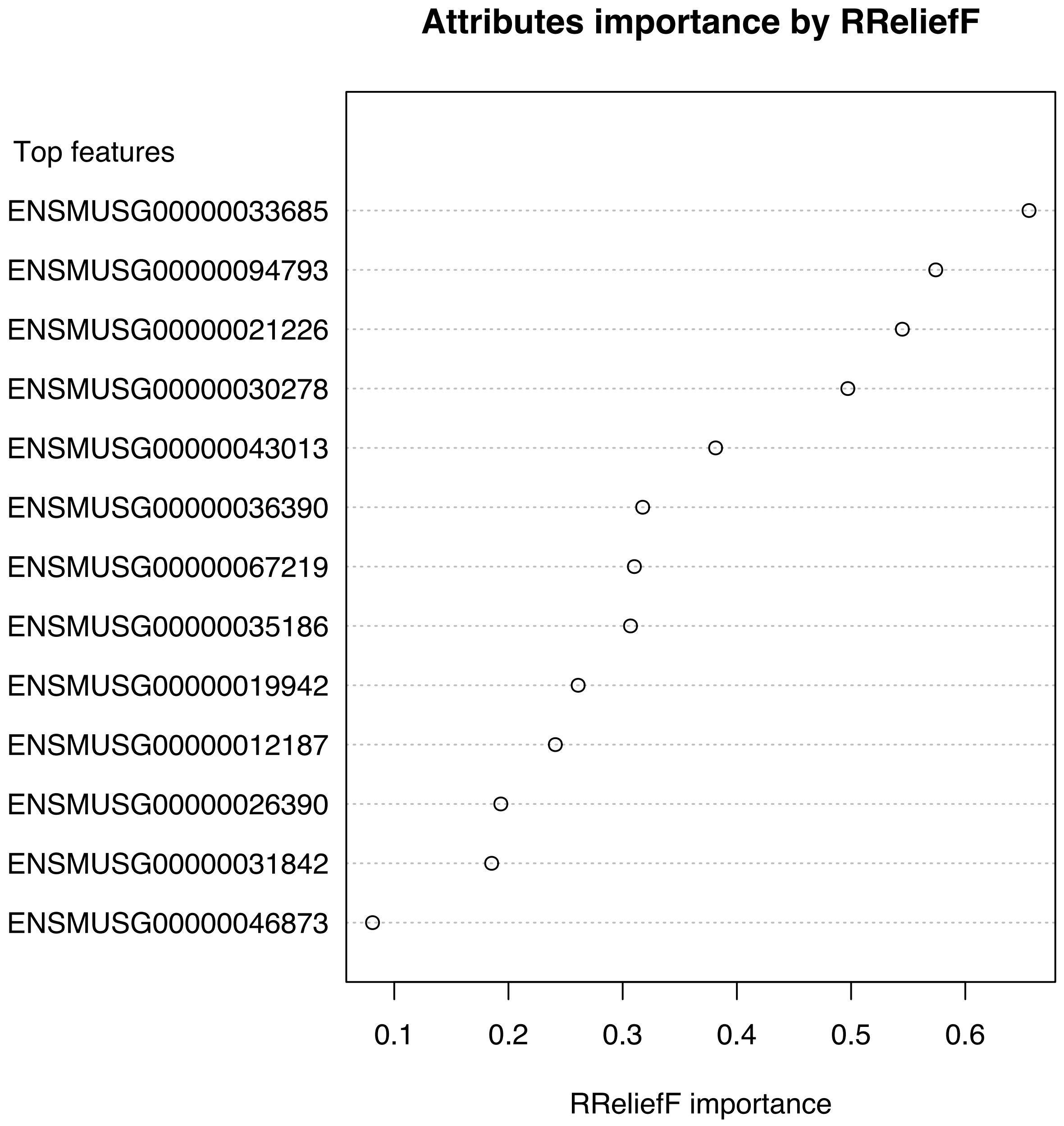

### rightPanel.pdf

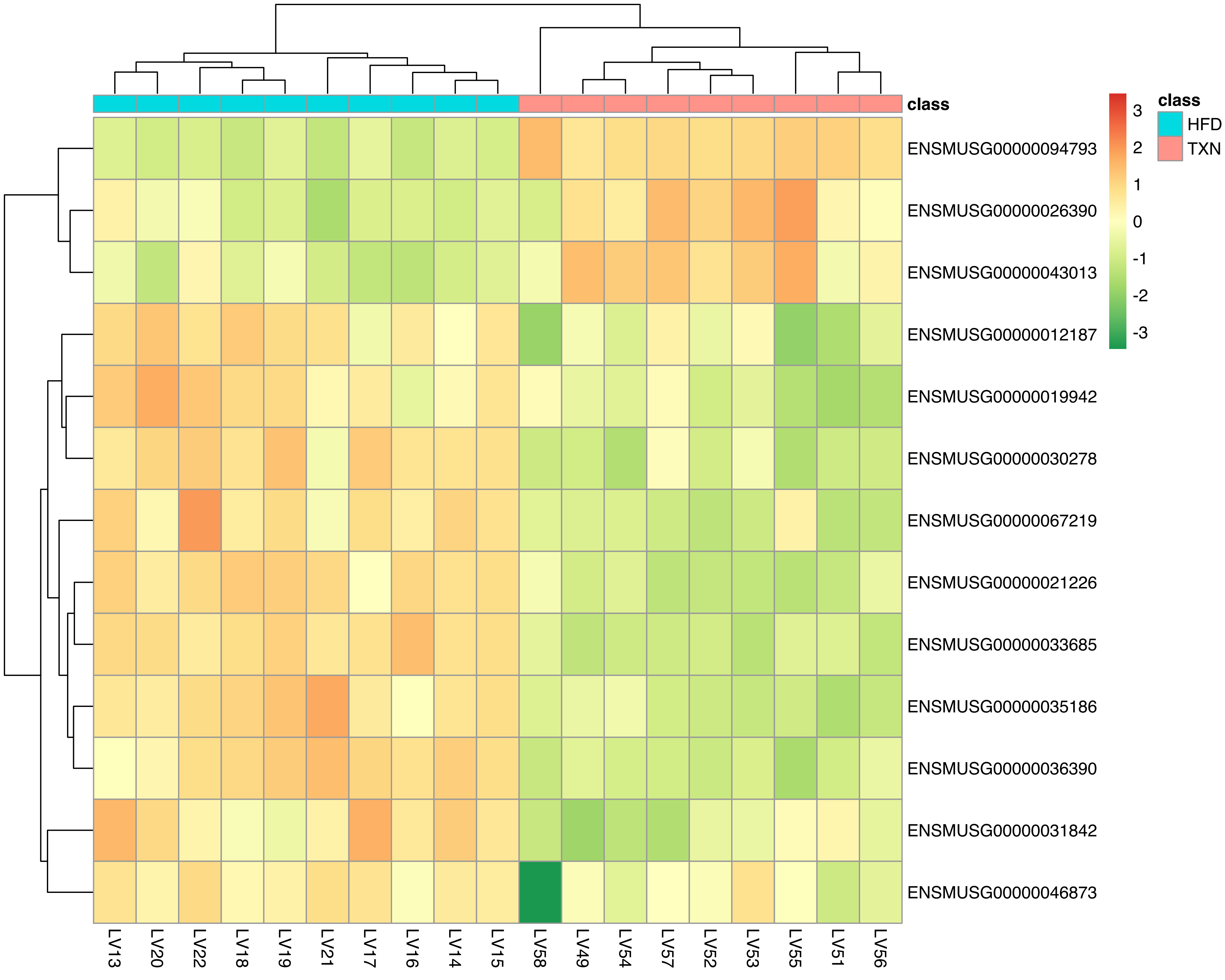

### Rosi2.jpg

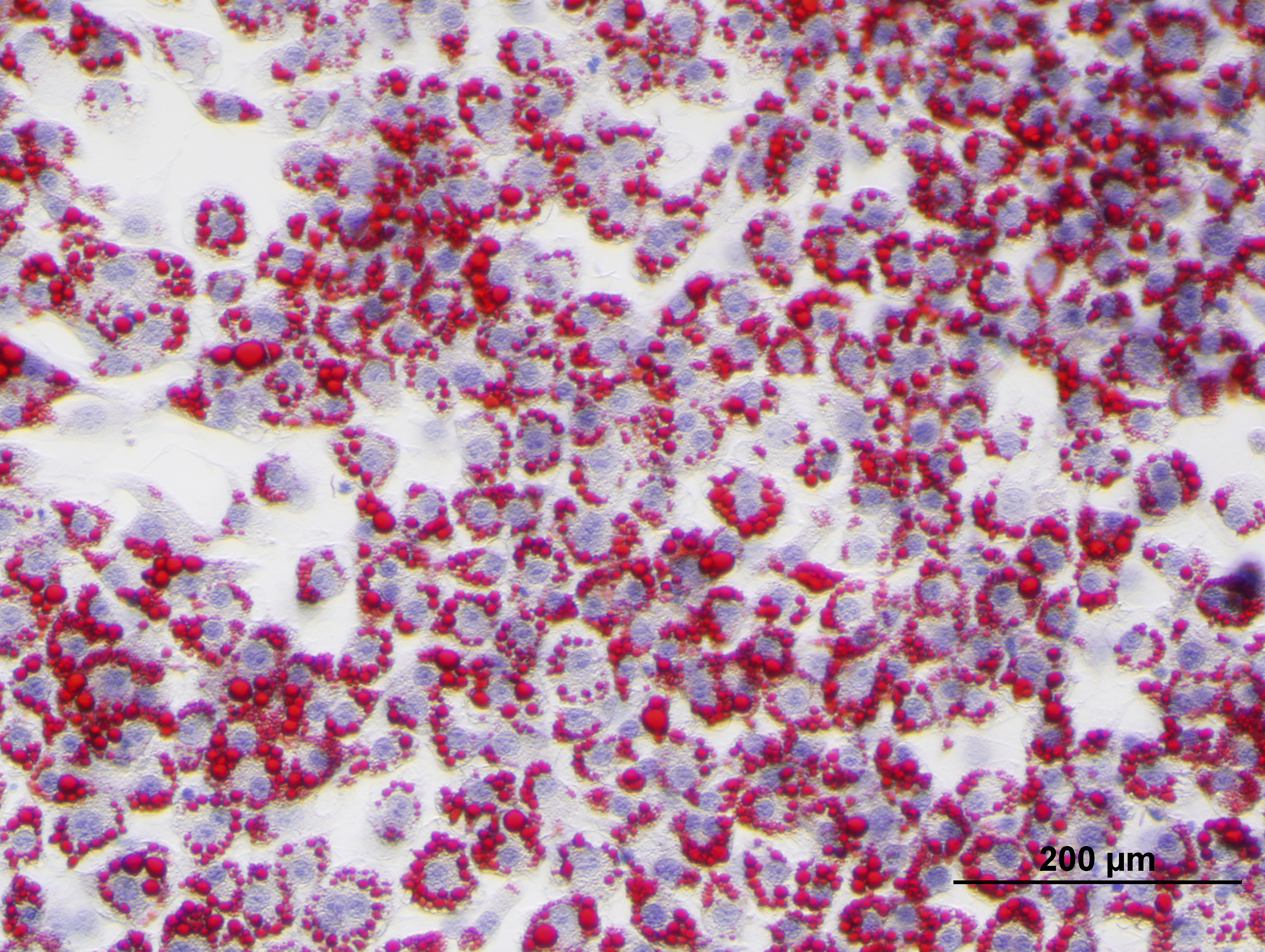

### Rosi3.jpg

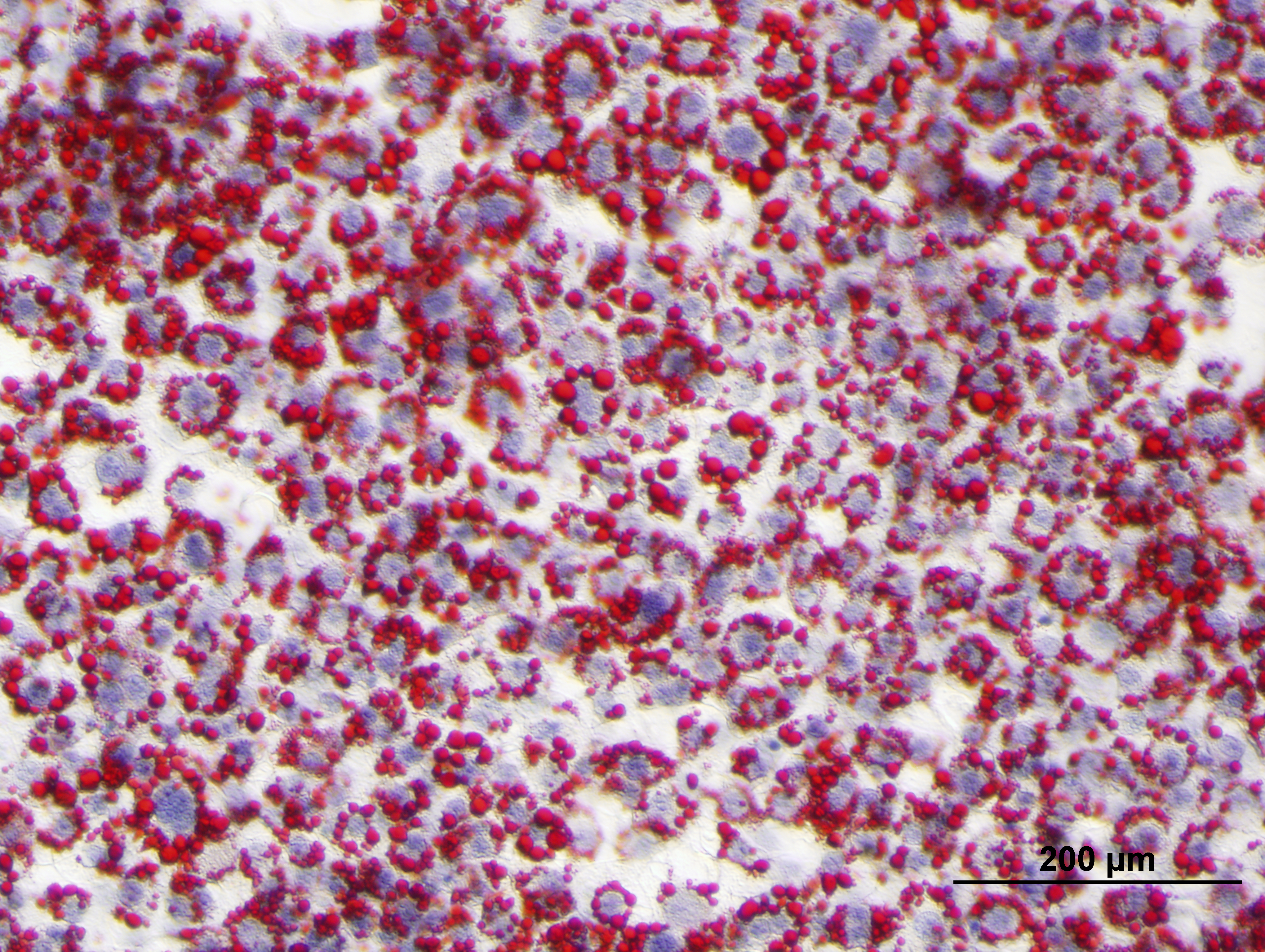

### Rosi+GW2.jpg

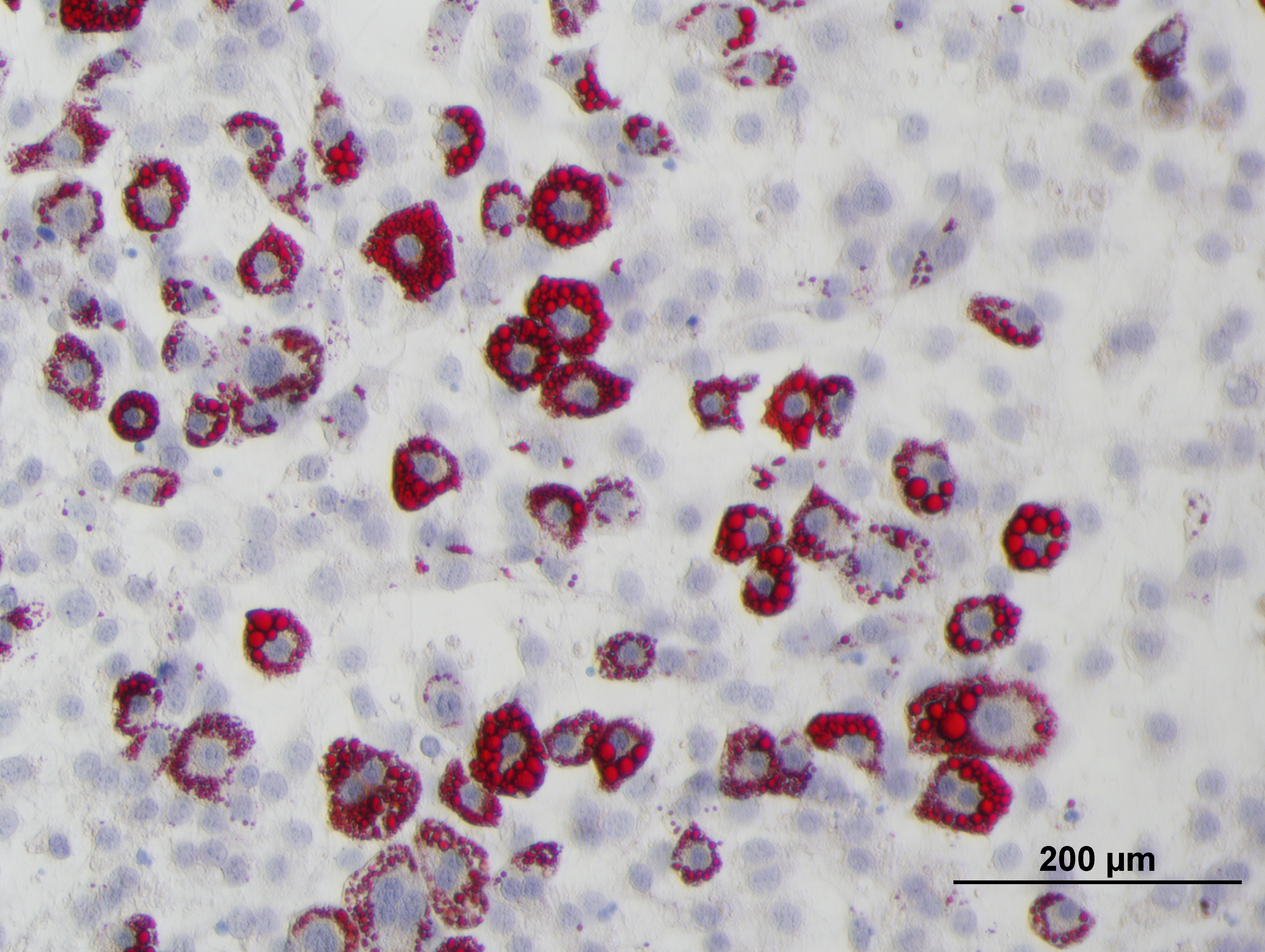

### Rosi+GW3.jpg

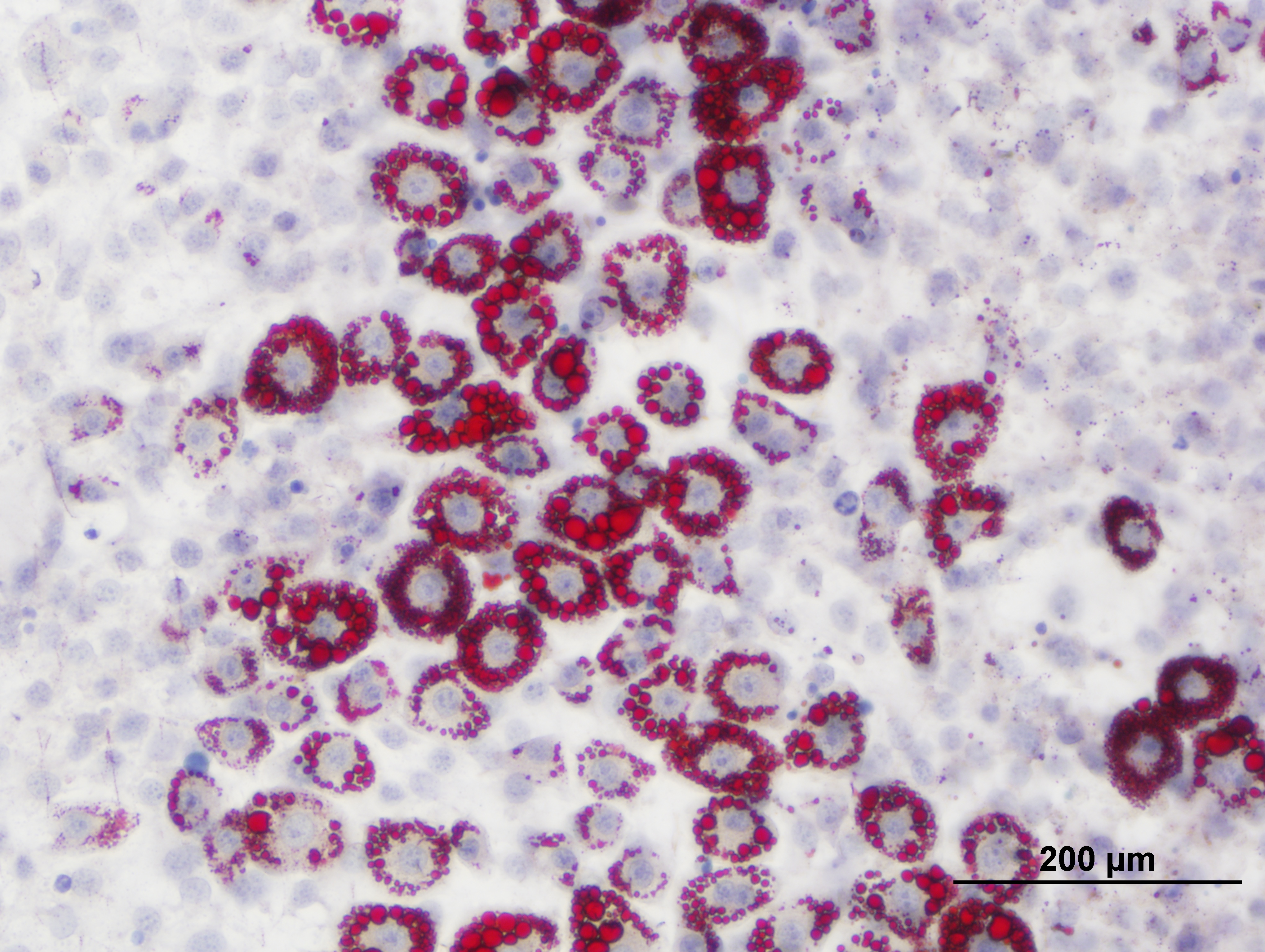

### Rosi+GW.jpg

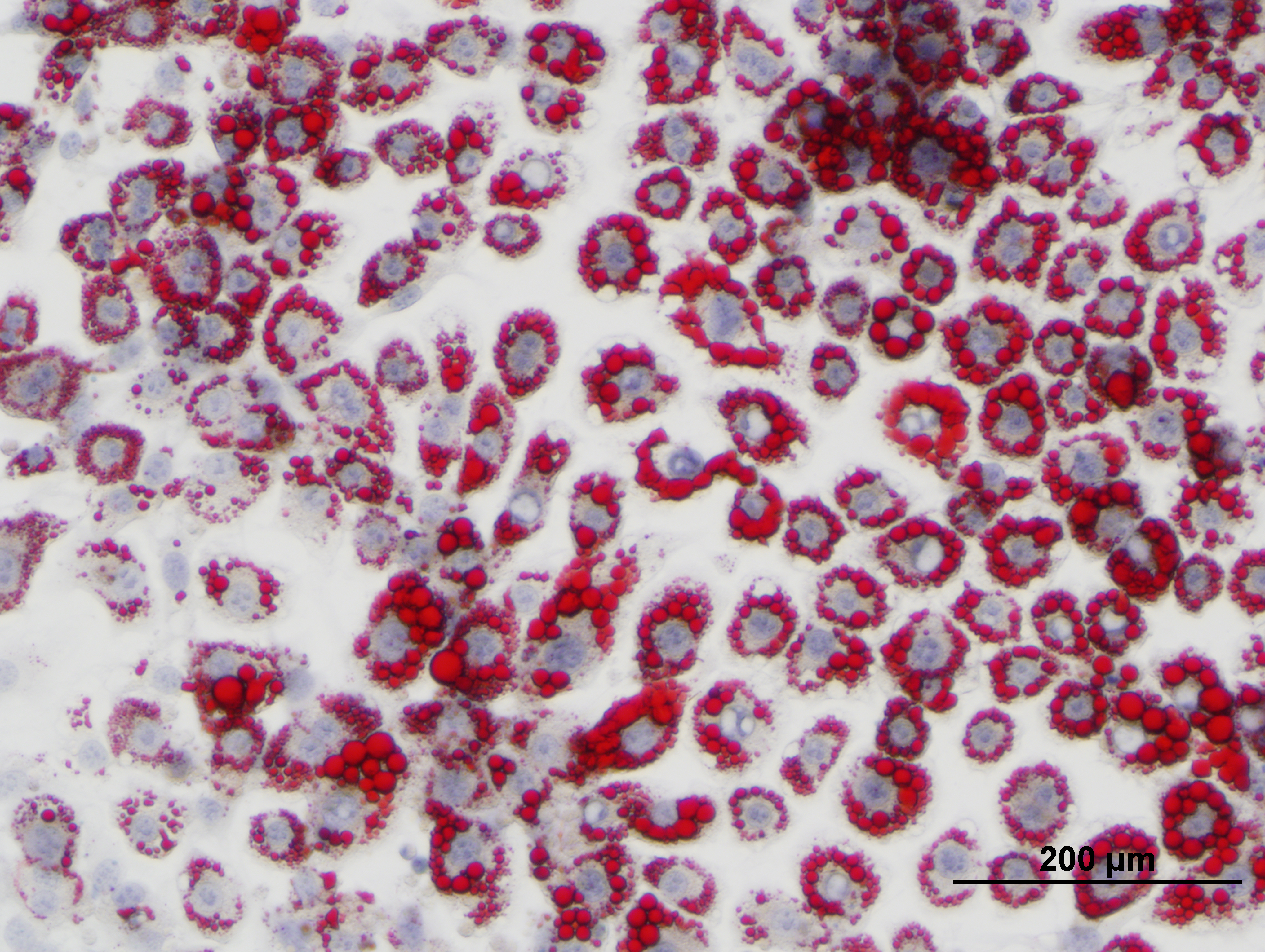

### Rosi.jpg

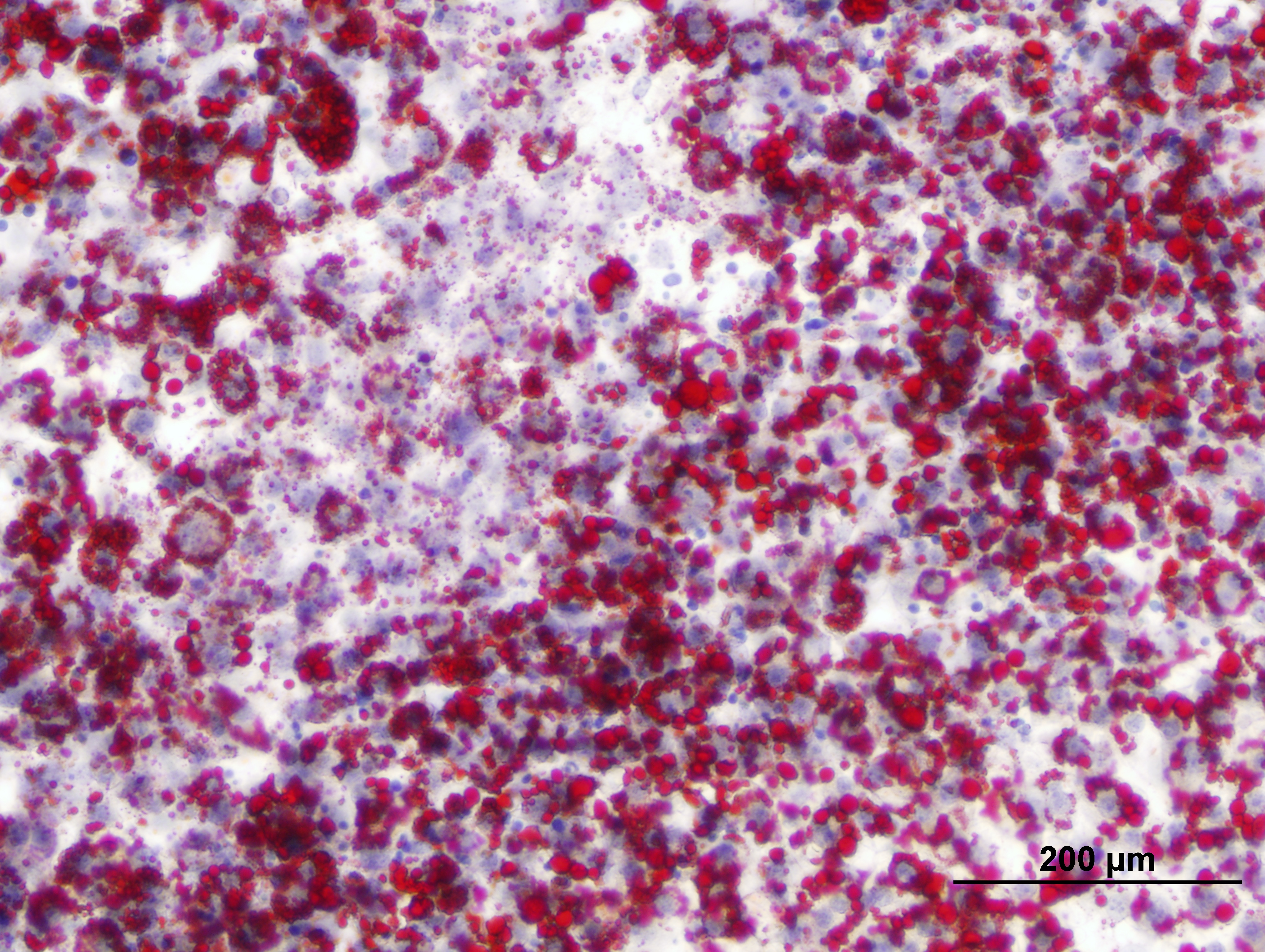

### txnHFDGO.pdf

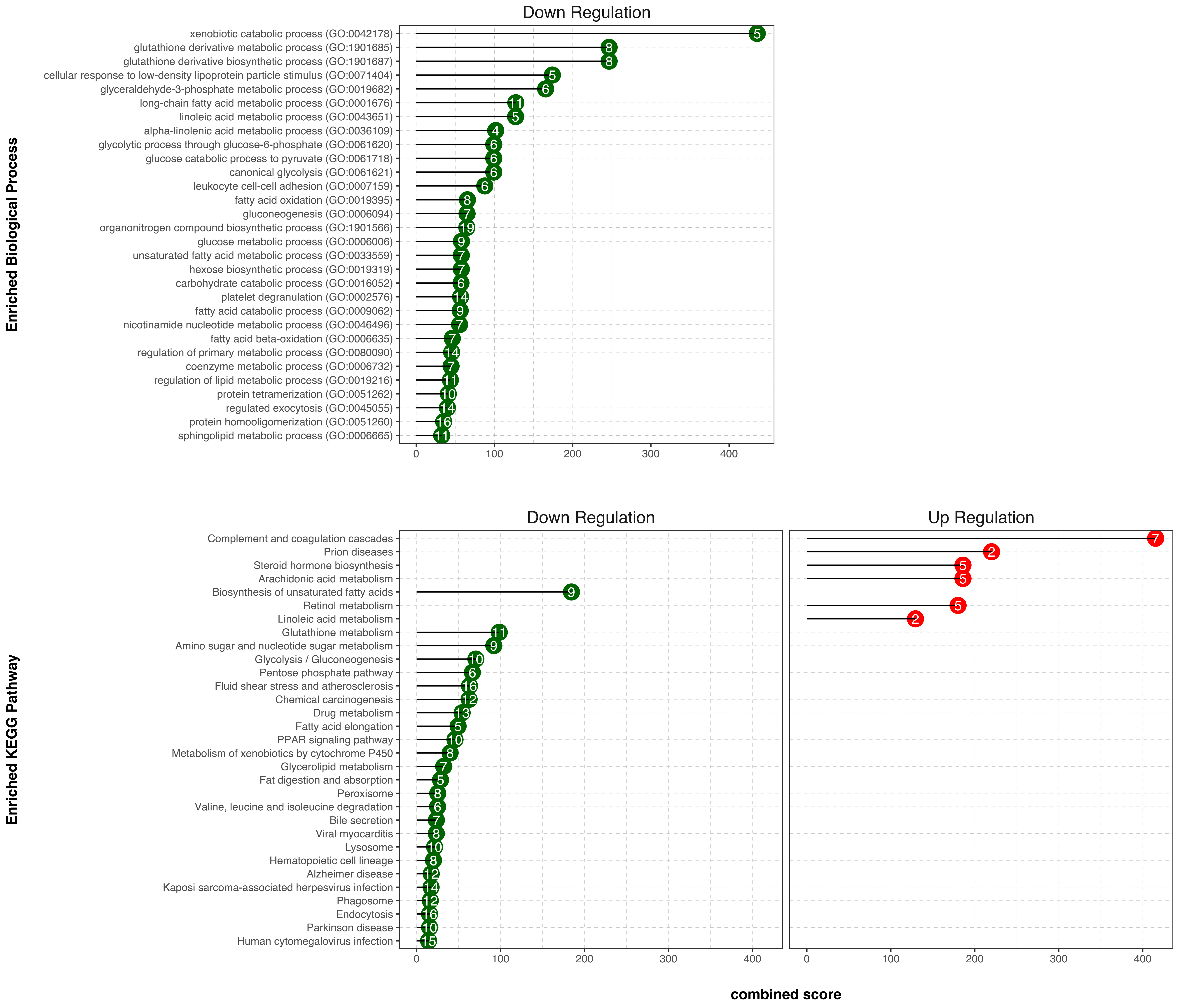

### volplot-hxn-hfd.pdf

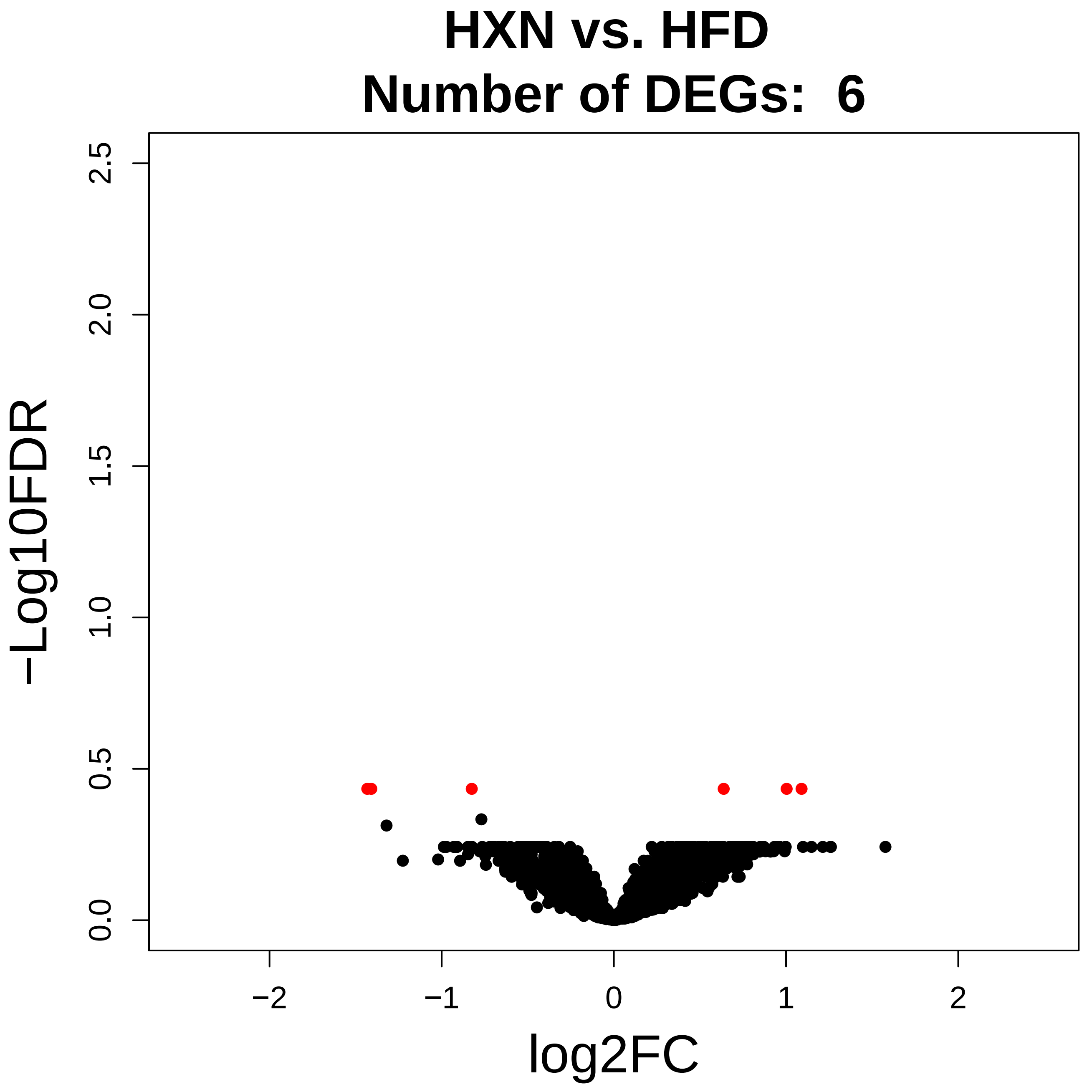

### volplot-lfd-hfd.pdf

##### LFD vs. HFD
##### Number of DEGs: 212

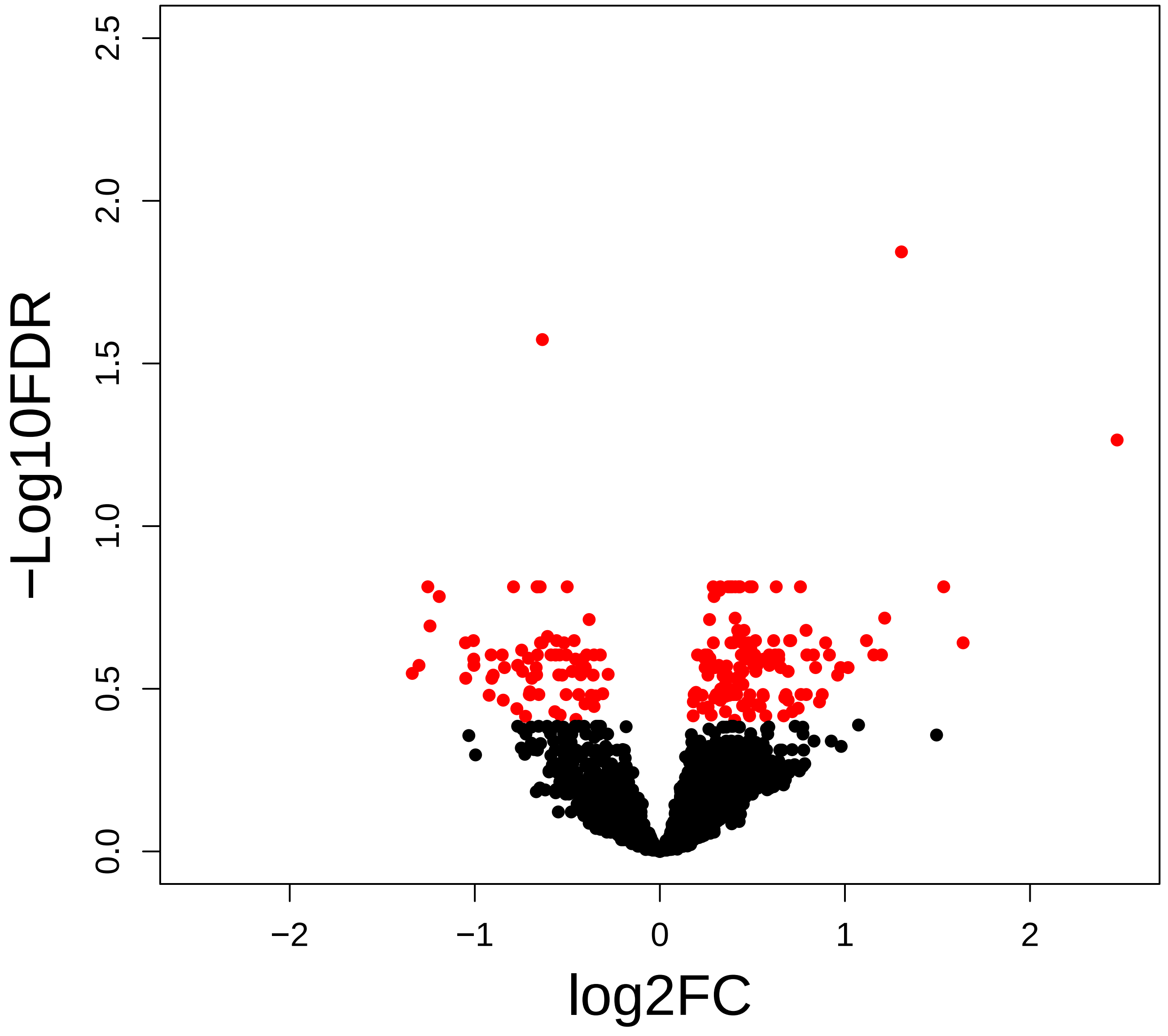

### volplot-txn-hfd.pdf

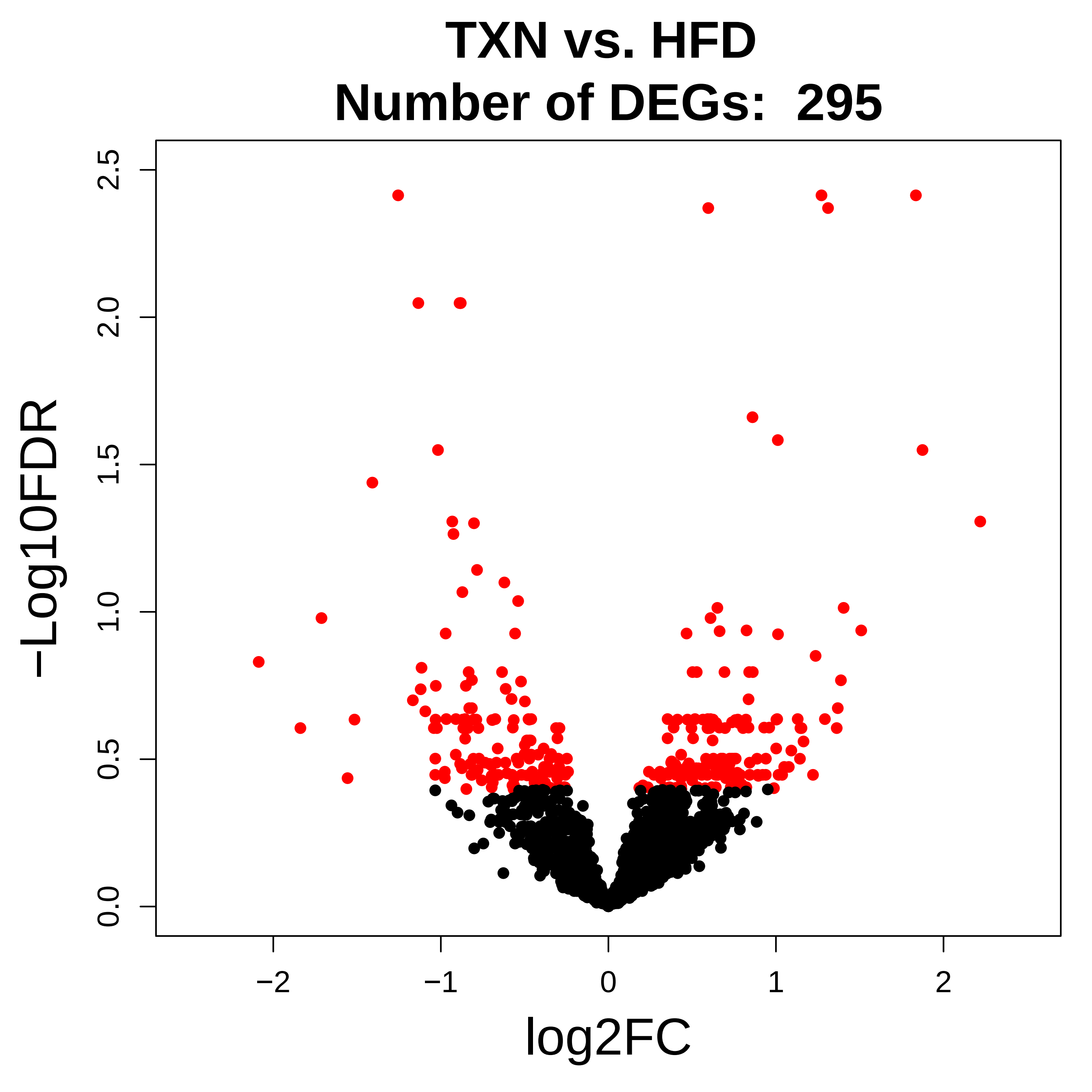
