## Supplementary material for "TXN, a Xanthohumol Derivative, Attenuates High-Fat Diet Induced Hepatic Steatosis by Antagonizing PPARγ": Figure 5 source data: figure5.ipynb

```
# %% [code]
suppressPackageStartupMessages(library(dplyr))
suppressPackageStartupMessages(library(ggplot2))
suppressPackageStartupMessages(library(lsmeans))
suppressPackageStartupMessages(library(patchwork))
suppressPackageStartupMessages(library(export))
R.Version()

# %% [markdown]
# Import data

# %% [markdown]
## - Phenotypes

# %% [code]
data <- read.csv("fig4_table.csv", header = T); 
data$vars <- factor(data$Treatment, levels = c("LFD", "HFD", "HFD+LXN", "HFD+HXN", "HFD+TXN"))
data$vars2 <- factor(data$Treatment, levels = c("HFD", "HFD+LXN", "HFD+HXN", "HFD+TXN", "LFD"))
data <- data %>% mutate(kkcal = Cal_cum/1000); head(data)
txn <- data %>% filter(vars2 == "HFD+TXN"); hfd <- data %>% filter(vars2 == "HFD");
lxn <- data %>% filter(vars2 == "HFD+LXN"); hxn <- data %>% filter(vars2 == "HFD+HXN");
lfd <- data %>% filter(vars2 == "LFD")

# %% [markdown]
# Set theme, import functions

# %% [code]
theme_set(theme_bw())

# %% [code]
theme_legend2 = function(){
	theme(axis.title.y = element_text(size = 15, 
	margin = margin(t=0, r=5, b=0, l=5)),
	axis.text.y = element_text(size = 13),
	axis.text.x = element_text(size = 13),
	axis.title.x = element_text(size = 15,
                               margin = margin(t=10, r=5, b=0, l=0)),
    plot.title = element_text(size = 15, face = "bold"),
	legend.title = element_text(colour = "white"),
	legend.text = element_text(size = 16),
	legend.position = "none",
	plot.margin = unit(c(0.5,0.5,0,0.5), "cm"),
	panel.grid = element_blank())
}

# %% [code]
source("ggplotRegression.R")

# %% [code]
mycolor <- c("#0080ff", "#ffa500", "red", "darkgreen", "black")
mycolor_alpha <- c("#8AB5F9", "#F8CB89", "#F08581", "#87A57E", "#797979")

# %% [code]
Contrasts <- list(
    'LFD vs HFD' = c(-1, 0, 0, 0, 1),
    'LXN vs HFD' = c(-1, 1, 0, 0, 0),
    'HXN vs HFD' = c(-1, 0, 1, 0, 0),
    'TXN vs HFD' = c(-1, 0, 0, 1, 0))

# %% [markdown]
# Figures

# %% [markdown]
## individual fat mass and caloric intake

# %% [markdown]
#### - A. subq

# %% [code]
model <- lm(subq_wt ~ vars2, data = data)
leastsquare <- lsmeans(model, "vars2")
contrast(leastsquare, Contrasts, adjust = "none")

# %% [code]
A <- ggplot(data = data, aes(x = vars2, y = subq_wt, fill = vars2)) +
	stat_boxplot(geom = 'errorbar', linetype = 1, width = 0.2) +
	geom_boxplot(outlier.shape = NA) +
	scale_fill_manual(values = mycolor_alpha) +
	geom_jitter(shape = 20, size = 3, width = 0.2, height = 0.1) +
    labs(x="",y="g",title="A. Subcutaneous fat") +
	theme_legend2()

label.df <- data.frame(vars2 = c("HFD+HXN", "HFD+TXN", "LFD"),
                       subq_wt = c(2.55, 1.75, 1))
A <- A + geom_text(data = label.df, label = c("**", "***", "***"), color='red', size = 8)

# %% [markdown]
#### - B. epididymal

# %% [code]
model <- lm(epi_wat ~ vars2, data = data)
leastsquare <- lsmeans(model, "vars2")
contrast(leastsquare, Contrasts, adjust = "none")

# %% [code]
B <- ggplot(data = data, aes(x = vars2, y = epi_wat, fill = vars2)) +
	stat_boxplot(geom = 'errorbar', linetype = 1, width = 0.2) +
	geom_boxplot(outlier.shape = NA) +
	scale_fill_manual(values = mycolor_alpha) +
	geom_jitter(shape = 20, size = 3, width = 0.2, height = 0.1) +
    labs(x="",y="g",title="B. Epididymal fat") +
	theme_legend2()

label.df <- data.frame(vars2 = c("HFD+HXN", "HFD+TXN", "LFD"),
                       epi_wat = c(3.55, 1.2, 2.22))
B <- B + geom_text(data = label.df, label = c("**", "0.06", "*"), color='red', size = 8)

# %% [markdown]
#### - C. mesenteric

# %% [code]
model <- lm(mesenteric_wt ~ vars2, data = data)
leastsquare <- lsmeans(model, "vars2")
contrast(leastsquare, Contrasts, adjust = "none")

# %% [code]
C <- ggplot(data = data, aes(x = vars2, y = mesenteric_wt, fill = vars2)) +
	stat_boxplot(geom = 'errorbar', linetype = 1, width = 0.2) +
	geom_boxplot(outlier.shape = NA) +
	scale_fill_manual(values = mycolor_alpha) +
	geom_jitter(shape = 20, size = 3, width = 0.2, height = 0.1) +
    labs(x="",y="g",title="C. Mesenteric fat") +
	theme_legend2()

label.df <- data.frame(vars2 = c("HFD+HXN", "HFD+TXN", "LFD"),
                       mesenteric_wt = c(1.83, 1.58, 0.66))
C <- C + geom_text(data = label.df, label = c("*", "***", "***"), color='red', size = 8)

# %% [markdown]
# Regression plots

# %% [code]
lfd_subq_wt_cal <- ggplotRegression(lm(subq_wt ~ kkcal, data = lfd), "black") + 
                        labs(title = "A-1 LFD", x = "1000 kcal", y = "Subcutaneous fat mass (g)")+ theme_legend2()
hfd_subq_wt_cal <- ggplotRegression(lm(subq_wt ~ kkcal, data = hfd), "#0080ff")+ 
                        labs(title = "A-2 HFD", x = "1000 kcal", y = "Subcutaneous fat mass (g)")+ theme_legend2()
lxn_subq_wt_cal <- ggplotRegression(lm(subq_wt ~ kkcal, data = lxn), "#ffa500")+ 
                        labs(title = "A-3 HFD+LXN", x = "1000 kcal", y = "Subcutaneous fat mass (g)")+ theme_legend2()
hxn_subq_wt_cal <- ggplotRegression(lm(subq_wt ~ kkcal, data = hxn), "red")+ ggtitle("HXN")+ 
                        labs(title = "A-4 HFD+HXN", x = "1000 kcal", y = "Subcutaneous fat mass (g)")+ theme_legend2()
txn_subq_wt_cal <- ggplotRegression(lm(subq_wt ~ kkcal, data = txn), "darkgreen")+ 
                        labs(title = "A-5 HFD+TXN", x = "1000 kcal", y = "Subcutaneous fat mass (g)")+ theme_legend2()

# %% [code]
lfd_epi_wat_cal <- ggplotRegression(lm(epi_wat ~ kkcal, data = lfd), "black") + 
                        labs(title = "A-1 LFD", x = "1000 kcal", y = "Epididymal fat mass (g)")+ theme_legend2()
hfd_epi_wat_cal <- ggplotRegression(lm(epi_wat ~ kkcal, data = hfd), "#0080ff")+ 
                        labs(title = "A-2 HFD", x = "1000 kcal", y = "Epididymal fat mass (g)")+ theme_legend2()
lxn_epi_wat_cal <- ggplotRegression(lm(epi_wat ~ kkcal, data = lxn), "#ffa500")+ 
                        labs(title = "A-3 HFD+LXN", x = "1000 kcal", y = "Epididymal fat mass (g)")+ theme_legend2()
hxn_epi_wat_cal <- ggplotRegression(lm(epi_wat ~ kkcal, data = hxn), "red")+ ggtitle("HXN")+ 
                        labs(title = "A-4 HFD+HXN", x = "1000 kcal", y = "Epididymal fat mass (g)")+ theme_legend2()
txn_epi_wat_cal <- ggplotRegression(lm(epi_wat ~ kkcal, data = txn), "darkgreen")+ 
                        labs(title = "A-5 HFD+TXN", x = "1000 kcal", y = "Epididymal fat mass (g)")+ theme_legend2()

# %% [code]
lfd_mesenteric_wt_cal <- ggplotRegression(lm(mesenteric_wt ~ kkcal, data = lfd), "black") + 
                        labs(title = "A-1 LFD", x = "1000 kcal", y = "Mesenteric fat mass (g)")+ theme_legend2()
hfd_mesenteric_wt_cal <- ggplotRegression(lm(mesenteric_wt ~ kkcal, data = hfd), "#0080ff")+ 
                        labs(title = "A-2 HFD", x = "1000 kcal", y = "Mesenteric fat mass (g)")+ theme_legend2()
lxn_mesenteric_wt_cal <- ggplotRegression(lm(mesenteric_wt ~ kkcal, data = lxn), "#ffa500")+ 
                        labs(title = "A-3 HFD+LXN", x = "1000 kcal", y = "Mesenteric fat mass (g)")+ theme_legend2()
hxn_mesenteric_wt_cal <- ggplotRegression(lm(mesenteric_wt ~ kkcal, data = hxn), "red")+ ggtitle("HXN")+ 
                        labs(title = "A-4 HFD+HXN", x = "1000 kcal", y = "Mesenteric fat mass (g)")+ theme_legend2()
txn_mesenteric_wt_cal <- ggplotRegression(lm(mesenteric_wt ~ kkcal, data = txn), "darkgreen")+ 
                        labs(title = "A-5 HFD+TXN", x = "1000 kcal", y = "Mesenteric fat mass (g)")+ theme_legend2()

# %% [markdown]
# Put together

# %% [code]
options(repr.plot.width = 20, repr.plot.height = 4, repr.plot.res = 150)
row1 <- (A | B | C) + plot_layout(nrow = 1, ncol=3)

# %% [code]
options(repr.plot.width = 20, repr.plot.height = 4, repr.plot.res = 150)
row2 <- (lfd_subq_wt_cal | lfd_epi_wat_cal | lfd_mesenteric_wt_cal) + plot_layout(nrow = 1, ncol = 3)

# %% [code]
options(repr.plot.width = 20, repr.plot.height = 4, repr.plot.res = 150)
row3 <- (hfd_subq_wt_cal | hfd_epi_wat_cal | hfd_mesenteric_wt_cal) + plot_layout(nrow = 1, ncol = 3)

# %% [code]
options(repr.plot.width = 20, repr.plot.height = 4, repr.plot.res = 150)
row4 <- (lxn_subq_wt_cal | lxn_epi_wat_cal | lxn_mesenteric_wt_cal) + plot_layout(nrow = 1, ncol = 3)

# %% [code]
options(repr.plot.width = 20, repr.plot.height = 4, repr.plot.res = 150)
row5 <- (hxn_subq_wt_cal | hxn_epi_wat_cal | hxn_mesenteric_wt_cal) + plot_layout(nrow = 1, ncol = 3)

# %% [code]
options(repr.plot.width = 20, repr.plot.height = 4, repr.plot.res = 150)
row6 <- (txn_subq_wt_cal | txn_epi_wat_cal | txn_mesenteric_wt_cal) + plot_layout(nrow = 1, ncol = 3)

# %% [code]
options(repr.plot.width = 20, repr.plot.height = 25, repr.plot.res = 200)
figure4 <- wrap_plots(row1, row2, row3, row4, row5, row6, nrow=6)

# %% [code]
#graph2pdf(figure4, "figure4.pdf", width = 20, height = 25)

# %% [code]
```
