## Supplementary material for "TXN, a Xanthohumol Derivative, Attenuates High-Fat Diet Induced Hepatic Steatosis by Antagonizing PPARγ": Figure 6 source data: fig6.ipynb

```
# %% [code]
## Load packages

# %% [code]
suppressPackageStartupMessages(library(dplyr))
suppressPackageStartupMessages(library(ggplot2))
suppressPackageStartupMessages(library(lsmeans))
suppressPackageStartupMessages(library(patchwork))
suppressPackageStartupMessages(library(export))
R.Version()

# %% [markdown]
# Import data

# %% [markdown]
## - Promethion

# %% [markdown]
## - Phenotypes

# %% [code]
data <- read.csv("fig6_table.csv", header = T); 
data$vars <- factor(data$Treatment, levels = c("LFD", "HFD", "HFD+LXN", "HFD+HXN", "HFD+TXN"))
data$vars2 <- factor(data$Treatment, levels = c("HFD", "HFD+LXN", "HFD+HXN", "HFD+TXN", "LFD"))
data <- data %>% mutate(lv2wt = liver_wt*100 / wt); head(data)

# %% [markdown]
# Set theme, import functions

# %% [code]
theme_set(theme_bw())

# %% [code]
theme_legend2 = function(){
	theme(axis.title.y = element_text(size = 15, 
	margin = margin(t=0, r=5, b=0, l=5)),
	axis.text.y = element_text(size = 13),
	axis.text.x = element_text(size = 13),
	axis.title.x = element_text(size = 15,
                               margin = margin(t=10, r=5, b=0, l=0)),
    plot.title = element_text(size = 15, face = "bold"),
	legend.title = element_text(colour = "white"),
	legend.text = element_text(size = 16),
	legend.position = "none",
	plot.margin = unit(c(0.5,0.5,0,0.5), "cm"),
	panel.grid = element_blank())
}

# %% [code]
mycolor <- c("#0080ff", "#ffa500", "red", "darkgreen", "black")
mycolor_alpha <- c("#8AB5F9", "#F8CB89", "#F08581", "#87A57E", "#797979")

# %% [code]
Contrasts <- list(
    'LFD vs HFD' = c(-1, 0, 0, 0, 1),
    'LXN vs HFD' = c(-1, 1, 0, 0, 0),
    'HXN vs HFD' = c(-1, 0, 1, 0, 0),
    'TXN vs HFD' = c(-1, 0, 0, 1, 0))

# %% [markdown]
# Figures

# %% [markdown]
## - A. fat mass boxplot

# %% [code]
model <- lm(lv2wt ~ vars2, data = data)
leastsquare <- lsmeans(model, "vars2")
contrast(leastsquare, Contrasts, adjust = "none")

# %% [code]
A <- ggplot(data = data, aes(x = vars2, y = lv2wt, fill = vars2)) +
	stat_boxplot(geom = 'errorbar', linetype = 1, width = 0.2) +
	geom_boxplot(outlier.shape = NA) +
	scale_fill_manual(values = mycolor_alpha) +
	geom_jitter(shape = 20, size = 3, width = 0.2, height = 0.1) +
    labs(x="",y="%",title="Liver mass relative to body weight") +
    geom_hline(yintercept=4, linetype="dashed", color = "black") +
	theme_legend2()

label.df <- data.frame(vars2 = c("HFD+HXN", "HFD+TXN", "LFD"),
                       lv2wt = c(5.5, 3.8, 4.65))
A <- A + geom_text(data = label.df, label = c("**", "***", "***"), color='red', size = 8)

# %% [markdown]
## - B. Liver TAG

# %% [code]
model <- lm(LV_TAG ~ vars2, data = data)
leastsquare <- lsmeans(model, "vars2")
contrast(leastsquare, Contrasts, adjust = "none")

# %% [code]
B <- ggplot(data = data, aes(x = vars2, y = LV_TAG, fill = vars2)) +
	stat_boxplot(geom = 'errorbar', linetype = 1, width = 0.2) +
	geom_boxplot(outlier.shape = NA) +
	scale_fill_manual(values = mycolor_alpha) +
	geom_jitter(shape = 20, size = 3, width = 0.2, height = 0.1) +
    labs(x="",y="mg/g",title="Liver TAG") +
	theme_legend2()
label.df <- data.frame(vars2 = c("HFD+TXN", "LFD"),
                       LV_TAG = c(630, 520))
B <- B + geom_text(data = label.df, label = c("***", "***"), color='red', size = 8)

# %% [code]
#options(repr.plot.width = 7, repr.plot.height = 6, repr.plot.res = 150)
graph2pdf(A, "B.pdf", width = 7, height = 6)
graph2pdf(B, "C.pdf", width = 7, height = 6)

# %% [code]
#
```
