## Supplementary material for "TXN, a Xanthohumol Derivative, Attenuates High-Fat Diet Induced Hepatic Steatosis by Antagonizing PPARγ": Figure 7 source data: bams2RDA.R

```
################################################################################
### Liver
### Input: BAM files produced by STAR, sample metadata file, GTF file
### Output: DGEList, count table 
################################################################################
library(knitr)
.cran_packages <- c("Rsamtools", "GenomicFeatures", "GenomicAlignments", "edgeR", "Mus.musculus")
.inst <- .cran_packages %in% installed.packages()
if(any(!.inst)) {
    install.packages(.cran_packages[!.inst])
}
sapply(.cran_packages, require, character.only = TRUE)
################################################################################
set.seed(20191103)

sampleTable <- read.csv("transcriptome_meta_LV.csv", row.names=1)
filenames <- file.path("BAMs", paste0(rownames(sampleTable), ".bam"))
bamfiles <- BamFileList(filenames, yieldSize=2000000)
gtffile <- file.path("gencode.vM22.primary_assembly.annotation.gtf")
txdb <- makeTxDbFromGFF(gtffile, format="gtf")
ebg <- exonsBy(txdb, by="gene")
se <- summarizeOverlaps(features=ebg, 
                        reads=bamfiles,
                        mode="Union",
                        singleEnd=TRUE)


colData(se) <- DataFrame(sampleTable)
row.names(se) = gsub("\\..*", "", row.names(se))
genetable <- data.frame(gene.id=rownames(se))
countdata <- assay(se)
coldata <- colData(se)

seGroups <- c(
    "LFD", "LFD", "LFD", "LFD", "LFD", "LFD", "LFD", "LFD", "LFD", "LFD", 
    "HFD", "HFD", "HFD", "HFD", "HFD", "HFD", "HFD", "HFD", "HFD", "HFD",  
    "HXN", "HXN", "HXN", "HXN", "HXN", "HXN", "HXN", "HXN", "HXN", "HXN", 
    "TXN", "TXN", "TXN", "TXN", "TXN", "TXN", "TXN", "TXN", "TXN")

y <- DGEList(counts=countdata, 
             samples=coldata, 
             group=factor(seGroups),
             genes=genetable)

y$genes$entrez <- mapIds(Mus.musculus,
                     keys=row.names(y$genes),
                     column="ENTREZID",
                     keytype="ENSEMBL",
                     multiVals="first")
y$genes$symbol <- mapIds(Mus.musculus,
                     keys=row.names(y$genes),
                     column="SYMBOL",
                     keytype="ENSEMBL",
                     multiVals="first")
y$genes$gene.name <- mapIds(Mus.musculus,
                     keys=row.names(y$genes),
                     column="GENENAME",
                     keytype="ENSEMBL",
                     multiVals="first")

toRemove <- c("ribosomal protein", "RIKEN", "predicted gene", "pseudogene",
			  "cDNA sequence", "nuclear encoded rRNA", "microRNA", 
			  "DNA segment", "uncharacterized", "pre-RNA", "20S rRNA",
			  "non-protein coding")

yy <- y$genes[!is.na(y$genes$gene.name), ]
yy <- yy[!grepl(paste(toRemove, collapse = "|"), yy$gene.name), ]
y_subset <- y[rownames(yy), ]

y_filtered <- y[ !(y$genes$gene.name %in% y_subset$genes$gene.name), ]


keep <- filterByExpr(y_subset, group=y$samples$Diet, 
    min.count=11, min.total.count=15)
y_keep <- y_subset[keep, , keep.lib.sizes=FALSE]

y_keep_norm <- calcNormFactors(y_keep, method = "TMM")

design <- model.matrix(~ 0 + y_keep_norm$samples$Diet)
colnames(design) <- levels(y_keep_norm$samples$Diet)
## estimate dispersions
y_keep_norm_disp <- estimateDisp(y_keep_norm, design, robust=TRUE)

filtered_counts <- data.frame(y_keep$counts)
filtered_counts$entrez <- mapIds(Mus.musculus,
                     keys=row.names(filtered_counts),
                     column="ENTREZID",
                     keytype="ENSEMBL",
                     multiVals="first")
filtered_counts$symbol <- mapIds(Mus.musculus,
                     keys=row.names(filtered_counts),
                     column="SYMBOL",
                     keytype="ENSEMBL",
                     multiVals="first")
filtered_counts$gene.name <- mapIds(Mus.musculus,
                     keys=row.names(filtered_counts),
                     column="GENENAME",
                     keytype="ENSEMBL",
                     multiVals="first")

save(y, y_subset, y_filtered, y_keep, filtered_counts, y_keep_norm, y_keep_norm_disp, file = "transcriptome_LV.rda")
write.csv(filtered_counts, "countTable_keptLV.csv")


sink("stdout.LV")

cat("\n===================================================================================\n")
cat("Original raw DGEList has:", dim(y)[1], "genes", "after removing NAs (unclassified genes) there are", dim(y_subset)[1], "genes left.\n")
print(toRemove)
cat("\nGenes that have thoses names were further removed before filtering out lowly expressed genes. A total of", dim(y_filtered)[1], "genes were removed.\n") 
cat("Finally, there are", dim(y_keep)[1], "liver genes retained for downstream analyses.")
cat("\n===================================================================================\n")
sessionInfo()

sink()
```
