## Supplementary material for "TXN, a Xanthohumol Derivative, Attenuates High-Fat Diet Induced Hepatic Steatosis by Antagonizing PPARγ": Figure 7 source data: fig7.ipynb

```
# %% [code]
suppressPackageStartupMessages(library(Rsamtools))
suppressPackageStartupMessages(library(GenomicFeatures))
suppressPackageStartupMessages(library(mixOmics))
suppressPackageStartupMessages(library(gplots))
suppressPackageStartupMessages(library(ggplot2))
suppressPackageStartupMessages(library(Mus.musculus))
suppressPackageStartupMessages(library(edgeR))
suppressPackageStartupMessages(library(GenomicAlignments))
R.Version()

# %% [code]
y.keep <- readRDS('y_keep.rds')

# %% [code]
design <- model.matrix(~ 0 + y.keep$samples$Diet)
colnames(design) <- levels(y.keep$samples$Diet)

# %% [code]
y.keep.disp <- estimateDisp(y.keep, design, robust=TRUE)

# %% [code]
logCPM <- cpm(y.keep.disp, prior.count=2, log=TRUE)
colnames(logCPM) <- rownames(y.keep.disp$samples)

# %% [code]
fit <- glmQLFit(y.keep.disp, design, robust=TRUE)
con <- makeContrasts(TXN - HFD, levels=design)
qlf <- glmQLFTest(fit, contrast=con)
o <- order(qlf$table$PValue)
length(o)

# %% [code]
all <- logCPM[o[1:200],]
all <- t(scale(t(all)))

# %% [code]
col.grp <- c("grey", "grey", "grey", "grey", "grey", "grey", "grey", "grey", "grey", "grey", 
             "#0080ff", "#0080ff", "#0080ff", "#0080ff", "#0080ff", "#0080ff", "#0080ff", "#0080ff", "#0080ff", "#0080ff", 
             "red",  "red", "red", "red", "red", "red", "red", "red", "red", "red", 
             "darkgreen",  "darkgreen", "darkgreen", "darkgreen", "darkgreen", "darkgreen", "darkgreen", "darkgreen", "darkgreen")
cond.col <- c("LFD" = "grey", "HFD" = "#0080ff", "HXN" = "red", "TXN" = "darkgreen")

# %% [code]
obj.cim=cim(all, cluster = 'both', col.sideColors=col.grp, 
            dist.method = c("euclidean", "euclidean"),
            clust.method = c("complete", "complete"),
            legend=list(legend = unique(y.keep.disp$samples$group), 
                        col = cond.col, title = "Diet", cex = 0.7)
           )

# %% [code]
pdf("heatmap.pdf", width = 12, height = 14)
obj.cim=cim(all, cluster = 'both', keysize=c(0.8,0.4),
            col.sideColors=col.grp, col.names=FALSE, row.cex=0.4,
            dist.method = c("euclidean", "euclidean"),
            clust.method = c("complete", "complete"),
            legend=list(legend = unique(y.keep.disp$samples$group), 
                       col = cond.col, title = "Diet", cex = 0.9)
           )
dev.off()

# %% [code]
#pdf("heatmap_groupTreatments.pdf")
#fig <- cim(all, col.names=TRUE, 
#    cluster = "row", symkey=FALSE, keysize=c(1.1,0.8),
#    row.cex=0.3, col.sideColors=col.grp, dist.method = c("euclidean", "euclidean"),
#    clust.method = c("complete", "complete"),
#    legend=list( legend = unique(y.keep.disp$samples$group), col = cond.col, title = "Diet", cex = 0.7))
#dev.off()

# %% [code]
con2 <- makeContrasts(HXN - HFD, levels=design)
con3 <- makeContrasts(LFD - HFD, levels=design)

stat_txnHFD_qlf <- topTags(qlf, n=nrow(y.keep.disp))$table
qlf2 <- glmQLFTest(fit, contrast=con2)
stat_hxnHFD_qlf <- topTags(qlf2, n=nrow(y.keep.disp))$table
qlf3 <- glmQLFTest(fit, contrast=con3)
stat_lfdHFD_qlf <- topTags(qlf3, n=nrow(y.keep.disp))$table

# %% [code]
pdf("volplot-txn-hfd.pdf")
par(mar = c(5, 5, 4.5, 1))
volcanoData <- cbind(stat_txnHFD_qlf$logFC, -log10(stat_txnHFD_qlf$FDR))
colnames(volcanoData) <- c("log2FC", "-Log10FDR")
DEGs <- stat_txnHFD_qlf$FDR < 0.4
point.col <- ifelse(DEGs, "red", "black")
sign.genes=which(stat_txnHFD_qlf$FDR<0.4)
plot(volcanoData, pch=16, col = point.col, cex = 1, 
     xlim = c(-2.5,2.5), ylim = c(0, 2.5),
     main=paste("TXN vs. HFD", "\nNumber of DEGs: ", table(DEGs)[2]),
     cex.main=2, cex.lab=2, cex.axis=1.2)
     text(x=stat_txnHFD_qlf$logFC[sign.genes] , y=-log10(stat_txnHFD_qlf$FDR[sign.genes]), 
     label=NULL)

dev.off()

# %% [code]
pdf("volplot-hxn-hfd.pdf")
par(mar = c(5, 5, 4.5, 1))

volcanoData <- cbind(stat_hxnHFD_qlf$logFC, -log10(stat_hxnHFD_qlf$FDR))
colnames(volcanoData) <- c("log2FC", "-Log10FDR")
DEGs <- stat_hxnHFD_qlf$FDR < 0.4
point.col <- ifelse(DEGs, "red", "black")
sign.genes=which(stat_hxnHFD_qlf$FDR<0.4)

plot(volcanoData, pch=16, col = point.col, cex = 1, xlim = c(-2.5,2.5), ylim = c(0, 2.5),
     main=paste("HXN vs. HFD", "\nNumber of DEGs: ", table(DEGs)[2]),
     cex.main=2, cex.lab=2, cex.axis=1.2)

text(x=stat_hxnHFD_qlf$logFC[sign.genes] , y=-log10(stat_hxnHFD_qlf$FDR[sign.genes]), 
    label=NULL)
dev.off()

# %% [code]
pdf("volplot-lfd-hfd.pdf")
par(mar = c(5, 5, 4.5, 1))
volcanoData <- cbind(stat_lfdHFD_qlf$logFC, -log10(stat_lfdHFD_qlf$FDR))
colnames(volcanoData) <- c("log2FC", "-Log10FDR")
DEGs <- stat_lfdHFD_qlf$FDR < 0.4
#table(DEGs)
point.col <- ifelse(DEGs, "red", "black")
sign.genes=which(stat_lfdHFD_qlf$FDR<0.4)

plot(volcanoData, pch=16, col = point.col, cex = 1, main=paste("LFD vs. HFD", "\nNumber of DEGs: ", table(DEGs)[2]),
     xlim = c(-2.5,2.5), ylim = c(0, 2.5), cex.main=2, cex.lab=2, cex.axis=1.2)
     text(x=stat_lfdHFD_qlf$logFC[sign.genes] , y=-log10(stat_lfdHFD_qlf$FDR[sign.genes]), 
     label=NULL)
dev.off()

# %% [code]
#library(imager)
#file <- load.image('volplot-txn-hfd.jpeg')
#fig <- plot(file, axes=FALSE)
```
