## Supplementary material for "TXN, a Xanthohumol Derivative, Attenuates High-Fat Diet Induced Hepatic Steatosis by Antagonizing PPARγ": Figure 8 source data: fig8.ipynb

```
# %% [code]
suppressPackageStartupMessages(library(export))
suppressPackageStartupMessages(library(edgeR))
suppressPackageStartupMessages(library(ggplot2))
suppressPackageStartupMessages(library(dplyr))
suppressPackageStartupMessages(library(patchwork))
theme_set(theme_bw())

# %% [code]
y_keep <- readRDS('y_keep.rds')
theme_legend_free = function(){
	theme(axis.title.y = element_text(size = 17, 
	margin = margin(t=0, r=5, b=0, l=10)),
	axis.text.y = element_text(size = 17, face = 'bold'),
	axis.text.x = element_text(size = 17, face= 'bold'),
	axis.ticks.x = element_blank(),
	legend.position = "none",
    plot.title = element_text(size = 24, face = 'bold'),
	plot.margin = unit(c(0.5,0.5,0,0.5), "cm"),
	panel.grid = element_blank(),
	panel.grid.major = element_blank(),
	panel.grid.minor = element_blank())
}

# %% [code]
design <- model.matrix(~ 0 + y_keep$samples$Diet)
colnames(design) <- levels(y_keep$samples$Diet)

# %% [code]
y.keep.disp <- estimateDisp(y_keep, design, robust=TRUE)

# %% [code]
fit <- glmQLFit(y.keep.disp, design, robust=TRUE)
con <- makeContrasts(TXN - HFD, levels=design)
qlf <- glmQLFTest(fit, contrast=con)
stat_txnHFD_qlf <- topTags(qlf, n=nrow(y.keep.disp))$table

# %% [code]
kegg <- read.csv('./processed/KEGGterms.csv', header=T)
orderedk <- kegg %>%
select(Direction, Combined.Score, pathwayTerms, enrichedGenes) %>%
arrange(desc(Combined.Score))

# %% [code]
plot <- ggplot(orderedk, aes(x=reorder(`pathwayTerms`, Combined.Score), y=Combined.Score, label=enrichedGenes)) + 
  geom_point(aes(color=Direction), stat='identity', size=6)  +
  scale_color_manual(values = c('darkgreen','red')) +
  geom_segment(aes(y = 0, 
                   x = `pathwayTerms`, 
                   yend = Combined.Score, 
                   xend = `pathwayTerms`)) +
  geom_text(color="white", size=4) +
  labs(#title="", 
       #subtitle="TXN vs. HFD",
       x = 'Enriched KEGG Pathway', y = 'combined score') + 
  coord_flip()+
  facet_wrap(~Direction, ncol=2) +
  theme(#axis.text.x = element_text(size = 10, face='bold'),
           #axis.text.y = element_text(size = 14),
           axis.title.y = element_text(margin = margin(t = 0, r = 20, b = 0, l = 0)),
           axis.title.x = element_text(margin = margin(t = 20, r = 0, b = 0, l = 0), size=12),
        axis.title = element_text(size = 12, face = "bold"),
        plot.title = element_text(size = 14, face = "bold"),
        strip.background = element_blank(),
        strip.text = element_text(size = 14),
        plot.margin = margin(0, 0.5, 0.5, 0.5, "cm"),
        panel.grid.major = element_line(color='grey', linetype='dashed', size=0.1),
        panel.grid.minor = element_line(color='grey', linetype='dashed', size=0.1),
        legend.position = "none")

# %% [code]
bp <- read.csv('./processed/BPTerms.csv', header=T)
ordered <- bp %>%
select(Direction, Combined.Score, bpTerms, enrichedGenes) %>%
arrange(desc(Combined.Score))

# %% [code]
bpplot <- ggplot(ordered, aes(x=reorder(`bpTerms`, Combined.Score), y=Combined.Score, label=enrichedGenes)) + 
  geom_point(stat='identity', color="darkgreen", size=6)  +
  geom_segment(aes(y = 0, 
                   x = `bpTerms`, 
                   yend = Combined.Score, 
                   xend = `bpTerms`)) +
  geom_text(color="white", size=4) +
  labs(#title="", 
       #subtitle="TXN vs. HFD",
       x = 'Enriched Biological Process', y = '') + 
  coord_flip()+
  facet_wrap(~Direction, ncol=2) +
  theme(axis.title.y = element_text(margin = margin(t = 0, r = 20, b = 0, l = 0)),
        axis.title.x = element_text(margin = margin(t = 20, r = 0, b = 0, l = 0), size=12),
        axis.title = element_text(size = 12, face = "bold"),
        plot.title = element_text(size = 14, face = "bold"),
        strip.background = element_blank(),
        strip.text = element_text(size = 14),
        plot.margin = margin(0, 0.5, 0, 0.5, "cm"),
        panel.grid.major = element_line(color='grey', linetype='dashed', size=0.1),
        panel.grid.minor = element_line(color='grey', linetype='dashed', size=0.1),
        legend.position = "none")

# %% [code]
GO <- (bpplot + plot_spacer())/plot

# %% [code]
GO

# %% [code]
#graph2pdf(GO, file = "txnHFDGO.pdf", width = 14, height = 12)
```
