## Supplementary material for "TXN, a Xanthohumol Derivative, Attenuates High-Fat Diet Induced Hepatic Steatosis by Antagonizing PPARγ": Figure 9 source data: SVM_hftxn.ipynb

```
# %% [code]
suppressPackageStartupMessages(library(DaMiRseq))
suppressPackageStartupMessages(library(export))

# %% [code]
count_data <- read.csv("countMatrix_final_hftxn.csv", row.names=1)
head(count_data)

# %% [code]
covariate_data <- read.csv("colData_hftxn.csv", row.names=1)
covariate_data$class = as.factor(covariate_data$class)
mytable <- DaMiR.makeSE(count_data, covariate_data)

# %% [code]
#colData(mytable)

# %% [markdown]
# Preprocessing and Normalization

# %% [markdown]
### normalization

# %% [code]
data_norm <- DaMiR.normalization(
mytable, minCounts=9, fSample=0.8, th.cv=3, type='logcpm')
#outputfile <- "DataNormalized_hfdtxn.txt"
#write.table(assays(data_norm)$'', file = outputfile, quote = FALSE, sep = "\t")

# %% [markdown]
### sample filtering

# %% [code]
data_filt <- DaMiR.sampleFilt(data_norm, th.corr=0.9)
#outputfile <- "dataFilt_hftxn.txt"
#write.table(assay(data_filt)[, 1:19], file = outputfile, quote=FALSE, sep = "\t", header=True)

# %% [markdown]
# Adjusting Data

# %% [markdown]
### identification of surrogate variables

# %% [code]
sv <- DaMiR.SV(data_filt)

# %% [markdown]
### cleaning expression data

# %% [code]
data_adjust <- DaMiR.SVadjust(data_filt, sv, n.sv=9)

# %% [markdown]
# Exploring Data

# %% [code]
#DaMiR.Allplot(data=data_filt, df=colData(data_filt))
#graph2pdf(file = "bf.pdf")

# %% [code]
DaMiR.Allplot(data_adjust, colData(data_adjust))

# %% [markdown]
# Feature Selection

# %% [markdown]
### variable selection in Partial Least Squares (PLS)

# %% [code]
set.seed(12345)
data_clean <- DaMiR.transpose(assay(data_adjust))
df <- colData(data_adjust)
data_reduced <- DaMiR.FSelect(data_clean, df, th.corr=0.4)

# %% [markdown]
### removing highly correlated features

# %% [code]
data_reduced <- DaMiR.FReduct(data_reduced$data)

# %% [code]
DaMiR.MDSplot(data_reduced, df)

# %% [markdown]
### ranking and selecting most relevant features

# %% [code]
options(repr.plot.width = 8, repr.plot.height = 8, repr.plot.res = 200)
df.importance <- DaMiR.FSort(data_reduced, df)
#graph2pdf(file = "leftPanel.pdf")

# %% [code]
outputfile <- "dfimportance_hftxn_lgcpm.txt"
write.table(df.importance, file = outputfile, quote=FALSE, sep = "\t")

# %% [code]
# select best predictors:
selected_features <- DaMiR.FBest(
data_reduced, ranking=df.importance, n.pred = 13)

# %% [code]
selected_features$predictors

# %% [code]
options(repr.plot.width = 10, repr.plot.height = 8, repr.plot.res = 200)
DaMiR.Clustplot(selected_features$data, df)
graph2pdf(file = "rightPanel.pdf", width = 10, height = 8)

# %% [code]
```
