## Supplementary material for "TXN, a Xanthohumol Derivative, Attenuates High-Fat Diet Induced Hepatic Steatosis by Antagonizing PPARγ": Figure 10 source data: fig10.ipynb

```
# %% [code]
suppressPackageStartupMessages(library(ggplot2))
suppressPackageStartupMessages(library(emmeans))
suppressPackageStartupMessages(library(tidyverse))
suppressPackageStartupMessages(library(export))
suppressPackageStartupMessages(library(patchwork))
suppressPackageStartupMessages(library(ggpubr))
R.Version()

# %% [code]
theme_legend_free = function(){
	theme(axis.title.y = element_text(size = 15, 
	margin = margin(t=0, r=5, b=0, l=10)),
	axis.text.y = element_text(size = 15, face = 'bold'),
	axis.text.x = element_text(size = 15, face= 'bold'),
	axis.ticks.x = element_blank(),
    plot.title = element_text(size=13),
	legend.position = "none",
	plot.margin = unit(c(0.5,0.5,0,0.5), "cm"),
	panel.grid = element_blank(),
	panel.grid.major = element_blank(),
	panel.grid.minor = element_blank())
}

italic.text <- element_text(face = "italic")
theme_legend = function(){
	theme(axis.title.y = element_text(size = 14,
	margin = margin(t=0, r=5, b=0, l=10)),
	axis.text.y = element_text(size = 15, face = 'bold'),
	axis.text.x = element_text(size = 15, face= 'bold'),
	axis.title.x = element_text(size = 14),
    axis.title = italic.text,
	axis.ticks.x = element_blank(),
    plot.title = element_text(size=13),
	legend.position = "bottom",
	plot.margin = unit(c(0.5,0.5,0,0.5), "cm"),
	panel.grid = element_blank(),
	panel.grid.major = element_blank(),
	panel.grid.minor = element_blank())
}

# %% [markdown]
# Liver tissue

# %% [code]
data_liver <- read.csv('fig10_table.csv')
data_liver$treatment <- as.factor(data_liver$treatment)
colors <- c("#0080ff", "darkgreen")
head(data_liver)

# %% [code]
mycolor_alpha <- c("#8AB5F9", "#87A57E")

# %% [code]
pparg <- ggplot(data = data_liver, aes(x = treatment, y = pparg, fill = treatment)) +
    geom_hline(yintercept = 0, lty = 5) +
    stat_boxplot(geom = 'errorbar', linetype = 1, width = 0.1) +
    geom_boxplot(width = 0.3)+
	scale_fill_manual(values = mycolor_alpha, guide = FALSE) +
	xlab(" ") + ylab("log2FoldChange") +
	ggtitle("Ppar\u03B3") +
	theme_bw() +
	theme_legend_free()

label.df <- data.frame(treatment = c("HFD+TXN"),
                       pparg = c(-0.3))
pparg <- pparg + geom_text(data = label.df, label = c("***"), color='red', size = 12)

# %% [code]
cid <- ggplot(data = data_liver, aes(x = treatment, y = cidec, fill = treatment)) +
    geom_hline(yintercept = 0, lty = 5) +
    stat_boxplot(geom = 'errorbar', linetype = 1, width = 0.1) +
    geom_boxplot(width = 0.3)+
	scale_fill_manual(values = mycolor_alpha, guide = FALSE) +
	xlab(" ") + ylab("log2FoldChange") +
	ggtitle("Cidec") +
	theme_bw() +
	theme_legend_free()

label.df <- data.frame(treatment = c("HFD+TXN"),
                       cidec = c(-2))
cid <- cid + geom_text(data = label.df, label = c("***"), color='red', size = 12)

# %% [code]
plin <- ggplot(data = data_liver, aes(x = treatment, y = plin4, fill = treatment)) +
    geom_hline(yintercept = 0, lty = 5) +
    stat_boxplot(geom = 'errorbar', linetype = 1, width = 0.1) +
    geom_boxplot(width = 0.3)+
	scale_fill_manual(values = mycolor_alpha, guide = FALSE) +
	xlab(" ") + ylab("log2FoldChange") +
	ggtitle("Plin4") +
	theme_bw() +
	theme_legend_free()

label.df <- data.frame(treatment = c("HFD+TXN"),
                       plin4 = c(-0.8))
plin <- plin + geom_text(data = label.df, label = c("***"), color='red', size = 12)

# %% [code]
mog <- ggplot(data = data_liver, aes(x = treatment, y = mogat1, fill = treatment)) +
    geom_hline(yintercept = 0, lty = 5) +
    stat_boxplot(geom = 'errorbar', linetype = 1, width = 0.1) +
    geom_boxplot(width = 0.3)+
	scale_fill_manual(values = mycolor_alpha, guide = FALSE) +
	xlab(" ") + ylab("log2FoldChange") +
	ggtitle('Mogat1') +
	theme_bw() +
	theme_legend_free()

label.df <- data.frame(treatment = c("HFD+TXN"),
                       mogat1 = c(-0.4))
mog <- mog + geom_text(data = label.df, label = c("***"), color='red', size = 12)

# %% [code]
scatter1 <- ggplot(data_liver, aes(x = pparg, y = cidec)) + 
    geom_rug(aes(color = treatment)) +
    geom_point(aes(color = treatment, size = lv_pct), alpha=0.5) +
    scale_size(range = c(1, 14), guide=guide_legend(title="Liver mass:BW (%)")) +
    stat_smooth(method = "lm", formula = y ~ x, col = "black", se = FALSE, linetype = "dashed", size = 0.5)+
    stat_cor(label.x = -0.7, label.y = -4.85, r.digits = 2, p.digits = 2, color = 'red') +
    #stat_regline_equation(label.x = -1.3, label.y = 0) +
    scale_color_manual(name = "Diet", values = colors, guide = FALSE)+
    labs(x = 'Ppar\u03B3', y = 'Cidec') +
	theme_bw() + theme_legend()

# %% [code]
scatter3 <- ggplot(data_liver, aes(x = pparg, y = plin4)) + 
    geom_rug(aes(color = treatment)) +
    geom_point(aes(color = treatment, size = lv_pct), alpha=0.5) +
    scale_size(range = c(1, 14), guide=guide_legend(title="Liver mass:BW (%)")) +
    stat_smooth(method = "lm", formula = y ~ x, col = "black", se = FALSE, linetype = "dashed", size = 0.5)+
    stat_cor(label.x = -0.7, label.y = -2.8, r.digits = 2, p.digits = 2, color = 'red') +
    scale_color_manual(name = "Diet", values = colors, guide = FALSE)+
    labs(x = 'Ppar\u03B3', y = 'Plin4') +
	theme_bw() + theme_legend()

# %% [code]
scatter4 <- ggplot(data_liver, aes(x = pparg, y = mogat1)) + 
    geom_rug(aes(color = treatment)) +
    geom_point(aes(color = treatment, size = lv_pct), alpha=0.5) +
    scale_size(range = c(1, 14), guide=guide_legend(title="Liver mass:BW (%)")) +
    stat_smooth(method = "lm", formula = y ~ x, col = "black", se = FALSE, linetype = "dashed", size = 0.5)+
    stat_cor(label.x = -0.7, label.y = -3.3, r.digits = 2, p.digits = 2, color = 'red') +
    scale_color_manual(name = "Diet", values = colors, guide = FALSE)+
    labs(x = 'Ppar\u03B3', y = expression(italic('Mogat1'))) +
	theme_bw() + theme_legend()

# %% [code]
scatter5 <- ggplot(data_liver, aes(x = mogat1, y = cidec)) + 
    geom_rug(aes(color = treatment)) +
    geom_point(aes(color = treatment, size = lv_pct), alpha=0.5) +
    scale_size(range = c(1, 14), guide=guide_legend(title="Liver mass:BW (%)")) +
    stat_smooth(method = "lm", formula = y ~ x, col = "black", se = FALSE, linetype = "dashed", size = 0.5)+
    stat_cor(label.x = -1.7, label.y = -4.5, r.digits = 2, p.digits = 2, color = 'red') +
    scale_color_manual(name = "Diet", values = colors, guide = FALSE)+
    labs(x = 'Mogat1', y = 'Cidec') +
	theme_bw() + theme_legend()

# %% [code]
plotA <- (pparg | cid | plin | mog)& theme(legend.position = 'none')
plotB <- (scatter5 | scatter1 | scatter3 | scatter4)& theme(legend.position = 'bottom')
pcr <- (plotA / plotB) + plot_layout(guides = "collect")& theme(legend.position = 'bottom')

# %% [code]
#graph2pdf(pcr, file = "figure10.pdf", width = 16, height = 10)
```
