## Supplementary material for "TXN, a Xanthohumol Derivative, Attenuates High-Fat Diet Induced Hepatic Steatosis by Antagonizing PPARγ": Figure 12 source data: fig12.pdf

A1: 1 µM Rosiglitazone

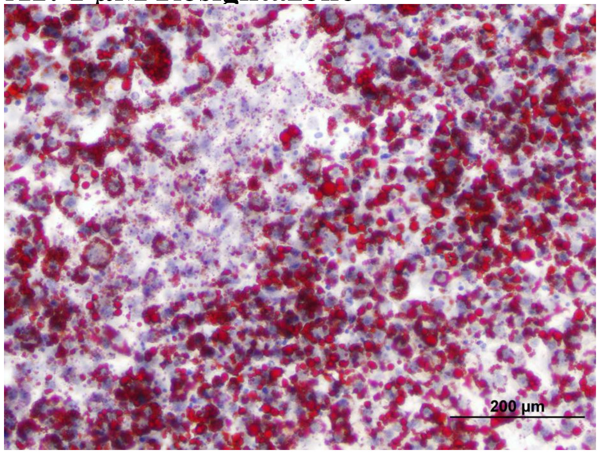


A2: 1 µM GW 9662

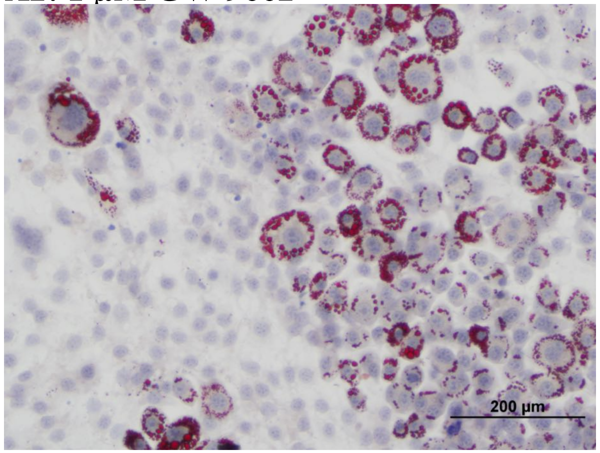


A3: 1 µM Rosiglitazone + 1 µM GW 9662

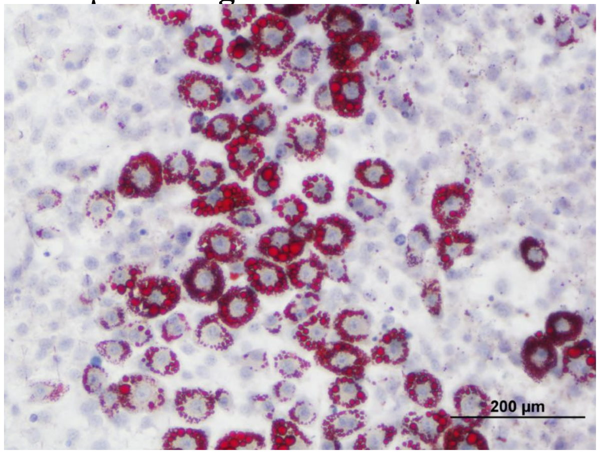


B1: 1 µM Rosiglitazone + 5 µM XN

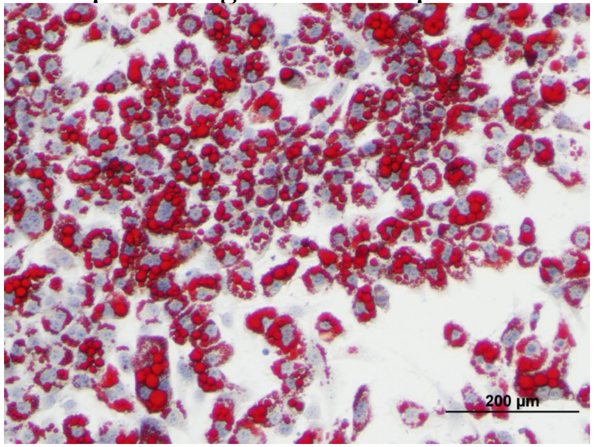


B2: 1 µM Rosiglitazone + 10 µM XN

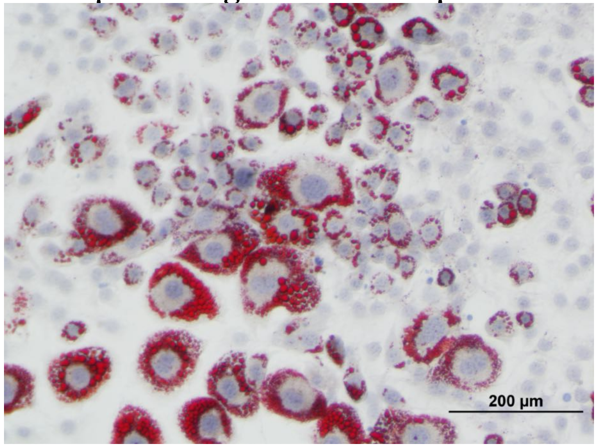


B3: 1 µM Rosiglitazone + 25 µM XN

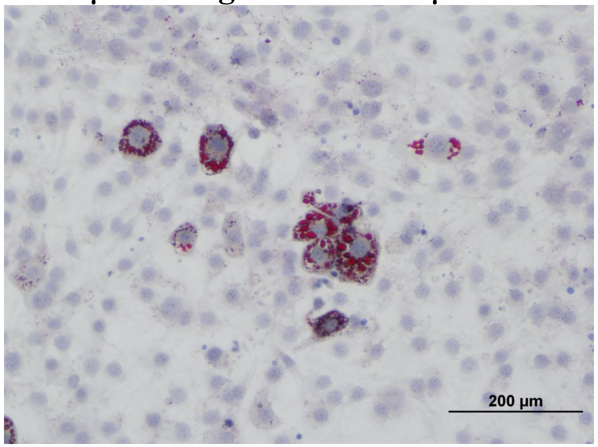


C1: 1 µM Rosiglitazone + 5 µM TXN

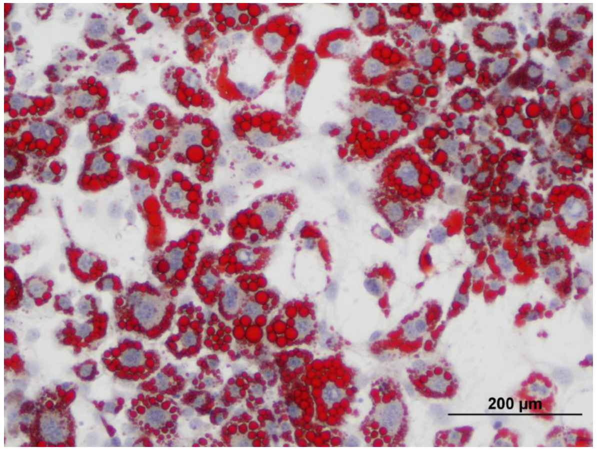


C2: 1 µM Rosiglitazone + 10 µM TXN

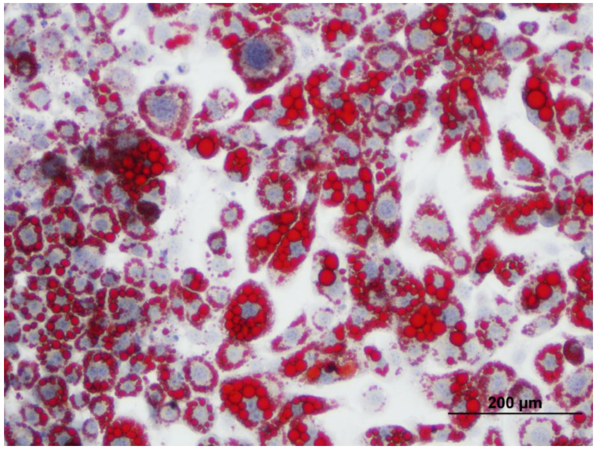


C3: 1 µM Rosiglitazone + 25 µM TXN

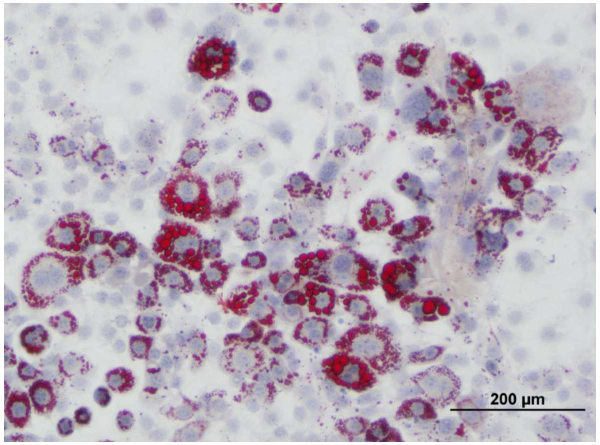
